# A multi-tissue atlas of genetic regulatory effects in sheep

**DOI:** 10.1101/2025.11.03.686346

**Authors:** Mian Gong, Ziyang Zhuang, Xiangrong Sun, Yuan Xu, Huanhuan Zhang, Yanan Wang, Dailu Guan, Ran Li, Xiaoning Lu, Zhonghao Bai, Pingjie Feng, Meiwen Song, Min Tian, Jingsheng Lu, Mingshan Wang, Xuemei Lu, Dongdong Wu, Peng Su, Peiyao Liu, Guoqing Zhang, Jianxin Shi, Mingzhu Shan, Yuanyuan Zhang, Zhu Meng, Hao Li, Xiaoyun He, Jianqi Yang, Yize Song, Xinyue Li, Xiaolong Du, Xiaoxu Zhang, Hao Yang, Jinyan Teng, Houcheng Li, Xiaoning Zhu, Huicong Zhang, Qing Lin, Di Zhu, Bingjin Lin, Xinfeng Liu, Jianquan Liu, Weijie Zheng, Wentao Gong, Bingxing An, Qi Zhang, Goutam Sahana, Mogens Sandø Lund, Cong Li, Jiazhong Guo, Xihong Wang, Yuwen Liu, Bingru Zhao, Xiaolei Yao, Yanli Zhang, Feng Wang, Wenxin Zheng, Juncheng Huang, Sen Wang, Jiang Di, Hanikezi Tulafu, Zhihong Liu, Shaoyin Fu, Yongbin Liu, Zijun Zhang, Yongju Zhao, Yinghui Lin, Jianning He, Jinshan Zhao, Hengbo Shi, Zhengguang Wang, Bingjie Li, Ruidong Xiang, Amanda J. Chamberlain, Weimin Wang, Qiuyue Liu, Jiyuan Li, Fenghua Lv, Ze Yan, Qien Yang, Guiping Zhao, Lin Jiang, Xianyong Lan, Huaijun Zhou, Richard P. M. A. Crooijmans, Ole Madsen, David E. MacHugh, John F. O’Grady, Marcel Amills, Gwenola Tosser-Klopp, Emily L. Clark, Jianlin Han, Mingxing Chu, Weiwei Wu, Yu Jiang, Zhangyuan Pan, Lingzhao Fang

## Abstract

Characterizing the impact of genomic variants on genome function and ultimately complex traits in livestock is essential for the development of sustainable precision agriculture and comparative genomics. Here, as part of the Farm animal Genotype-Tissue Expression (FarmGTEx) project, we present the pilot phase of the SheepGTEx resource through analyzing 6,761 RNA-sequencing samples of 51 primary tissues in a multi-breed population of sheep. We identify millions of regulatory variants associated with seven types of molecular phenotypes, and fine-map 322,468 primary and 114,017 non-primary effects, revealing a high degree of regulatory allelic heterogeneity. We systematically characterize the pleiotropic effects of these variants on molecular phenotypes, assess their context-specific regulatory patterns across tissues, breeds, sexes, and developmental stages, as well as explore their evolutionary constraints across mammals. Finally, we demonstrate the substantial potential of the SheepGTEx resource (https://sheepgtex.farmgtex.org), by providing examples of regulatory mechanisms underpinning 34 complex traits, population divergence between European and Asian breeds, and adaptive evolution in sheep over the past ten millennia.

## Introduction

Sheep (*Ovis aries*) is one of the first livestock species that were domesticated in the Fertile Crescent over 10,000□years ago^1^. Subsequent global dispersal, artificial selection, climate adaptation, and introgression from wild relatives have given rise to thousands of breeds that thrive in a milieu of agro□ecological environments while providing wool, meat, milk and other by-products to human societies^2^. Beyond their essential roles in the global agri-food system, sheep also serve as versatile translational models for human biology. For example, they can shed light on cardiopulmonary, neurological, immunological, and reproductive conditions, as well as high-altitude adaptation and fetal development^3,4^.

To substantially unlock the potential of sheep as both an agricultural and a model organism, understanding the genetic and molecular architecture underpinning diverse complex phenotypes of economic, ecological, and medical importance in this species is paramount. However, while genome□wide association studies (GWAS) have identified thousands of variants associated with complex traits in a variety of farm animal species, including over 8,000 associations of 182 base traits in sheep curated in the Animal QTLdb^5^, these variants often reside in non-coding regions of the genome, which makes functional assignment challenging^6^. Nonetheless, a growing amount of evidence supports the hypothesis that such variants influence complex phenotypes through modulating gene expression in a context-specific manner, such as tissue, sex and developmental stages^7,8^. Population-based molecular quantitative trait loci (molQTL) mapping is a promising way to functionally characterize regulatory mutations by systematically associating natural sequence variants with gene expression, as demonstrated by the human Genotype-Tissue Expression (GTEx)^9^ and Farm animal GTEx (FarmGTEx) projects^10–14^. However, such efforts remain unexplored in sheep. To date, there has only been a single expression QTL (eQTL) mapping study in sheep that used RNA sequencing (RNA-seq) data from two tissues (liver and muscle) in 149 animals^15^.

To address this gap, we have initiated the SheepGTEx project, as part of the ongoing international FarmGTEx effort^10^. In this pilot phase, we integrated 6,761 bulk RNA-seq datasets, including 1,565 newly generated and 5,196 existing samples from publicly available datasets, representing 49 tissues and 2 cell types (hereafter referred to as “tissues”) from 2,816 animals worldwide (**Fig. 1a–c; Supplementary Table 1**). The mean sample size was 174 across tissues, ranging from 40 samples for rectum and oviduct, respectively, to 713 for muscle. We systematically associated 3,427,511 genomic variants with seven types of transcriptomic phenotypes, followed by conditional and fine-mapping analyses for detecting primary effects (i.e., the lead association signal) and non-primary effects (including secondary, tertiary and higher-order independent signals)^16,17^. We then assessed the context-specificity of these regulatory variants across tissues, populations, sexes, and developmental stages (i.e., prenatal, juvenile and adult), as well as exploring their evolutionary constraints across several mammals, including cattle, pigs and humans (**Fig. 1d**). Finally, through examining GWAS results of 34 complex traits (median *n* = 2,131), selection signatures between current European and Asian breeds, and 52 ancient DNA samples spanning around 10,000 years, we demonstrated the utility of this SheepGTEx resource for dissecting the genetic and molecular architecture underlying complex traits, application in selective breeding, and for understanding environmental adaptation, and domestication (**Fig. 1e**). All uniformly processed data, results, and computational codes are freely available through the SheepGTEx web portal (https://sheepgtex.farmgtex.org).

**Fig. 1.**
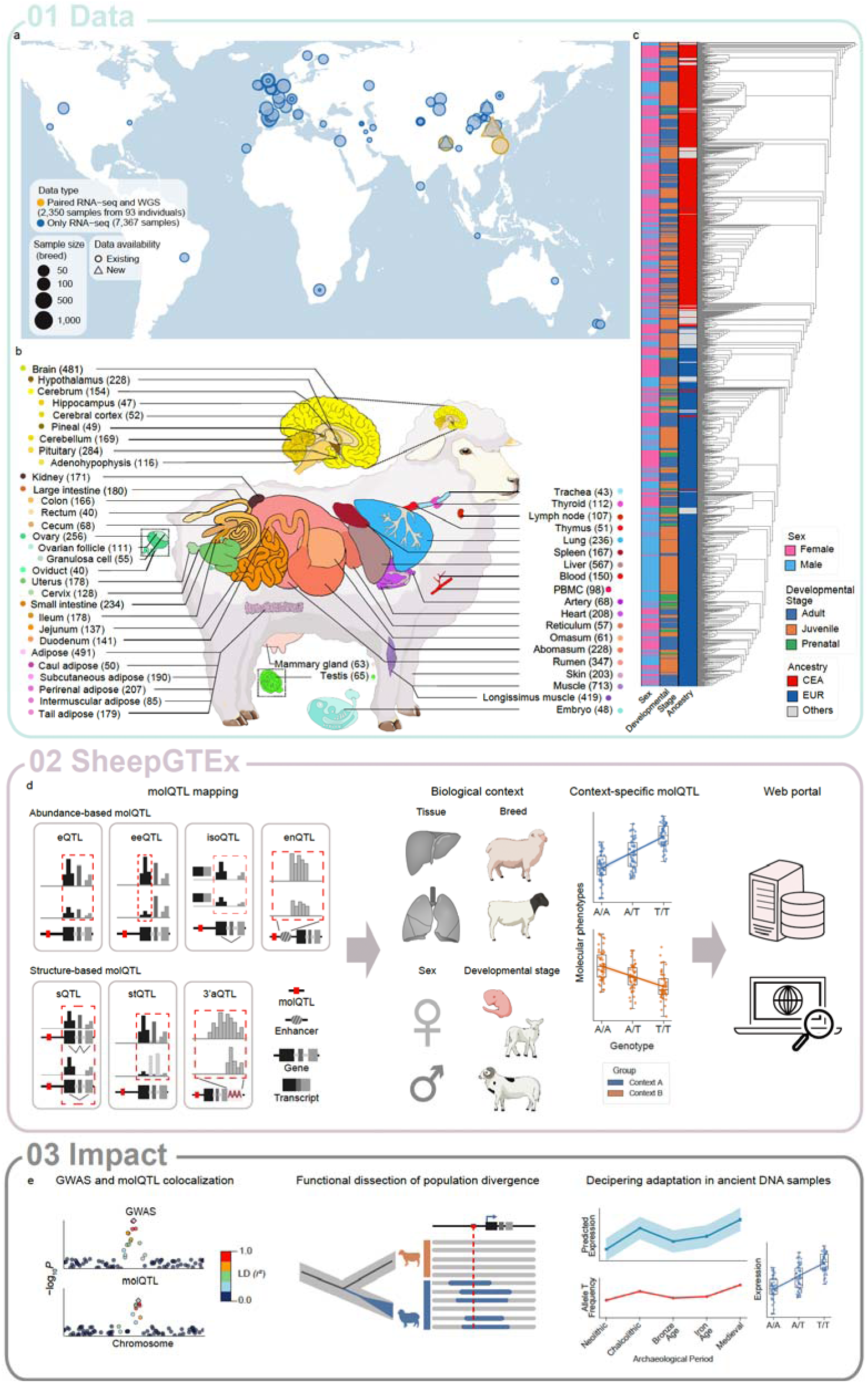
Overview of the pilot SheepGTEx project. **a**, Geographic origins of all 9,717 sheep RNA-seq samples that have been analysed in this study. **b**, Summary of 51 tissues profiled for molecular quantitative trait loci (molQTL) mapping in sheep. Tissue sample sizes (n□≥□40) are indicated in the respective parentheses, representing 2,816 animals. The colour key of tissue is consistently used across the entire manuscript. **c**, Neighbor-joining tree of all 2,816 animals based on 3,427,511 imputed variants, revealing two major ancestry groups (Central and East Asia -CEA and Europe -EUR), along with metadata on sex and developmental stage. **d**, The molQTL mapping and resource sharing in the pilot SheepGTEx project. We mapped seven types of steady molQTL and four types of context-specific molQTL. eQTL, expression QTL; eeQTL, exon expression QTL; isoQTL, isoform expression QTL; enQTL, enhancer expression QTL; sQTL, splicing QTL; stQTL, QTL for RNA stability of genes; 3’aQTL, QTL for 3’ UTR alternative polyadenylation (APA). All data are accessible at https://sheepgtex.farmgtex.org/, **e**, Applications of the SheepGTEx resource in deciphering genome-wide association studies (GWAS) results of complex traits, population divergence between current European and Asian breeds, and adaptive evolution of ancient samples across the past 10,000 years in sheep. LD, linkage disequilibrium.

## Results

### Overview of transcriptomic and genomic data in the SheepGTEx Project

For the transcriptomic data, we initially processed a total of 7,817 publicly available and 1,900 newly generated (1–42 tissues from 115 animals) RNA-seq samples, including 2,350 samples with matched whole-genome sequencing (WGS) data from 93 animals (**Fig. 1a; Supplementary Fig. 1a–f; Supplementary Tables 1 and 2**). After rigorous quality control (**Methods; Supplementary Fig. 1g–o; Supplementary Table 1**), 8,171 RNA-seq samples from 70 tissues were retained for quantifying seven different types of molecular phenotypes, including four abundance-based phenotypes [i.e., the expression levels of 25,699 protein-coding genes (PCGs) and long non-coding RNA genes (lncRNAs), 252,949 exons, 54,667 isoforms, and 81,004 enhancers], and three structure-based phenotypes (i.e., 1,611,287 alternative splicing, 12,466 RNA stability of genes^18^, and 13,842 3’UTR alternative polyadenylation- 3’UTR APA^19^) (**Fig. 1d**). The average Pearson’s correlation (*r*) between gene expressions and other molecular phenotypes of the same gene across samples within tissue was 0.23, ranging from 0.53 for exon expression to −0.05 for alternative splicing, providing evidence that these molecular phenotypes may represent distinct biological aspects of the transcriptome^9,20,21^ (**Supplementary Fig. 2a,b**).

Tissue type was the primary driver of variation in molecular phenotypes (**Extended Data Fig. 1a,b; Supplementary Figs. 2c and 3a,b; Supplementary Table 3**), and function enrichment of genes with tissue-specific expression [false discovery rate (FDR) < 0.05 and Log_2_FoldChange > 2 using the Wilcoxon rank-sum test^22^; see **Supplementary Note**] in Gene Ontology (GO) database, as expected, recapitulated the known tissue biology (**Extended Data Fig. 1c**). For example, genes with expression patterns confined to adipose (*n* = 458) and cerebrum (*n* = 2,238) tissues, were significantly enriched in *Fatty acid metabolic process* (FDR = 2.40 × 10^−10^) and *Regulation of trans-synaptic signaling* (FDR = 5.53 × 10^−107^), respectively (**Supplementary Table 4**). In addition, the tissue specificity of gene expression was evolutionarily conserved between sheep and three other mammals (i.e., cattle, pigs, and humans), with a slight decline corresponding to increasing evolutionary divergence (**Extended Data Fig. 1d; Supplementary Fig. 3c**). We further demonstrated the potential of this transcriptomic atlas in understanding the molecular basis of monogenic traits and for functional annotation of sheep genes (**Supplementary Figs. 4 and 5**). For example, 46 genes with causal effects on 39 Mendelian traits in sheep^23^ exhibited tissue-specific expression patterns, e.g., the *CLCN1* gene, with causal effects on myotonia, showed muscle-specific expression (**Supplementary Fig. 4**).

For RNA-seq samples without WGS data, we imputed their genotypes by using a multi-breed sheep genotype imputation reference panel comprising 3,125 existing WGS animals worldwide (**Supplementary Table 5**), which showed a matched population structure as RNA-seq samples (**Extended Data Fig. 2a**). After stringent quality control and comprehensive validation of genotype imputation in independent populations (**Supplementary Note**), we obtained a total of 3,427,511 common (minor allele frequency, MAF > 0.05) and high-quality (median imputation accuracy of 94.7%) genetic variants for all the 8,171 RNA-seq samples, with an average of 4,227 variants in the *cis*-window (±1 Mb from the transcription start site, TSS) for expressed genes from NCBI *Ovis aries* Annotation Release 104 (ARS-UI_Ramb_v2.0^24^, GCF_016772045.1) (**Extended Data Fig. 2b–o**). Furthermore, we predicted missing metadata of RNA-seq samples, including sex (inferred from chromosomal coverage and expression profiles), developmental stage (predicted via a Random Forest classifier applied to expression matrix), and breed (based on imputed genotypes), with a mean prediction accuracy of 97.0%, 84.5%, and 96.8%, respectively (**Supplementary Note; Supplementary Figs. 6–8; Supplementary Table 1**). We removed duplicated samples within each tissue based on both identity-by-state (IBS) ≥ 0.9 and genetic relationship ≥ 0.5, resulting in 6,761 RNA-seq samples representing 51 tissues (where *n* ≥ 40 animals, according to previous studies^11–13^) from 2,816 animals for subsequent molQTL mapping (**Fig. 1b,c; Supplementary Figs. 1o and 9; Supplementary Table 6**). Among these samples, missing information for ancestry (breed), sex, and developmental stage were imputed for 639 (9.5%), 854 (12.6%) and 321 (4.7%) samples, respectively (**Supplementary Fig. 10**).

### molQTL mapping and validation

After accounting for potential confounding factors (**Methods; Supplementary Fig. 11**), the estimated *cis*-heritability (*cis*-*h*^2^) of seven molecular phenotypes ranged from 0.073 to 0.135, which were lower than their *trans*-*h*^2^ (SNPs in all other chromosomes), which ranged from 0.313 to 0.521 (**Supplementary Figs. 12–14**). We subsequently conducted a *cis*-molQTL mapping analysis via TensorQTL^25^ in each one of the 51 tissues. From this, we identified a total of 19,213 (82.3% of all tested genes) eGenes for gene expression, 12,224 (65.9%) eeGenes for exon expression, 9,587 (85.6%) isoGenes for isoform expression, 3,904 (39.2%) enGenes for enhancer expression, 9,212 (54.0%) sGenes for alternative splicing, 10,345 (86.0%) stGenes for RNA stability, and 4,231 (35.9%) 3’aGenes for 3’UTR APA, which were collectively referred to as molGenes (FDR < 0.05; **Fig. 2a and Supplementary Table 7**). The associated variants were then termed eQTL, eeQTL, isoQTL, enQTL, sQTL, stQTL, and 3’aQTL, respectively. Of these molGenes, approximately 54.6% were regulated by more than one type of molQTL (**Supplementary Fig. 15a,b**). Genes associated with more molQTL exhibited higher expression levels, possessed a larger *cis*-*h*^2^ in gene expression, lower tissue specificity of gene expression (measured by TAU score^26^), weaker evolutionary constraint, and reduced network connectivity (measured by absolute eigengene-based connectivity, |kME|, in the WGCNA analysis^27^) (**Fig. 2b–e; Supplementary Figs. 15c, and 16).**

**Fig. 2.**
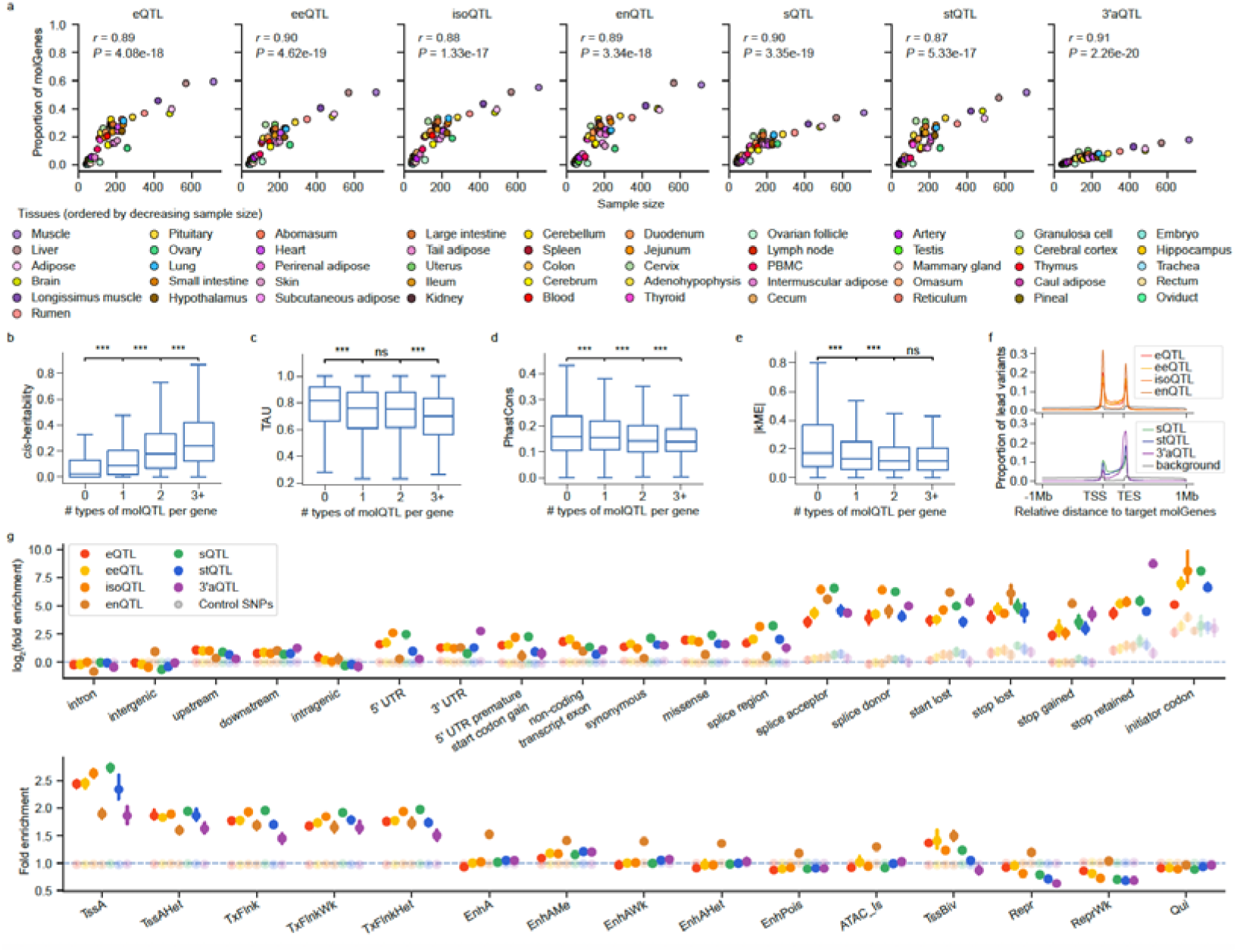
Regulatory architecture of seven molecular phenotypes across 51 tissues. **a**, Pearson’s correlation (*r*) between the proportion of molGenes (i.e., genes with significant molecular quantitative trait loci (molQTL) / all tested genes) and sample size across 51 tissues. *P* values were calculated using a two-sided Student’s *t*-test. eQTL, expression QTL; eeQTL, exon expression QTL; isoQTL, isoform expression QTL; enQTL, enhancer expression QTL; sQTL, splicing QTL; stQTL, QTL for RNA stability of genes; 3’aQTL, QTL for 3’ UTR alternative polyadenylation (APA). **b–e**, Boxplots showing differences in *cis*-heritability (**b**), tissue specificity (measured by TAU score in tspex^26^) (**c**), and DNA sequence conservation (PhastCons score of 100 vertebrate genomes) (**d**), and gene network connectivity (measured by absolute eigengene-based connectivity, |kME|, in WGCNA^27^) (**e**), across genes stratified by the number of molQTL types detected per gene. Statistical significance was assessed using the two-sided Mann-Whitney U test. ns: non-significant; *: *P* < 0.05; **: *P* < 0.01; ***: *P* < 0.001. **f**, Distribution of lead variants of molGenes relative to transcription start sites (TSS), and transcription end sites (TES). **g**, Enrichment (mean and 95% confidence interval) of seven types of lead *cis*-molQTL in 19 sequence features (top) and 15 chromatin states (bottom). Point, mean; error bar, 95% confidence interval. Lighter-colored points denote control SNPs matched for minor allele frequency (MAF) and linkage disequilibrium (LD) score. Chromatin state abbreviations: TssA, active promoter; TssAHet, flanking active TSS without ATAC; TxFlnk, transcribed gene; TxFlnkWk, weakly transcribed gene; TxFlnkHet, transcribed without ATAC; EnhA, strong active enhancer; EnhAMe, medium enhancer with ATAC; EnhAWk, weak enhancer; EnhAHet, active enhancer without ATAC (heterochromatic); EnhPois, poised enhancer; ATAC_Is, ATAC island; TssBiv, bivalent/poised TSS; Repr, polycomb-repressed; ReprWk, weakly repressed polycomb; Qui, quiescent.

To assess the reliability of the molQTL identified in the current work, we conducted a series of validations, which encompassed the use of linear mixed models, internal validation, external/independent validation, allele-specific expression (ASE), functional enrichment (to provide biological evidence), and cross-species comparison (to offer evolutionary evidence). All of these approaches demonstrated a high reliability of characterized molQTL (**Supplementary Figs. 17–19**). For instance, the internal validation across 33 tissues with sample sizes > 100 revealed an average replication rate of 0.90 and an effect size correlation of 0.92 (**Supplementary Fig. 18a**). The effect sizes derived from ASE were significantly correlated (mean Spearman’s ρ = 0.72) with those of eQTL (**Supplementary Fig. 18b–d**). The abundance-based molQTL (i.e., eQTL, eeQTL, isoQTL, and enQTL) showed a higher enrichment near TSS in comparison to the transcription end site (TES), whereas the opposite pattern was observed for structure-based molQTL (sQTL, stQTL, and 3’aQTL) (**Fig. 2f**). All types of molQTL were significantly enriched in regulatory elements (e.g., 5’UTR, 3’UTR, and promoters) and splice-related regions (e.g., splice acceptor and donor) (**Fig. 2g**). More specifically, among all molQTL, isoQTL and sQTL displayed the highest enrichment in splice-related regions, 3’aQTL in stop retained codon, and enQTL in enhancer-like chromatin states, recapitulating the known biology of these variants (**Fig. 2g**). Furthermore, both *cis*-*h*^2^ of orthologous gene expression and effect sizes of ancestral alleles were significantly correlated between sheep and three other mammalian species (cattle, pigs, and humans), while the lead eQTL in sheep resided closer to TSS (**Extended Data Fig. 3; Supplementary Fig. 19**).

To assess the impact of genotype imputation on the eQTL detection, we systematically compared *cis*-eQTL results derived from RNA-imputed genotypes with those from WGS-called genotypes across 21 tissues in 93 animals with matched RNA-seq and WGS data (**Extended Data Fig. 4; Supplementary Table 2**). In general, these two approaches exhibited similar statistical power for eGene discovery (**Extended Data Fig. 4b–d**). Lead variants of shared eGenes exhibited larger effect sizes than those of non-shared genes (**Extended Data Fig. 4e**). Lead and fine-mapped variants identified by both methods were often different but in high correlation of eQTL effect sizes (**Extended Data Fig. 4f–i**). Moreover,87.4% of lead and fine-mapped eQTL for the same eGene were in high linkage disequilibrium (LD; *r*^2^ > 0.5) between both approaches (**Extended Data Fig. 4j,k**), reflecting the same underlying causal variants. While both datasets showed significant enrichment in active regulatory regions, WGS-called genotypes consistently exhibited stronger enrichment signals than RNA-imputed genotypes (**Extended Data Fig. 4l**), indicative of improved fine-mapping resolution. Next, we explored whether the imputation of gene expression in unmeasured tissues of these animals could improve eQTL mapping^28^ (**Supplementary Note; Supplementary Fig. 20**), as many animals have RNA-seq data derived from a small number of tissues, particularly for datasets in the public domain. The statistical power to discover eGenes using imputed expression data was higher than that based on raw expression data, particularly in tissues with small sample sizes (*n* < 100) (**Extended Data Fig. 5a–h; Supplementary Table 8**). Compared to existing eQTL, the newly detected eQTL showed smaller effect sizes, lower MAF, and slightly higher enrichment in enhancers (**Extended Data Fig. 5i–l**). Beyond *cis*-eQTL mapping, we conducted an exploratory analysis for *trans*-eQTL in 15 tissues with *n* ≥ 200 animals (**Supplementary Note; Supplementary Fig. 21; Supplementary Table 9**). On average, we detected 222 *trans*-eGenes across tissues (**Supplementary Fig. 21b**) and observed 79 genomic hotspots of lead *trans*-eQTL that affected multiple *trans*-eGenes across the genome (**Supplementary Fig. 21c**).

The 135 genes located within hotspots were significantly enriched in fundamental biological processes like *Ribosomal subunit* (FDR = 1.13 × 10^−4^) and *Cytoplasmic translation* (FDR = 4.98 × 10^−4^) GO terms (**Supplementary Table 10**). Furthermore, we observed significant enrichment of lead *trans*-eQTL among *cis*-eQTL within the same tissue (**Supplementary Fig. 21d**), aligning with established models where distal regulatory signals often originate from local genetic effects^9^. Among 3,325 detected *trans*-eQTL, an average of 48.8% were significantly mediated by a *cis*-eQTL (**Supplementary Fig. 21e**). This confirms that nearly half of the detected *trans*-regulatory signals are likely to be driven by the propagation of local *cis*-regulatory effects. Of the 818 *cis*-mediating eGenes that colocalized with *trans*-eGenes (posterior probability of colocalization, PP_H4_ > 0.8 and average mappability of *trans*-eGenes > 0.8), 121 showed multiple colocalized *trans*-eQTL (**Supplementary Fig. 21f,g**), and 20 encoded transcription factors (TFs) annotated in AnimalTFDB (v4.0)^29^ (**Supplementary Table 11**).

### Shared and specific regulatory mechanisms underlying seven types of molQTL

To investigate whether certain genomic variants regulate multiple molecular phenotypes associated with the same gene, we performed colocalization analyses between eQTL and each of the six molQTL types. Overall, 71.5% of eeQTL, 58.7% of isoQTL, 32.3% of enQTL, 30.9% of sQTL, 39.6% of stQTL and 36.5% of 3’aQTL colocalized with an eQTL of the same gene (posterior probability for shared causality, PP_H4_ > 0.5) (**Fig. 3a**). The LD between lead eQTL and lead variants of other molQTL were generally low (median *r*^2^ = 0.23), particularly for structure-based molQTL (median *r*^2^ = 0.14) (**Supplementary Fig. 22a**), suggesting that distinct causal variants often regulate different molecular phenotypes for the same gene. As shown in **Fig. 3b**, several genes exhibited independent regulatory signals between molecular phenotypes, including *ARIH2* in longissimus muscle (eQTL vs. isoQTL), *C21H11orf86* in small intestine (eQTL vs. enQTL), *STXBP1* in brain (eQTL vs. sQTL), and *SAMD14* in pituitary (eQTL vs. 3’aQTL). To further explore the pleiotropic effects of regulatory variants on different molecular phenotypes, we classified lead eQTL based on the number of additional molecular phenotypes they influenced within the same genes. Approximately 59.0% of all eQTL exhibited regulatory effects on other molecular phenotypes (**Fig. 3c; Supplementary Fig. 22b,c**). The eQTL affecting the broader range of molecular phenotypes tended to show larger effect sizes on gene expression, higher MAF, shorter distance to the TSS, and higher enrichment in regulatory regions (**Fig. 3d–g; Supplementary Fig. 22d**).

**Fig. 3.**
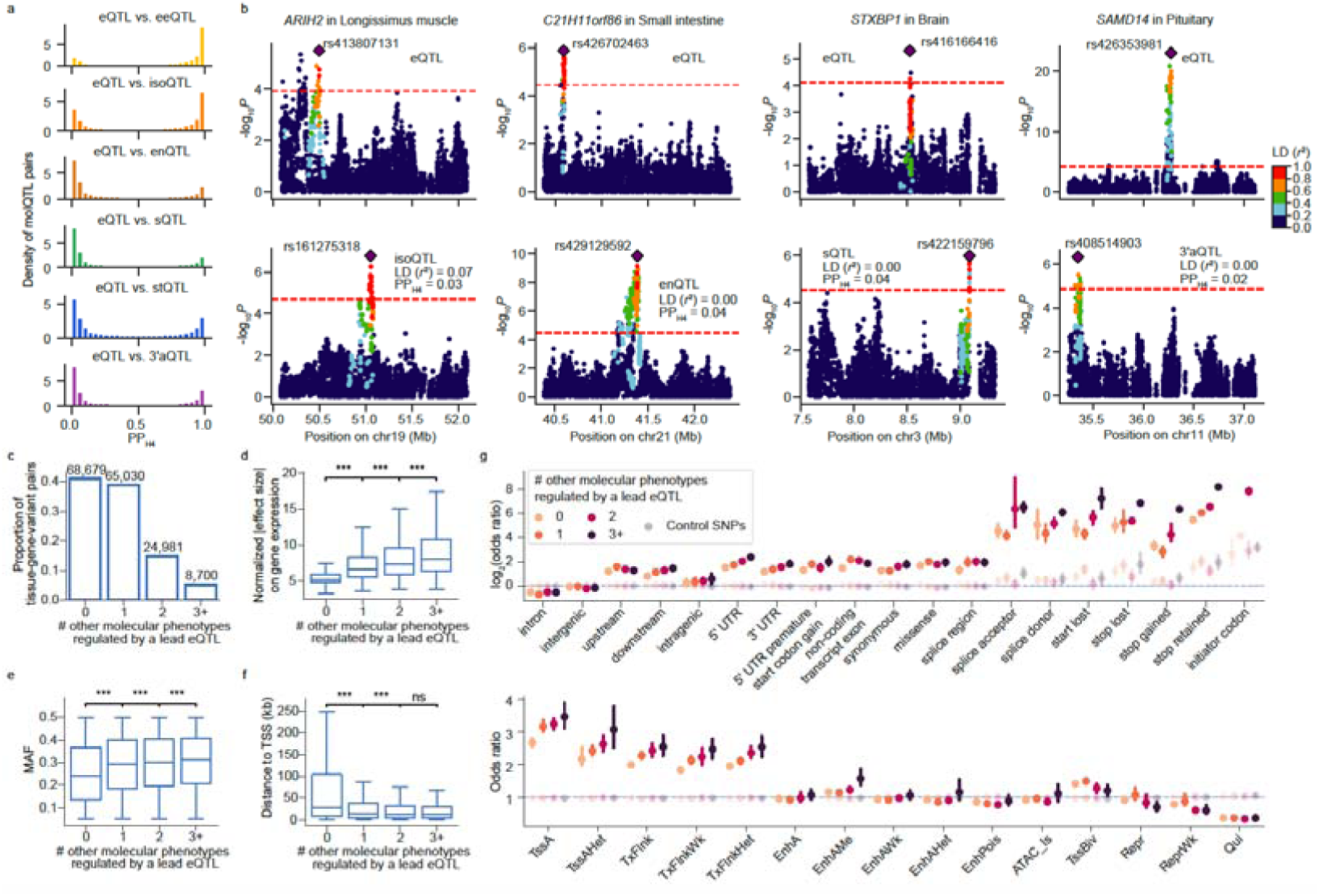
Distinct regulatory mechanisms underlying expression quantitative trait loci (eQTL) and six other types of molecular QTL (molQTL). **a**, Colocalization between eQTL and the other six types of molQTL within the same gene, inferred from PP_H4_ (posterior probability that two phenotypes share the same causal variant). **b**, Examples of genes with distinct molQTL types (PP_H4_ < 0.5). *r*^2^ indicates linkage disequilibrium (LD) between lead variants of eQTL and those of other molQTL. **c**, Proportion of lead eQTL with regulatory effects on other molecular phenotypes within the same gene. Numbers above bars denote lead eQTL affecting 0, 1, 2, or ≥ 3 additional molecular phenotypes. **d–f**, Boxplots comparing normalized |effect size| of respective eQTL (**d**), minor allele frequency (MAF) (**e**), and distance to transcription start site (TSS) (**f**), among lead eQTL with different numbers of pleiotropic regulatory effects. *P* values were calculated using the two-sided Mann-Whitney U test. ns: non-significant; *: *P* < 0.05; **: *P* < 0.01; ***: *P* < 0.001. **g**, Enrichment (mean ±95% confidence interval) of eQTL with varying degrees of molecular pleiotropy in 19 sequence features (top) and 15 chromatin states (bottom). Lighter-colored points denote control SNPs matched for MAF and LD score. Chromatin state abbreviations: TssA, active promoter; TssAHet, flanking active TSS without ATAC; TxFlnk, transcribed gene; TxFlnkWk, weakly transcribed gene; TxFlnkHet, transcribed without ATAC; EnhA, strong active enhancer; EnhAMe, medium enhancer with ATAC; EnhAWk, weak enhancer; EnhAHet, active enhancer without ATAC (heterochromatic); EnhPois, poised enhancer; ATAC_Is, ATAC island; TssBiv, bivalent/poised TSS; Repr, polycomb-repressed; ReprWk, weakly repressed polycomb; Qui, quiescent.

### Fine mapping revealed distinct features of primary and non-primary effects

To investigate the allelic heterogeneity of molQTL, we first performed conditional analysis using stepwise regression and detected, on average, 11.3% of eGenes, 13.5% of eeGenes, 16.7% of isoGenes, 10.4% of enGenes, 15.1% of sGenes, 8.1% of stGenes, and 5.9% of 3’aGenes harbored at least two independent signals across 51 tissues (**Fig. 4a; Supplementary Table 12**). To further pinpoint candidate causal variants, we conducted fine mapping analysis and identified at least one credible set (variants with cumulative PIP ≥ 95%) in 35.2% of eGenes, 19.5% of eeGenes, 21.1% of isoGenes, 28.8% of enGenes, 16.8% of sGenes, 34.8% of stGenes, and 38.6% of 3’aGenes (**Extended Data Fig. 6a; Supplementary Table 13**). The number of independent molQTL per gene was significantly correlated with the number of credible sets (**Extended Data Fig. 6b–h**). Among all molGenes, 7.0**–**12.3% exhibited multiple credible sets (**Extended Data Fig. 6a**), highlighting the complexity of genetic regulation of molecular phenotypes. Furthermore, 7.6**–**11.4% of credible sets contained a single variant, providing additional evidence that the current multi-breed SheepGTEx data facilitated high-resolution molQTL mapping analyses due to reduced LD (**Fig. 4b**). In addition, compared to lead variants, fine-mapped variants showed a higher enrichment in regulatory regions (**Extended Data Fig. 6i**). Genes with more independent signals tended to have a higher *cis*-*h*^2^, lower network connectivity, and lower evolutionary constraint at the sequence level, indicative of relaxed purifying selection^30^ (**Fig. 4c–f; Supplementary Fig. 23a–d**). Among molGenes with ≥ 4 independent signals, compared to primary variants, non-primary variants often showed smaller effect sizes, lower MAF, were further away from the TSS, and were more enriched in splice-related variants and repressed chromatin states (e.g., poised promoters and repressed Polycomb regions) but less in promoter-like elements (**Fig. 4g–j; Supplementary Fig. 23e–g**).

**Fig. 4.**
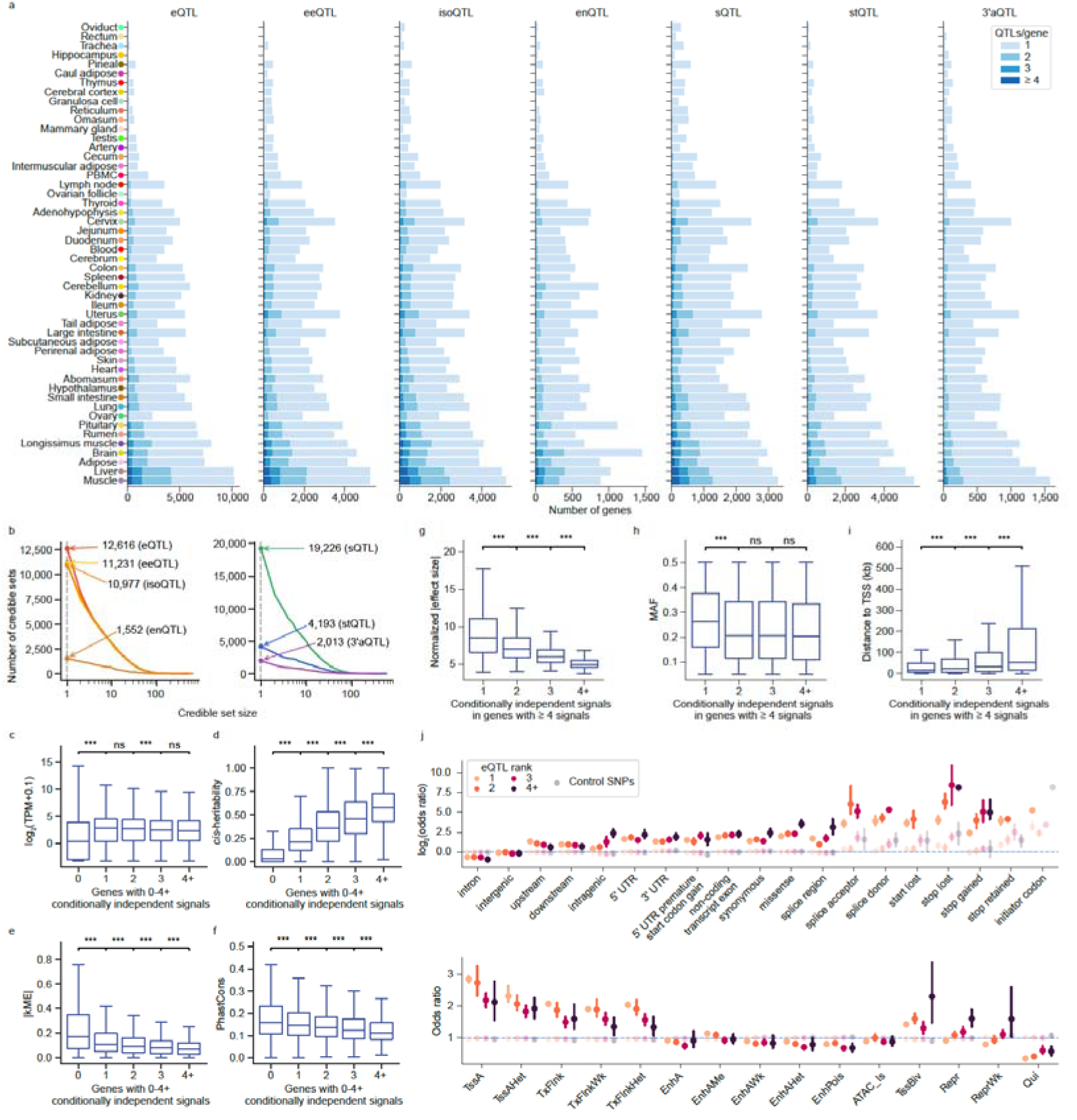
Main features of primary and non-primary molQTL. **a**, Number of molGenes harbouring one or more independent *cis*-molQTL signals across tissues. Tissues are ordered from smallest (top) to largest (bottom) sample size. **b**, The number of credible sets as a function of credible set size (i.e., the number of variants in the credible set) across seven types of molQTL. The values denote the number of credible sets with a single causal variant. **c–f**, Boxplots showing comparisons among genes stratified by the number of independent *cis*-eQTL signals per gene, including median gene expression levels (**c**), *cis*-heritability (**d**), gene network connectivity (absolute eigengene-based connectivity, |kME|) (**e**), and sequence conservation (PhastCons score) (**f**). **g–i**, Boxplots illustrating normalized |effect sizes| (**g**), minor allele frequency (MAF) (**h**), and distances to transcription start site (TSS) (**i**) for independent eQTL signals ranked by discovery order within each eGene. *P* values in **c-i** were calculated using the two-sided Mann-Whitney U test. ns: non-significant; *: *P* < 0.05; **: *P* < 0.01; ***: *P* < 0.001. **j**, Enrichment analysis (mean ±95% confidence interval) of eQTL signals detected by order of discovery across 19 sequence features (top) and 15 chromatin states (bottom). Lighter-colored points denote control SNPs matched for minor allele frequency (MAF) and linkage disequilibrium (LD) score. Chromatin state abbreviations: TssA, active promoter; TssAHet, flanking active TSS without ATAC; TxFlnk, transcribed gene; TxFlnkWk, weakly transcribed gene; TxFlnkHet, transcribed without ATAC; EnhA, strong active enhancer; EnhAMe, medium enhancer with ATAC; EnhAWk, weak enhancer; EnhAHet, active enhancer without ATAC (heterochromatic); EnhPois, poised enhancer; ATAC_Is, ATAC island; TssBiv, bivalent/poised TSS; Repr, polycomb-repressed; ReprWk, weakly repressed polycomb; Qui, quiescent.

### Tissue-specific regulatory effects

In general, tissues with similar biological functions (e.g., central nervous system and immune tissues) tended to cluster together with respect to both molecular phenotypes and their corresponding molQTL effects (**Fig. 5a,b; Supplementary Figs. 24–26**). Notably, this occurs despite the largely independent regulatory architectures operating across different molecular layers within a given tissue (**Supplementary Fig. 2a,b**), consistent with observations in humans^9,20,21^ and other livestock species^12,13,31^. In agreement with previous reports^9,11–13^, testis, embryonic tissues, and liver clustered separately from other primary tissues. Notably, the forestomaches of ruminants (i.e., rumen, reticulum, and omasum) exhibited distinct clustering from other digestive tissues such as the abomasum and intestines, underscoring their specialized cellular composition and biological roles in plant-based digestion^32,33^. The extent of tissue sharing for molQTL followed a U-shaped distribution (**Fig. 5c**), indicating that genetic regulatory effects tend to be either highly tissue-specific or ubiquitous^9,11–13^. The eGenes that were active in more tissues displayed higher network connectivity, lower tissue specificity of expression, but weaker evolutionary constraint (**Fig. 5d; Extended Data Fig. 7a,b**). Moreover, the respective eQTL exhibited smaller effect sizes, higher MAF, shorter distances to TSS, more enrichment in promoters, but less enrichment in enhancers (**Fig. 5e; Extended Data Fig. 7c–e**). Of note, non-primary eQTL were more broadly shared across tissues than primary eQTL (**Extended Data Fig. 7f**).

**Fig. 5.**
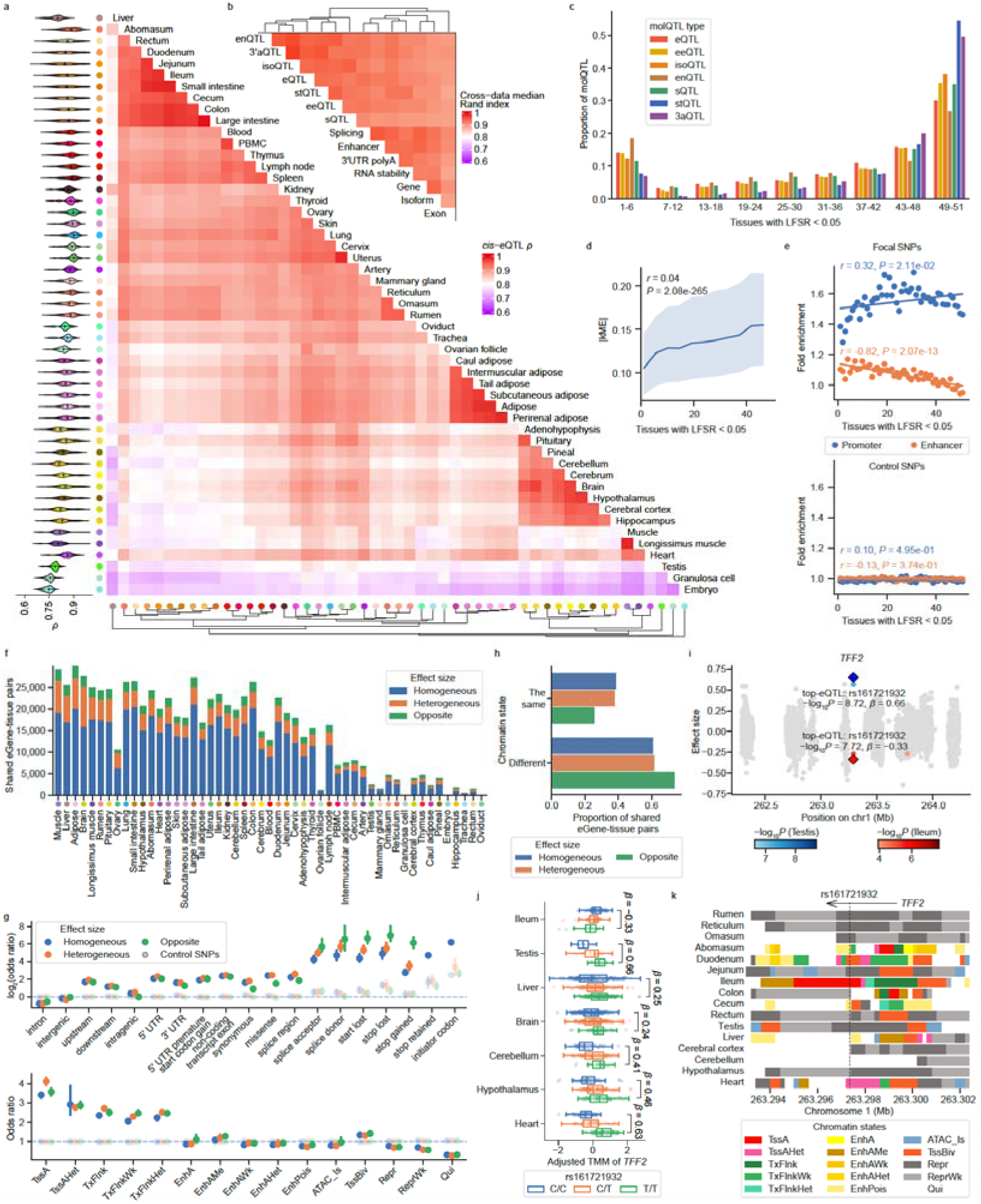
Tissue sharing and specificity of molQTL. **a**, Heatmap showing pairwise Spearman’s correlation (ρ) of *cis*-eQTL effect sizes across 51 tissues. Tissues were hierarchically clustered using the complete linkage method based on the maximum distance of ρ. Violin plots (left) display the distribution of Spearman’s ρ between each target tissue and all other tissues. **b**, Similarity of tissue clustering across seven molecular phenotypes and their respective *cis*-molQTL, measured by the median pairwise Rand index. **c**, Proportion of molQTL active across tissues, measured by the number of tissues with a *mashr* local false sign rate (LFSR, equivalent to FDR) <□0.05. **d**, Gene network connectivity (absolute eigengene-based connectivity, |kME|) as a function of the number of tissues sharing active eQTL. Solid lines indicate medians, and shading represents the interquartile range. The *P* value was assessed by Pearson’s correlation (*r*) test. **e**, Enrichment of *cis*-eQTL in promoter and enhancer regions, stratified by the number of tissues in which eQTL are active. Focal *cis*-eQTL SNPs are shown in the top panel, with control SNPs matched for minor allele frequency (MAF) and linkage disequilibrium (LD) score in the bottom panel. Linear regression models were fitted, and significance assessed by Pearson’s correlation with a two-sided Student’s *t*-test. **f**, Number of shared eGenes between each target tissue and all other tissues, stratified by concordance of lead variant effect sizes: homogeneous (overlapping 95% confidence intervals, CI), heterogeneous (non-overlapping 95% CI but the same direction of effect), and opposite (non-overlapping 95% CI with opposite directions of effect). **g**, Functional enrichment of lead *cis*-eQTL for the three categories of tissue-shared eGenes across 19 sequence features (top) and 15 chromatin states (bottom). Points indicate means; error bars represent 95% confidence intervals. Lighter-colored points denote control SNPs matched for MAF and LD score. **h**, Proportion of the three categories of tissue-shared eGenes whose lead *cis*-eQTL are located on the same or different chromatin states between tissues. **i**, *cis*-eQTL effects of *TFF2* in testis and ileum. Diamonds indicate the lead eQTL, while grey dots represent non-significant variants. **j**, Boxplots showing expression patterns of *TFF2* according to three genotypes of the lead eQTL (rs161721932) in testis and six additional tissues with significant associations. **k**, Chromatin states surrounding rs161721932 (denoted by the dashed line) of *TFF2* across ten digestive-related tissues and six additional tissues with significant signals. Chromatin state abbreviations: TssA, active promoter; TssAHet, flanking active TSS without ATAC; TxFlnk, transcribed gene; TxFlnkWk, weakly transcribed gene; TxFlnkHet, transcribed without ATAC; EnhA, strong active enhancer; EnhAMe, medium enhancer with ATAC; EnhAWk, weak enhancer; EnhAHet, active enhancer without ATAC (heterochromatic); EnhPois, poised enhancer; ATAC_Is, ATAC island; TssBiv, bivalent/poised TSS; Repr, polycomb-repressed; ReprWk, weakly repressed polycomb; Qui, quiescent.

To characterize the degree of heterogeneity of regulatory effects across tissues^12,34^, we classified eGene-tissue pairs into three categories based on the overlap of 95% confidence intervals (CI) of lead eQTL effect sizes: homogeneous (overlapping), heterogeneous (non-overlapping but with concordant direction of effect), and opposite (discordant direction of effect). On average, 71.8%, 19.4%, and 8.8% of eGenes exhibited homogeneous, heterogeneous, and opposite regulatory effects between tissues, respectively (**Fig. 5f**). eGenes exhibiting opposite effect directions between tissues were significantly enriched for variants located in splice-related regions (e.g., splice acceptor and donor sites), high-impact coding variants (e.g., start lost, stop gained, and stop lost), as well as chromatin states associated with active enhancers and weakly transcribed regions (**Fig. 5g**). Moreover, genes showing opposite regulatory effects were more likely to encode TFs (3.3%) than those with heterogeneous (2.5%) or homogeneous (1.9%) effects, based on sheep TF annotations from AnimalTFDB (v4.0)^29^, suggesting that these variants may act as “master switches” in divergent gene regulatory networks between tissues. Consistently, eGenes with opposite effects were also more likely to reside in different chromatin states between tissues compared to the other two categories (**Fig. 5h**). A compelling example is rs161721932, which modulates *TFF2* expression (**Fig. 5i**). The T allele is associated with decreased *TFF2* expression in the ileum but increased expression in other six tissues (**Fig. 5j**). Epigenomic annotations revealed an active promoter state specifically in the ileum that was absent in other tissues (**Fig. 5k**). Given the known role of *TFF2* in mucosal integrity and inflammatory bowel disease^35^, this antagonistic effect likely reflects tissue-specialized regulatory mechanisms tailored to ileal physiology. Additional examples included: rs412019428 showing opposite effects on *EXOSC1* expression between jejunum and heart, rs420177818 displaying opposite effects on *RGP1* expression between duodenum and perirenal adipose, and rs160450947 exhibiting heterogeneous effects on *POMC* expression between cerebellum and skin (**Extended Data Fig. 7g–o**).

### Context-specific molQTL shaped by ancestry, sex, and developmental stage

Population genetic analysis revealed that all the 2,816 animals in this phase of SheepGTEx were primarily derived from two major ancestry groups: Central and East Asia (CEA) and Europe (EUR) (**Fig. 1c; Supplementary Figs. 10a and 27**). We then systematically assessed how genetic background impacts the landscape of regulatory effects by comparing fine-mapped *cis*-eQTL between ancestry groups (**Fig. 6a**). In general, eQTL effects were shared between ancestries (mean π□ = 0.86; mean Pearson’s *r* = 0.82 for effect sizes) (**Extended Data Fig. 8a,b**). Compared to the single-ancestry population, the combined population could increase the eGene discovery by 136.4% and also improve fine-mapping resolution, due to the increased sample size and reduced LD (**Extended Data Fig. 8c–e**). We further identified ancestry-specific eQTL (anc-eQTL) following a previous framework^36^ and classified them into three categories: frequency-difference eQTL (fd-eQTL, driven by differences in MAF), LD-difference eQTL (ld-eQTL, due to differences in LD patterns), and heterogeneous-effect eQTL (he-eQTL, with significantly different effect sizes) (**Fig. 6a; Methods**). In total, we detected 2,327 ancestry-specific eGenes (anc-eGenes, genes harboring at least one anc-eQTL) across 14 tissues, ranging from 40 in skin to 419 in liver (**Fig. 6b; Supplementary Table 14**). Among these, fd-eQTL were the most prevalent, representing 1,141 anc-eGenes, such as rs405681252 of *ACAA1* in adipose (MAF_CEA_ = 0.175 and MAF_EUR_ = 0.005) (**Fig. 6c**). We also identified 954 anc-eGenes with ld-eQTL, such as *ABCB6* in liver, where independent lead SNPs were observed in EUR (rs159573658) and CEA (rs159573658) populations (*r*² = 0.067) (**Fig. 6d**). The discovery power of both fd-eQTL and ld-eQTL were correlated with sample size differences between ancestry groups (**Extended Data Fig. 8f,g**). Finally, we detected 505 anc-eGenes containing he-eQTL (from 8 in hypothalamus to 94 in pituitary), which may possibly reflect genotype × genotype or genotype × environment interactions. For instance, rs402321648 exhibited opposite regulatory effects on *RGS6* expression in liver between the EUR and CEA populations (**Fig. 6e**).

**Fig. 6.**
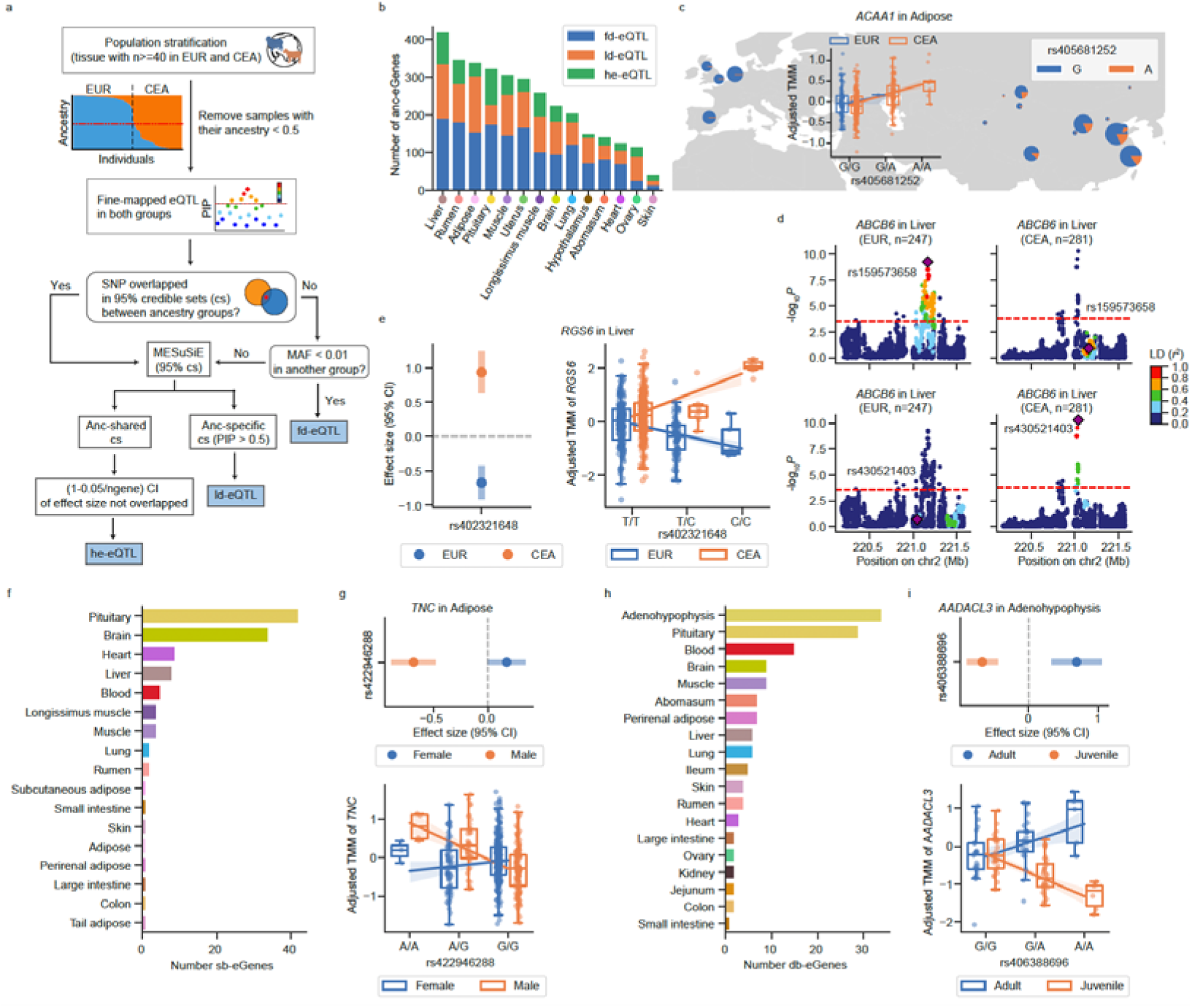
Context-specific molQTL. **a**, Workflow for the identification of ancestry-specific eQTL (anc-eQTL), which are classified into three categories: frequency-difference eQTL (fd-eQTL, driven by MAF differences between ancestry groups), LD-difference eQTL (ld-eQTL, due to linkage disequilibrium differences between ancestry groups identified via fine-mapping using MESuSiE (v1.0))^112^, and heterogeneous-effect eQTL (he-eQTL, shared between ancestry groups but exhibiting significantly different effect sizes, based on non-overlapping adjusted confidence intervals -CI). **b**, Number of ancestry-specific eGenes (anc-eGenes) identified across 14 tissues, stratified by anc-eQTL types. **c–e**, Examples of three types of anc-eQTL: fd-eQTL for *ACAA1* in adipose tissue (**c**), ld-eQTL for *ABCB6* in liver (**d**), and he-eQTL for *RGS6* in liver (**e**). Point plots show eQTL effect sizes with 95% CI; boxplots include fitted linear regression lines with 95% CI. Conventions are consistent across subsequent panels. **f**, Number of sex-biased eGenes (sb-eGenes) identified across 17 tissues. **g**, Example of sb-eQTL for *TNC* in adipose. **h**, Number of developmental stage-biased eGenes (db-eGenes) identified across 19 tissues. **i**, Example of db-eQTL for *AADACL3* in adenohypophysis.

Furthermore, we investigated the impact of sex and developmental stage (categorized as: prenatal; juvenile, ≤ 1 year; adult, > 1 year) on gene expression and genetic regulatory effects (**Supplementary Note; Supplementary Fig. 10b,c**). On average, 1,674 genes were differentially expressed between sexes across 46 tissues, and 3,760 genes were differentially expressed between developmental stages across 51 tissues, hereafter referred to as sex-biased and developmental stage-biased genes, respectively (**Supplementary Figs. 28–30**). The sex-biased genes were significantly enriched in pathways such as *Gamete generation* (*P*_adj_ = 2.10 × 10^−11^), *Sexual reproduction* (*P*_adj_ = 1.80 × 10^−12^), *Multicellular organismal reproductive* (*P*_adj_ = 9.36 × 10^−13^), and *Positive regulation of inflammatory response* (*P*_adj_ = 2.02 × 10^−6^) (**Supplementary Fig. 28c; Supplementary Table 15**), while developmental stage-biased genes were enriched in pathways involved in sensory organ development (*P*_adj_ = 4.96 × 10^−10^), nuclear chromosome segregation (*P*_adj_ = 1.95 × 10^−11^), and adaptive immune response (*P*_adj_ = 3.82 × 10^−13^) (**Supplementary Fig. 29c,d; Supplementary Tables 16 and 17**). Next, we detected a total of 114 sex-biased eGenes (sb-eGenes) across 17 tissues, of which 72 exhibited opposite allelic effects between males and females **(Fig. 6f; Supplementary Table 18)**. For instance, rs427632886 showed opposite effects on the expression of the *TNC* gene, which has been associated with ewe lamb growth^37^, when comparing adipose tissue between males and females (**Fig. 6g**). Furthermore, we detected 138 developmental stage-biased eGenes (db-eGenes) across 19 tissues (**Fig. 6h; Supplementary Table 19**), among which 79 db-eGenes showed opposite allelic effects between developmental stages. For instance, rs406388696 exhibited opposite effects on *AADACL3* in adenohypophysis between adult and juvenile sheep (**Fig. 6i**). This is noteworthy as *AADACL3* has been associated with body weight in a fine-wool sheep population^38^.

### Functional interpretation of GWAS loci using molQTL

To investigate the regulatory mechanisms underlying complex traits in sheep, we integrated molQTL data with GWAS summary statistics for 34 complex traits derived from 11 sheep populations/breeds (median *n* = 2,131), encompassing six trait categories, including body size, reproduction, facial morphology, horn, tail and wool-related traits (**Supplementary Table 20**). Across 34 complex traits, all seven types of molQTL consistently explained more heritability than MAF- and LD-matched SNPs (**Fig. 7a**). This was consistent with previous findings in humans^9^, cattle^11,14^, pigs^12^, and chickens^13^. To further pinpoint the potential causal variants, genes, and tissues underlying complex traits, we conducted colocalization analyses and transcriptome-wide association studies (TWAS) to integrate molQTL with GWAS results of all 34 complex traits. Among the 340 significant GWAS loci examined, 40.3% colocalized (PP_H4_ > 0.8) with at least one molQTL (**Fig. 7b; Supplementary Table 21**), whereas 6.2% and 8.8% were associated with at least one molecular phenotype by single-tissue TWAS (S-PrediXcan)^39^ and multi-tissue TWAS (S-MultiXcan)^40^, respectively (**Extended data figure 9a,b**). The relatively small proportion of TWAS-significant genes may reflect limited predictive performance of TWAS models due to modest tissue sample sizes (**Extended data figure 9c,d**), as well as the small sample size of current GWAS datasets. The proportion of colocalized signals was dependent on the colocalization (PP_H4_) threshold and GWAS significance threshold used (**Extended data figure 9e,f**). To prioritize relevant tissues for complex traits, we adjusted the number of tissue sample size in each tissue (**Supplementary Table 22**). The adenohypophysis, a key endocrine gland that produces and releases several hormones modulating a wide range of biological functions^41^, showed a top relevance for many complex traits, including body size measurements such as cannon bone circumference (CBC) and hip width (HW); reproductive traits such as multiple litter size (MLS); facial features including ear length (EL) and ear width (EW); and hair characteristics such as original/greasy wool weight (OHW) and whiteness index (WHTN) (**Fig. 7c**). Below, we present several examples that demonstrate the potential of the SheepGTEx resource to systematically generate biological hypotheses for future experimental studies.

**Fig. 7.**
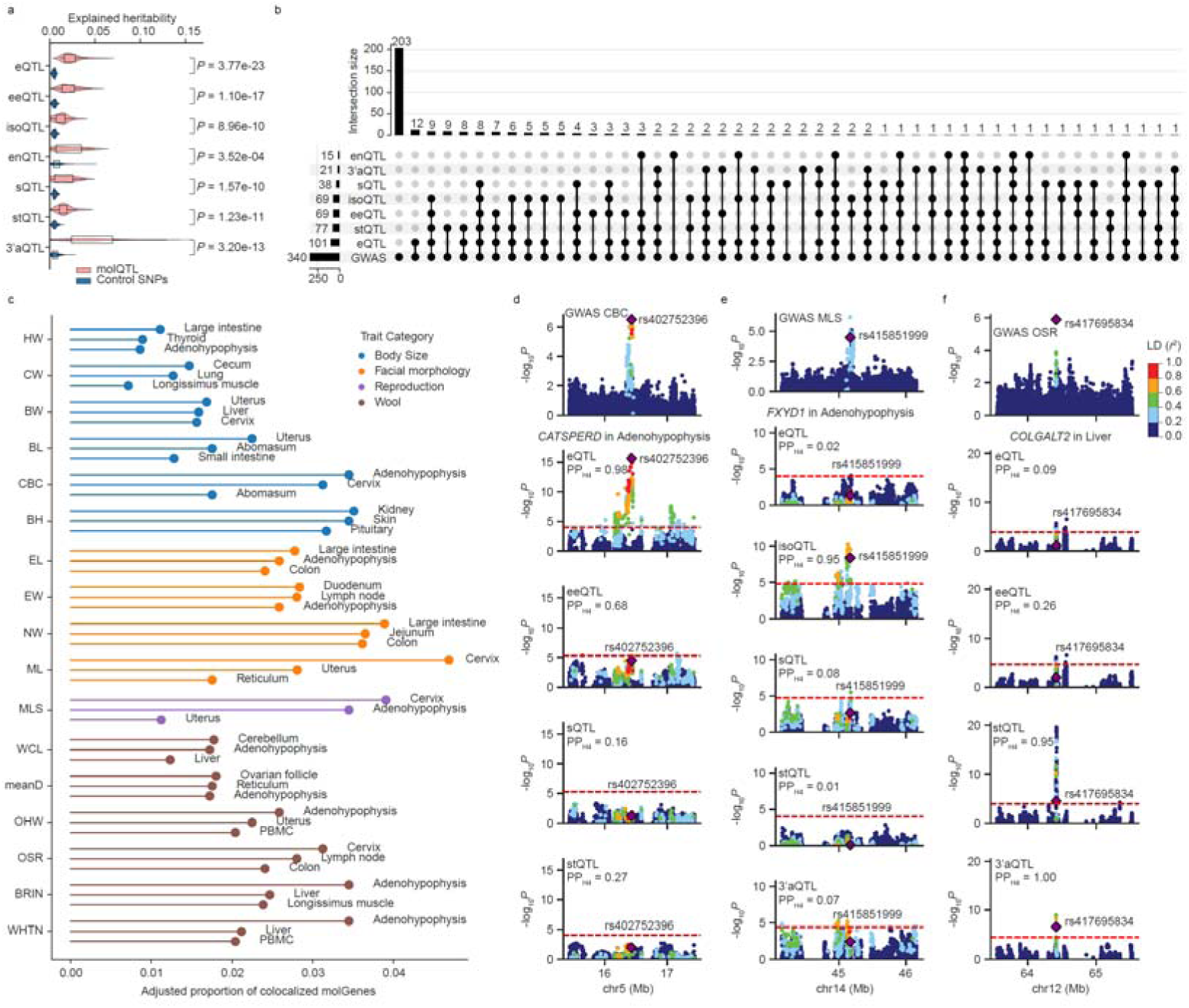
Interpreting GWAS of complex traits using molQTL. **a,** Heritability of 34 complex traits in sheep explained by molQTL and by control SNPs matched for minor allele frequency (MAF) and linkage disequilibrium (LD) score across 51 tissues. Values on the right indicate nominal *P* values (uncorrected) from two-sided paired Student’s *t* tests. **b,** UpSet plot showing the number of GWAS loci colocalizing with at least one molQTL type across 51 tissues using coloc (v5.2.3) package^107^. The bottom matrix displays intersections of loci colocalized with different types of molQTL. **c,** Top three tissues showing the highest adjusted proportions of colocalized molGenes for each trait (trait abbreviations are provided in **Supplementary Table 20**), grouped into four categories: body size, facial morphology, reproduction, and wool traits. Colocalization counts were adjusted for tissue sample size. **d–f,** Regional colocalization plots for molQTL and GWAS loci of three complex traits. SNPs are colored according to LD (*r*^2^) with the colocalized variant. **d**, GWAS locus for cannon bone circumference (CBC) on chromosome 5 (chr5) showing significant colocalization with eQTL (PP_H4_ = 0.98) of *CATSPERD* (rs402752396) specifically in adenohypophysis. **e**, GWAS locus for multiple litter sample (MLS) on chr14 showing significant colocalization with isoQTL (PP_H4_ = 0.95) of *FXYD1* (rs415851999) specifically in adenohypophysis. **f**, GWAS locus for oil/suint ratio (OSR) on chr12 showing significant colocalization with both 3’aQTL (PP_H4_ = 1.00) and stQTL (PP_H4_ = 0.95) of *COLGALT2* (rs417695834) in liver.

The eQTL of *CATSPERD* in adenohypophysis, rather than other molQTL types, showed significant colocalizations with CBC at rs402752396 on chromosome 5 (PP_H4_ = 0.98; **Fig. 7d**). Previous studies on the CatSper gene family members in pigs (including porcine *CATSPER1*–*4*) have reported their expression in hypothalamus and pituitary^42^. These observations raise the hypothesis that *CATSPERD* may have functional roles within adenohypophysis, potentially influencing overall body size through neuroendocrine mechanisms, and thereby contributing to variation in CBC. In addition, the stQTL of *DUS3L* in uterus, rather than other molQTL types, also colocalized with CBC at the same variant (PP_H4_ = 0.96; **Extended data figure 9g**).

The isoQTL rather than other molQTL of *FXYD1* in adenohypophysis colocalized with MLS at rs415851999 on chromosome 14 (PP_H4_ = 0.95; **Fig. 7e**). *FXYD1* encodes a transmembrane modulator of Na□/K□-ATPase activity, enriched in neural tissues, which likely contributes to maintaining GnRH neuron excitability by regulating ionic homeostasis after neuronal activation^43^. The developmentally regulated expression of *FXYD1* around pubertal onset and presence in approximately half of GnRH neurons in rodents supports a potential role in the neuroendocrine regulation of reproductive maturation^43^.

The 3’aQTL (PP_H4_ = 1.00) and stQTL (PP_H4_ = 0.95) of *COLGALT2* in liver colocalized with oil/suint ratio (OSR) at rs417695834 on chromosome 12 (**Fig. 7f**). *Colgalt2−/−* mice exhibited lipodystrophy, hepatic steatosis, and altered body weight, with phenotypes further aggravated under high-fat or methionine- and choline-deficient diets^44^. Transcriptomic profiling revealed dysregulation of lipid metabolic pathways, including AMPK and adipocytokine signaling, alongside reduced expression of key lipogenic genes (*Cidea*, *Cidec*, *Cd36* and *Pparg*)^44^. These findings suggest that *COLGALT2* regulates hepatic fat deposition and adipocyte development in sheep primarily through transcriptional control of lipid metabolic networks, thereby potentially influencing phenotypes related to wool oil content. Additionally, the 3’aQTL (PP_H4_ = 0.96) and eeQTL (PP_H4_ = 0.99) of *COLGALT2* in muscle also colocalized with OSR at the same variant (**Extended Data Fig. 9h**). Its role in muscle collagen glycosylation and maturation may affect fat distribution and thus wool quality traits like oil content^45^. Therefore, *COLGALT2* likely impacts OSR by coordinating lipid metabolism in liver and collagen metabolism in muscle, modulating wool fiber oil content.

### Deciphering the regulatory architecture underlying population selection and domestication

To investigate the regulatory mechanisms underlying selection, we conducted a fixation index (*F*_ST_) test between 710 EUR and 1,104 CEA sheep (**Fig. 8a; Supplementary Table 5**), which revealed well-known genes previously detected as under selection (e.g., *RXFP2*, *IRF2BP2*, *BMP2* and *PDGFD*)^46,47^. We then evenly divided the genome-wide regions into 10 *F*_ST_ deciles from the smallest (non-selected regions) to the largest (highly selected) and identified a total of 386 non-overlapping candidate selection regions (spanning 40,97 Mb with 477,685 SNPs) defined by the top 1% *F*_ST_ decile (**Supplementary Table 23**). Broadly, tissue-shared eQTL tended to be enriched in non-selected regions, whereas tissue-specific eQTL were disproportionately enriched in the top *F*_ST_ deciles and showed a higher enrichment than tissue-shared eQTL (**Fig. 8b; Extended Data Fig. 10a,b**). Among all the 51 tissues being tested, eQTL specific in ovary and abomasum showed the highest enrichment in the top selected regions, a result that was consistent across varying *F*_ST_ thresholds (**Fig. 8c; Extended Data Fig. 10c–f**). Furthermore, compared to standard eQTL, anc-eQTL showed a higher enrichment top-selected regions, and confirmed that ovary and abomasum were the top-relevant tissues (**Fig. 8d; Extended Data Fig. 10g**). Similar tissue-specific enrichment patterns were observed for non-primary eQTL and other types of molQTL in regions under strong selection (**Extended Data Fig. 10h,i**). Consistent with the “rare variant, large effect” model often seen in complex traits^31^, we observed a pervasive negative correlation between MAF and effect size across all seven molQTL types, ranging from the strongest signal in sQTL (Pearson’s *r* = −0.35) to the weakest in eQTL (Pearson’s *r* = −0.23) (**Extended Data Fig. 10j**). The molQTL with non-primary effects, tissue-specific activity, and lower pleiotropic complexity appeared to be under stronger selective constraint (**Extended Data Fig. 10k–m**). As an illustrative case, a CEA-specific fd-eQTL (rs423495724 as lead variant) regulating *CCL25* expression in ovary, was located within a region under strong selection in EUR sheep (**Fig. 8e–h**). This gene was associated with fetal development in sheep and ovulation-related disorders in humans and cattle^48–50^. The A allele, which increased *CCL25* expression in both ancestries, was nearly fixed in EUR (frequency = 0.98) but remained polymorphic in CEA with a frequency of 0.77 (**Fig. 8i**). Another interesting example involved *RIPK1* in abomasum: its lead eQTL (rs404131822) resided within a EUR-selected region (MAF = 0.03) but exhibited no regulatory activity in CEA (MAF = 0.08) (**Fig. 8j–n**). This gene has been implicated in human obesity, starvation resistance in mice, and heat stress response in sheep^51–53^.

**Fig. 8.**
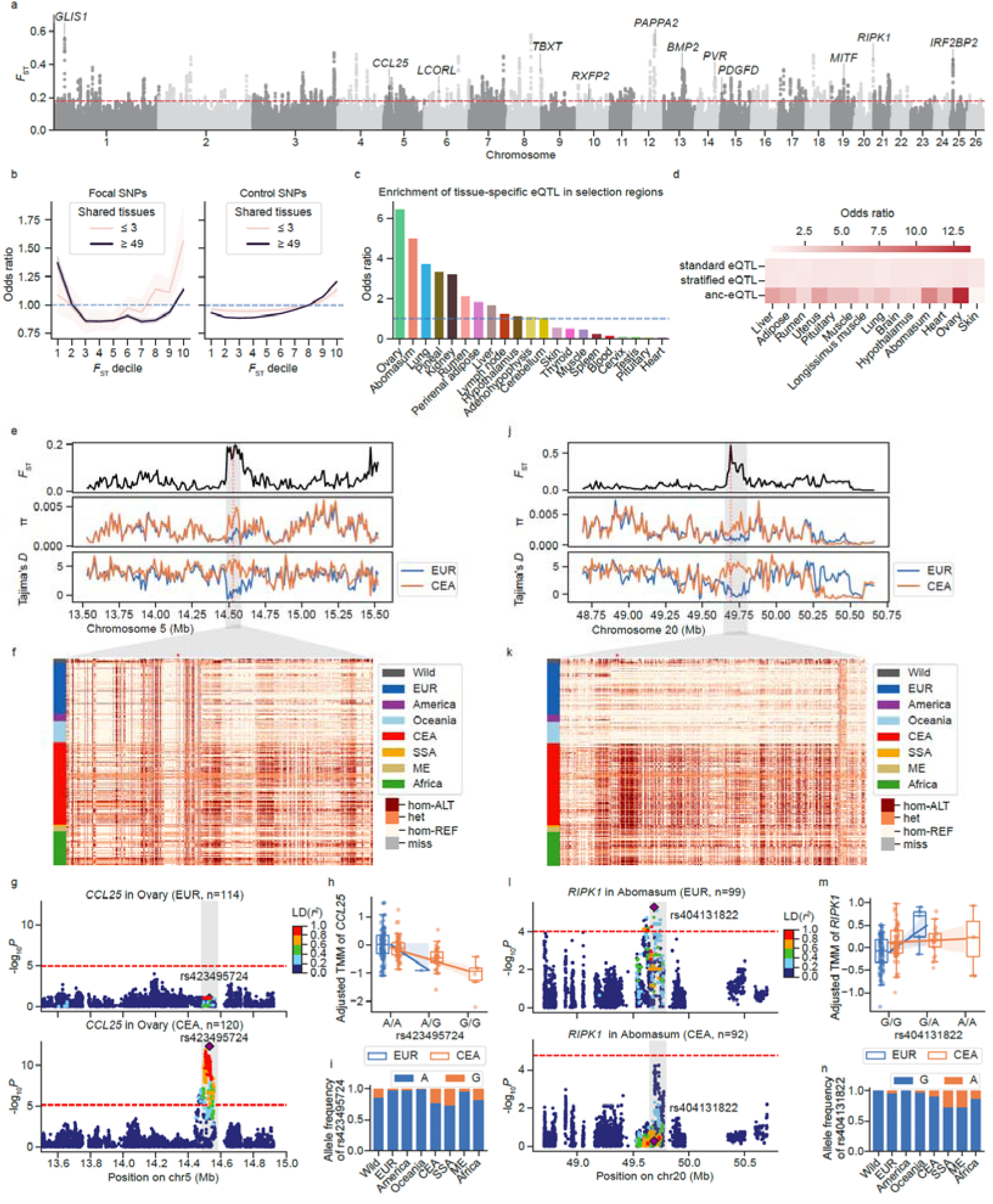
Regulatory variants underlying selection signatures between European and Asian domestic sheep. **a**, Manhattan plot of fixation index (*F*_ST_) values between modern European (EUR, *n* = 710) and Central and East Asian (CEA, *n* = 1,104) ancestry groups. The dashed red line indicates the top 1% cutoff. Genes previously reported in sheep are labelled in the plot. **b**, Enrichment of *cis*-eQTL shared in ≤ 3 tissues or ≥ 49 tissues in genomic regions grouped by *F*_ST_ deciles. Focal *cis*-eQTL SNPs are shown on the left, with control SNPs matched for minor allele frequency (MAF) and linkage disequilibrium (LD) score on the right. **c**, Enrichment of tissue-specific eQTL in highly selected regions (top 1% *F*_ST_) across 51 tissues. **d**, Enrichment of standard eQTL (using all samples in each tissue), stratified eQTL (eQTL mapping separately in EUR and CEA ancestry groups), and anc-eQTL in highly selected regions across 14 tissues. **e**, Genomic profiles of *F*_ST_, π, and Tajima’s *D* around the *CCL25* selected region (shaded area identified by all three methods). **f**, Haplotype structure of the *CCL25* selected region. Rows represent individuals and columns represent polymorphic sites. The target eQTL (rs423495724) is marked by an asterisk. hom-ALT, homozygous alternate; het, heterozygous; hom-REF, homozygous reference. **g**, Manhattan plot for *cis*-eQTL of *CCL25* in ovary in the EUR (top) and CEA groups (bottom). SNPs are colored by LD (*r*^2^) with the target variant. **h**, Expression, defined as the adjusted trimmed mean of M values (TMM), of *CCL25* across genotypes of the lead SNP rs423495724 in ovary. **i**, Allele frequency of the SNP rs423495724 across global sheep populations. **j**, Distributions of *F*_ST_, π, and Tajima’s *D* around the *RIPK1* selected region (shaded area). **k**, Haplotype structure of the *RIPK1* selected region (shaded area). The target eQTL (rs404131822) is marked by an asterisk. **l**, Manhattan plot for *cis*-eQTL of *RIPK1* in abomasum in EUR (top) and CEA groups (bottom). **m**, Expression of *RIPK1* across rs404131822 genotypes in abomasum. **n**, Allele frequency of rs404131822 across sheep populations. Sheep population abbreviations: EUR, Europe; CEA, Central and East Asia; SSA, South and Southeast Asia; ME, Middle East.

To investigate temporal dynamics in regulatory effects throughout extended domestication processes over the past 10,000 years, we analyzed 52 publicly available ancient sheep genomes (mean coverage ≥ 0.5×) from Europe and Asia^1,54–56^ (**Fig. 9a; Supplementary Figs. 31 and 32; Supplementary Table 24**). After genotype imputation, we obtained 27,866,951 common SNPs (MAF > 0.01), with a mean imputation accuracy of 90.0% (**Methods; Supplementary Note; Supplementary Fig. 31a–e**). We identified selection signals (*F*_ST_) in ancient European and Asian sheep separately by comparing samples dated to > 4,000 years before present (YBP) with those dated < 4,000 YBP (**Fig. 9b; Supplementary Fig. 33a; Supplementary Tables 25 and 26**). In general, selection signals were correlated between European and Asian populations (**Fig. 9c**). We confirmed that regions associated with coat color and pigmentation (e.g., *PDGFRA*, *KIT*, *MC1R*, and *IFT88*) and wool traits (e.g., *TCF3* and *EDN3*) were under strong selection^1,57,58^ (**Fig. 9b; Supplementary Fig. 33a**). Enrichment of tissue-specific eQTL in top-selected regions pinpointed skin and cervix as the top tissues with significant regulatory divergence over time, which was consistent in both European and Asian populations (**Fig. 9d; Supplementary Fig. 31f**). This suggests that the physiological functions of integumentary and reproductive organ systems were under strong selection over time. Furthermore, we predicted multi-tissue gene expression for all the 52 ancient individuals using the Elastic Net-based TWAS model trained on the SheepGTEx data (**Methods**). We then identified four gene clusters across five archaeological periods (**Fig. 9e,f; Supplementary Fig. 31g**). For example, 449 skin genes in cluster 2 showed increasing expression, while 789 cervix genes in cluster 1 showed decreasing expression, across the five archaeological periods (**Fig. 9e,f**). A representative gene, *THNSL2*, associated with wool traits and skin wrinkles^59^, was regulated by rs406457274, which exhibited selection for the A allele alongside increasing gene expression in skin over time (**Fig. 9g**). Similarly, lead variants 1_197848536 of *CLDN16* (associated with coat color)^60^ and rs422778558 of *PPIH* (associated with wool shedding)^61^ showed parallel allele frequency changes and also gene expression shifts in skin (**Supplementary Fig. 33b,c**). In cervix, the frequency of C allele at rs427632973 dropped over time, leading to the reduced expression of *STK10* (**Fig. 9h**), which was relevant to reproduction traits^62^. Interestingly, skin-related regulatory variants (e.g., those affecting *THNSL2*, *CLDN16*, and *PPIH*) also exhibited broad regulatory effects across various tissues (e.g., rumen and adenohypophysis) (**Fig. 9i; Supplementary Fig. 33d–h**), whereas such pleiotropy was not observed for *STK10* in cervix (**Fig. 9j; Supplementary Fig. 33i**), suggesting that skin-associated regulatory variants may play broader roles in gene regulation across tissues^63,64^.

**Fig. 9.**
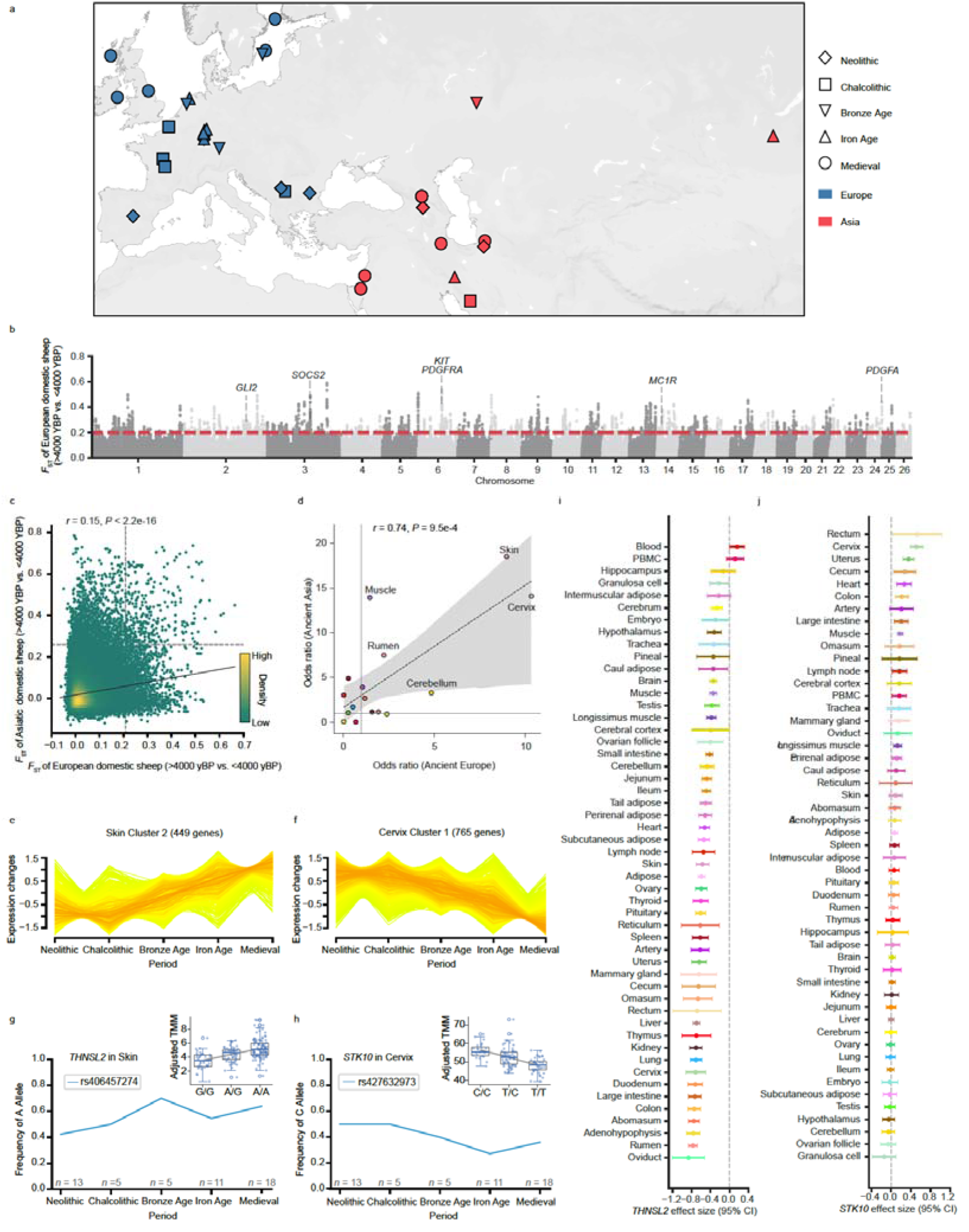
Spatiotemporal genomic differentiation and tissue-specific regulatory evolution in ancient domestic sheep. **a**, Geographic distribution of 52 ancient sheep samples across regions and archaeological periods. **b**, Manhattan plot of fixation index (*F*_ST_) values between ancient European sheep dated to > 4,000 vs. < 4,000 years before present (YBP). The dashed red line indicates the top 1% threshold. Genes previously reported in sheep are labelled in the plot. **c**, Density plot comparing *F*_ST_ values for SNPs in ancient European and Asian sheep (> 4,000 vs. < 4,000 YBP). Dashed lines show top 1% thresholds. **d**, Comparison of enrichment (odds ratio) of tissue-specific *cis*-eQTL in genomic regions under strong selection in ancient European or Asian sheep populations. Ordinary least squares (OLS) regression indicates significant Pearson’s correlations (two-sided Student’s *t*-test) in **c** and **d**. **e–f**, Temporal trajectories of genetically predicted gene expression in ancient European sheep, showing increased expression in skin (**e**) and decreased expression in cervix (**f**). **g–h**, Allele frequency changes of SNPs rs406457274 (**g**) and rs427632973 (**h**) over time, alongside expression of *THNSL2* across three genotypes in skin and *STK10* in cervix, respectively. Sample sizes for gene expression: G/G = 39, A/G = 54, A/A = 110 for *THNSL2*; C/C = 24, T/C = 62, T/T = 42 for *STK10*. **i–j**, eQTL effect sizes of rs406457274 on *THNSL2* expression (**i**), and rs427632973 on *STK10* expression (**j**) across tissues, with 95% confidence intervals.

## Discussion

The pilot SheepGTEx, built on 6,761 RNA□seq samples of 51 tissues from 2,816 animals worldwide, provides a valuable catalogue of regulatory variants in sheep. It includes 159,140 independent eQTL of 19,213 eGenes, 89,839 eeQTL of 12,224 eeGenes, 91,341 isoQTL of 9,587 isoGenes, 19,386 enQTL of 3,904 enGenes, 57,877 sQTL of 9,212 sGenes, 80,099 stQTL of 10,345 stGenes, and 18,236 3’aQTL of 4,231 3’aGenes, thus revealing the polygenic nature of molecular phenotypes. By jointly analyzing seven types of transcriptomic phenotypes, we showed both distinct and pleiotropic regulatory effects, highlighting the value of integrating diverse molQTL for a more comprehensive view of gene regulation^13^. Furthermore, this multi-tissue SheepGTEx atlas provided novel insights into the regulatory mechanisms underlying complex traits such as growth, reproduction, morphology and wool traits in sheep. This allowed us to link 40.3% of GWAS loci to molecular phenotypes, genomic variants, target genes, and relevant tissues, similar to findings in humans and other livestock species^9,12–14^. For instance, *CATSPERD* and *FXYD1*, genes traditionally associated with reproductive and neural functions^42,43^, exhibited strong colocalizations with cannon bone circumference (CBC) and multiple litter size (MLS) in adenohypophysis, respectively, highlighting adenohypophysis as a key regulatory hub linking growth and reproductive traits in sheep. Moreover, we demonstrated the temporal nature of regulatory variants frequencies due to population selection and domestication over the past 10,000 years in sheep. These results echo previous findings in humans showing that population□selected or introgressed regulatory variants have driven local adaptation, such as breast cancer risk alleles in Peruvians^65^, UV-resistance loci in Han Chinese^66^, and Neanderthal□derived regulatory variants enriched in Eurasians and linked to immune function^67,68^.

Despite these advances, we noted several limitations that will be addressed through future development of the SheepGTEx Project. First, our current analyses were limited to common SNPs. More RNA-seq samples with paired WGS, improved reference genomes (i.e., telomere-to-telomere genome^69^ and pangenome^70^), and enhanced functional annotations (i.e., by integrating CAGE data to refine transcription start site resolution and mitigate tissue-specific annotation artifacts^71,72^) will enable us to explore regulatory effects of rare variants, transposable elements, short tandem repeats and complex structural variants. Second, this phase of SheepGTEx was focused on bulk short-read RNA-seq data. Expanding additional omics, such as long-read isoform sequencing, single-cell RNA-seq, and epigenomics, will improve functional annotation of genomic variants. Third, the statistically inferred regulatory variants cannot be treated as causal variants. CRISPR-based perturbation^73^ and high-throughput reporter assays (e.g., STARR-seq^74^) will be critical for systematically validating putative regulatory variants *in vivo* and *in vitro*. Fourth, context□dependent molQTL (ancestry□, sex□, and developmental stage□specific) remain understudied in this study due to limited sample size, residual confounding, and metadata uncertainty in publicly available datasets. This underscores the need for larger, well-controlled cohorts to uncover such gene-by-environment interactions and better capture the genetic architecture of complex traits^75,76^. Finally, large language models (LLMs) have recently garnered considerable attention. They are being increasingly applied in biological research, including for characterizing regulatory variants^77^, offering promising opportunities to overcome limitations of LD in population-based molQTL mapping. In the future, incorporating LLM-based approaches may enable more accurate prediction of regulatory variants and improved functional fine-mapping and causal inference, thereby improving our understanding of the molecular mechanisms underlying complex traits. Collectively, a fully developed SheepGTEx will serve as a comprehensive and robust reference panel for deciphering the genetic and molecular architecture underlying complex traits and selective breeding in sheep.

## Online Methods

### Ethics and sample collection

All experimental procedures in this study were approved by the Science Research Department, which oversees animal welfare, at the Institute of Animal Sciences, Chinese Academy of Agricultural Sciences (IAS-CAAS, Beijing, China). Ethical approval for animal use and survival was granted by the IAS-CAAS Animal Ethics Committee (No. IAS2024-129). Detailed description for tissue collection, RNA extraction, and RNA sequencing for the datasets generated specifically for this project are provided in **Supplementary Note**.

### Bulk RNA-seq data processing

We compiled a total of 9,717 bulk RNA-seq datasets, comprising 7,817 publicly available raw datasets (retrieved from the NCBI Sequence Read Archive as of December 16, 2023) and 1,900 newly generated samples (**Supplementary Note; Supplementary Fig. 1a–f; Supplementary Table 1**). We then processed all data using a unified analytical pipeline. Briefly, adapter trimming and quality filtering were performed using fastp (v0.23.2)^78^ with the parameters “-f 3 -t 3 -l 36 -r -W 4 -M 15”, followed by two-pass alignment of clean reads to the sheep reference genome (ARS-UI_Ramb_v2.0^24^, GCF_016772045.1) using STAR (v2.7.10b)^79^ with the “--twopassMode Basic” option. Gene-level quantification was performed with featureCounts (v2.0.5)^80^, yielding raw read counts for 21,257 PCGs and 4,442 lncRNAs annotated in NCBI *Ovis aries* Annotation Release 104 (ARS-UI_Ramb_v2.0, GCF_016772045.1). Normalized transcript abundance (transcripts per million, TPM) was calculated using StringTie (v2.1.1)^81^. We retained 8,621 high-quality samples with ≥ 10 million clean reads, uniquely mapping rates ≥ 60%, and detectable expression (TPM□≥□0.1) in ≥ 50% of annotated genes. To remove outlier samples that did not cluster with their labelled tissue, we assessed sample-level expression variance using hierarchical clustering with *hclust* function in R (v4.3.0)^82^ based on log_2_(TPM□+□0.25) of the 5,000 (∼20%) most variable genes (ranked by standard deviation) and *t*-distributed stochastic neighbor embedding (*t*-SNE)^83^, resulting in a final dataset of 8,171 RNA-seq samples for downstream analysis (**Supplementary Fig. 1g–o; Supplementary Table 1**). Unknown batch effects were estimated as gene expression principal components (PCs) within tissues using the PCAForQTL package^84^ and removed using the *removeBatchEffect* function implemented in the limma package (v3.56.2)^85^. The tree clustering of all samples before and after batch correction was visualized with the ggtree (v3.14.0)^86^ and ggtreeExtra (v1.16.0)^87^ packages (**Extended Data Fig. 1a; Supplementary Fig. 2c**).

### Genotype imputation for RNA-seq samples using a multi-breed reference panel

To construct a comprehensive sheep genotype imputation reference panel, we analyzed existing whole-genome sequencing (WGS) data from 3,125 animals worldwide (**Supplementary Table 5**). This panel encompassed 249 modern domestic breeds/populations and seven wild species, spanning Asia (*n* = 1,236), Europe (*n* = 715), Africa (*n* = 459), Oceania (*n* = 293), and the Americas (*n* = 97). Raw reads were trimmed using Trimmomatic (v0.39)^88^ (parameters: “MINLEN:50 LEADING:20 TRAILING:20 SLIDINGWINDOW:5:20”) to remove adapter sequences and low-quality bases, and subsequently aligned to the sheep reference genome (ARS-UI_Ramb_v2.0, GCA_016772045.1) using BWA-MEM (v0.7.5a-r405)^89^. Aligned reads were sorted with SAMtools (v1.9)^90^, and PCR duplicates were marked and removed using Picard (v2.21.2) (https://broadinstitute.github.io/picard). Variant calling was conducted with GATK (v4.1.4.1)^91^. Per-sample gVCF files were generated using HaplotypeCaller, merged via CombineGVCFs and jointly genotyped with GenotypeGVCFs. SNPs were filtered using VariantFiltration with the following parameters: QD < 2.0, FS > 60.0, MQRankSum < −12.5, ReadPosRankSum < −8.0, SOR > 3.0, and MQ < 40.0. High-confidence variants were retained using BCFtools (v1.9)^90^ with the following criteria: (1) biallelic SNPs; (2) missing rate < 0.1; (3) minor allele count (MAC) > 2; and (4) read depth between 0.33× and 3× of the mean depth of all individuals. The final dataset of 60,840,500 high-quality SNPs was imputed and phased using Beagle (v5.4)^92,93^ to generate a multi-breed haplotype reference panel.

For genotype imputation of RNA-seq samples, we marked and removed PCR duplicates, split reads that contained ‘N’s in their CIGAR string and performed base quality recalibration using GATK (v4.3.0.0), with 60,840,500 high-quality SNPs from the reference panel serving as known variants set. Genotype imputation was then performed using GLIMPSE2 (v2.0.0)^94^ with the parameters “--mapq 20 --ne 1000”, based on the above reference panel. After rigorous evaluation and validation (**Supplementary Note; Extended Data Fig. 2**), we retained common variants with INFO ≥ 0.75 and MAF ≥ 0.05, yielding 3,427,511 SNPs for subsequent molQTL mapping. To mitigate potential artifacts from allelic mapping bias in splicing quantification and ASE analysis, RNA-seq reads were re-aligned using STAR with the “--waspOutputMode SAMtag” option^95^ guided by imputed genotypes, and reads failing WASP filtering (i.e., tagged with “vW:i:[2-7]”) were excluded.

To validate imputed genotypes, we analyzed 2,728 RNA-seq samples with matched WGS from nine independent populations/breeds and assessed accuracy using genotype concordance rate (CR) and squared Spearman’s correlation (ρ^2^) between RNA-seq and WGS data (**Supplementary Note**; **Extended Data Fig. 2**).

### Quantification of molecular phenotypes

To quantify gene expression, we used featureCounts (v2.0.5) to obtain raw read counts for 25,699 PCGs and lncRNAs from NCBI *Ovis aries* Annotation RefSeq Release 104 (ARS-UI_Ramb_v2.0, GCA_016772045.1) and normalized their expression to transcripts per million (TPM) across 8,171 RNA-seq samples. Given the limited number of annotated lncRNAs, we jointly performed gene expression normalization and downstream eQTL mapping for 21,257 PCGs and 4,442 lncRNAs, despite their potentially distinct regulatory roles^12,13^ (**Supplementary Fig. 3d–g**). The quantification of individual exons was also performed with featureCounts with the options “-f -t exon”, yielding read counts for 252,949 exons from 23,189 multi-exonic genes. Isoform expression was quantified using Salmon (v1.10.0)^96^, with 54,667 isoforms retained from 11,746 genes with multiple transcripts. For enhancer expression, we compiled 323,820 nonredundant predicted active enhancers (EnhA) from 43 tissues and restricted analysis to 81,004 intergenic transcribed enhancers to mitigate contamination from gene transcription (https://genome.ucsc.edu/s/mengzhu/Ramb_v2.0_regulatory_elements). Enhancer read counts were quantified using featureCounts with the Simplified Annotation Format (SAF) annotation file (-F SAF -a), a tab-delimited annotation format defining genomic regions used for read assignment, derived from the 81,004 sheep intergenic transcribed enhancers.

Alternative splicing was quantified using Leafcutter (v0.2.7)^97^ on WASP-filtered alignments. Junction counts were extracted with the bam2junc.sh script, intron clusters were defined with leafcutter_cluster.py (parameters: “--min_clu_reads 30 --min_clu_ratio 0.001 --max_intron_len 500000”). Clusters were mapped to genes using map_clusters_to_genes.R. Introns were filtered following published criteria^9,12,13^, excluding those with > 50% missing or zero counts, fewer than max(10, 0.1*n*) unique values (*n* = sample size), or low complexity (∑*_i_*(|*z_i_*| < 0.25) ≥ n − 3 and ∑*_i_*(|*z_i_*| > 6) ≤ 3, where *z_i_* is the *z* score of the *i*th cluster read fraction across individuals). The remaining counts were normalized using prepare_phenotype_table.py. RNA stability was assessed following the Pantry framework^98^. We quantified read counts of constitutive exons and introns (defined via the CRIES pipeline^18^ in https://github.com/csglab/CRIES) using featureCounts. Constitutive exon/intron ratios were computed per gene, with those showing > 50% missing values removed, followed by K-nearest neighbor (KNN) imputation and two-stage quantile normalization. For 3’UTR APA, we employed a unified protocol^99^ implemented in DaPars2 (v2.1)^100^. Briefly, 3’UTR annotations were extracted with the DaPars_Extract_Anno.py script, and genome coverage tracks were generated using BEDTools (v2.31.0)^101^. PolyA events were then quantified with the Dapars2_Multi_Sample.py script, yielding PDUI (percentage of distal polyA site usage index) values. Sample-level PDUI matrices were merged using the merge_apa_quant_res_by_chr.R script. Finally, Genes with > 50% missing PDUI entries and individuals with > 80% missing data were removed, and the final PDUI matrix was processed using KNN imputation, removal of constant features, and quantile normalization.

### molQTL mapping

After sample deduplication, 6,761 RNA-seq samples of 51 tissues from 2,816 animals were retained for molQTL mapping (**Supplementary Note; Supplementary Figs. 1o, and 9**; **Supplementary Table 6**). Within each of 51 tissues, SNPs with MAF ≥ 5% and minor allele count ≥ 6 were kept for subsequent analysis using PLINK (v1.90b7)^102^. For abundance-based molecular phenotypes (i.e., gene, exon, isoform, and enhancer expression), we excluded phenotypes with TPM□<□0.1 or raw read counts□<□6 in > 20% samples within each tissue. Read counts of remaining phenotypes were normalized using trimmed mean of M values (TMM)^103^, followed by the inverse normal transformation. Structure-based molecular phenotypes (i.e., alternative splicing, RNA stability, and 3’UTR APA) were processed and normalized per tissue as described in **molecular phenotype quantification**. To account for technical, biological, and population-related confounders, we computed phenotype principal components (PCs) using the PCAForQTL package^84^ and genotype PCs using PLINK for each tissue. The number of phenotype PCs was automatically inferred with the *runElbow* function implemented in PCAForQTL. Covariates used for molQTL mapping included inferred phenotype PCs, the top five genotype PCs (for tissues with sample size < 200) or top ten genotype PCs (≥ 200), as well as sex and age. Highly colinear covariates (Pearson’s *r* > 0.9) were excluded.

We performed *cis*-molQTL mapping using TensorQTL (v1.0.9)^25^ with a linear regression model, accounting for all covariates above. The *cis*-window was defined as ±1□Mb around the TSS of each corresponding gene. To identify significant molGenes and molQTL, we applied the same two-step multiple-testing correction framework used in the human GTEx and other FarmGTEx projects^9,10^. Briefly, for all variant-molecular phenotype pairs, nominal associations were computed using the “--mode cis_nominal” option. In parallel, we employed permutation testing to derive empirical *P* values for each gene using the “--mode cis” option. For *cis*-eeQTL, *cis*-isoQTL, *cis*-enQTL, *cis*-sQTL and *cis*-3’aQTL, we grouped features by gene using the “--phenotype_groups” option. Genes significantly regulated by at least one variant (molGenes) were identified by controlling the false discovery rate (FDR) < 5% via Benjamini-Hochberg correction on Beta-approximated empirical *P* values. To determine significant *cis*-molQTL, we derived gene-level *P* value thresholds from the fitted Beta distribution using qbeta(pt, beta_shape1, beta_shape2) in R, where pt is the genome-wide empirical *P* value threshold closest to FDR = 0.05, and beta_shape1, beta_shape2 are the fitted Beta distribution parameters estimated by TensorQTL. Variants with nominal *P* values below the gene-level threshold were reported as significant *cis*-molQTL.

### Conditional and fine-mapping analysis of molQTL

To detect multiple independent *cis*-molQTL signals for each molGene, we employed two complementary approaches. First, we performed conditional analysis using a forward-backward stepwise regression procedure^104^, implemented in TensorQTL (--mode cis_independent). For each molecular phenotype, molQTL were ranked by their conditional association strength, with the top-ranked variant (rank□=□1) defined as the primary signal and lower-ranked variants (rank□>□1) as non-primary signals. Second, we applied the Sum of Single Effects (SuSiE) regression framework to fine-map likely causal variants, using the *susie_rss* function^105^ from the susieR package (v0.12.35)^106^ with default settings. For each molGene, the SuSiE model was fitted using molQTL summary statistics, and credible sets were derived by aggregating variants whose posterior inclusion probabilities (PIPs) cumulatively summed to a value of 95% or higher.

### Comparison between eQTL and other types of molQTL

To assess whether different types of molecular phenotypes shared common genetic regulatory mechanisms, we compared eQTL with other types of molQTL derived from the same gene. First, we conducted Bayesian colocalization analysis using the *coloc.abf* function in the coloc (v5.2.3) package^107^, which computed posterior probabilities (PP) for five hypotheses: H_0_ (no association with either phenotype), H_1_ (association with phenotype1 only), H_2_ (association with phenotype2 only), H_3_ (distinct signals for phenotype1 and phenotype2), and H_4_ (shared causal variant for both phenotypes). molGenes with PP_H4_□>□0.5 were considered to exhibit colocalized regulatory effects between eQTL and other molQTL. Second, we calculated the pairwise LD (*r*^2^) between lead variants of eQTL and those of corresponding molQTL using PLINK with the parameter “--ld”. Additionally, we applied OPERA (v1.18)^108^ for eQTL-molQTL associations, which implements a Bayesian extension of Summary data-based Mendelian randomization (SMR) and heterogeneity independent instruments (HEIDI) methods for joint analysis of multi-omics QTL summary statistics. We defined pleiotropic associations as those with posterior probabilities of association (PPA) ≥□0.9.

### Tissue-sharing/specificity analysis of molQTL

To investigate the sharing pattern of regulatory effects among tissues, we performed cross-tissue analyses of *cis*-molQTL using *mashr* (v0.2.79)^109^. We fitted the *mash* model using the top molQTL of each molGene and a random subset of one million variant-gene pairs from the nominal association results across all tissues. For each pair, normalized effect sizes (Z-scores, calculated as slope/slope_se from the TensorQTL outputs) were used as input. Missing Z-scores were imputed as 0 with a standard error set to 1□×□10^6^. Posterior effect sizes and local false sign rates (LFSR), as estimated by *mashr*, were used to infer molQTL magnitude and significance, respectively. A molQTL was considered active in a given tissue if the respective LFSR□<□0.05. To quantify cross-tissue similarity in regulatory effects, we further computed pairwise Spearman’s correlation coefficients (ρ) for the effect sizes of molQTL between tissues, restricting to variants with LFSR□<□0.05 in at least one tissue. Hierarchical clustering was then performed on the maximum distance matrix of ρ using complete linkage in the *hclust* function in R (v4.4.1).

To assess consistency of tissue-clustering patterns across all data types (that is, seven types of molecular phenotypes and their corresponding molQTL), we performed k-means clustering (k = 2–20) with a maximum of 50,000 iterations per run using the *kmeans* function from the stats package (v4.4.1). For each k value, we computed the pairwise Rand index to evaluate clustering similarity using the *rand.index* function implemented in the fossil package (v0.4.0)^110^.

To investigate the heterogeneity of regulatory effect size among tissue-shared eGenes, we compared *cis*-eQTL effect sizes between tissues. For each tissue pair, if the lead *cis*-eQTL of an eGene was identical or in strong LD (*r*^2^ > 0.8) across tissues, we classified the eGene-tissue pairs into three categories based on the overlap of the 95% confidence intervals (CI) of effect sizes: homogeneous (overlapping CI), heterogeneous (non-overlapping CI but concordant direction of effect), and opposite (discordant directions of effect).

### Context-specific molQTL analysis

To investigate the context-specificity of *cis*-eQTL regarding genetic background, we adapted a previously described framework^36^ with minor modifications to identify ancestry-specific eQTL (anc-eQTL) between individuals from European (EUR) and Central and East Asia (CEA) ancestries (**Fig. 6a**). Global ancestry was estimated by ADMIXTURE (v1.3.0)^111^ within each tissue using EUR and CEA samples. To ensure clear genetic assignments, we excluded individuals whose estimated proportion of self-identified ancestry was less than 50%. A total of 14 tissues with ≥ 40 samples in each of two ancestry groups were retained for downstream analyses. We then conducted *cis*-eQTL mapping and fine-mapping analysis separately in the EUR and CEA groups using the same procedures as described in **molQTL mapping** and **Conditional and fine-mapping analysis of molQTL**. The anc-eQTL were further classified into three categories based on differences in allele frequency, LD, and effect size (**Fig. 6a**). More specifically, for non-overlapping 95% credible sets between populations, variants were defined as fd-eQTL if they were rare (MAF□<□0.01) in one ancestry but common (MAF□≥□0.05) in the other one. For eGenes detected in either group, we further applied MESuSiE (v1.0)^112^, which accounted for population-specific LD patterns, and defined ld-eQTL as variants with a posterior inclusion probability (PIP)□>□0.5 for ancestry-specific signals. The he-eQTL were defined as cases where the (1□−□0.05/ngene) confidence intervals (CIs) of lead eQTL effects did not overlap, where ngene is the number of genes tested. Genes with at least one such variant were designated as ancestry-specific eGenes (anc-eGenes).

We also applied this framework to assess differences in regulatory effects between the sexes and between developmental stages (adult vs. juvenile), considering he-eQTL as sex-biased or development-biased eQTL, respectively. Genes with at least one such variant were classified as sex-biased eGenes (sb-eGenes) or developmental stage-biased eGenes (db-eGenes), respectively.

### GWAS summary statistics and SNP heritability estimation

To elucidate the regulatory mechanisms underlying complex traits in sheep, we integrated molQTL data with GWAS summary statistics of 11 populations/breeds across 34 economically important traits, spanning six categories, including body size, reproduction, facial morphology, horn, tail and wool-related characteristics (**Supplementary Table 20**). Meta-analyses across multiple sheep populations were conducted for each trait using METAL (v2020-05-05)^113^. For integrative analyses, given the limited power of the GWAS, we considered suggestively significant loci (*P* < 1 × 10^−5^). Each locus was defined as a genomic interval encompassing contiguous significant SNPs within 1□Mb, centered on the lead variant. The locus boundaries were extended by up to 0.5□Mb on either side until the −log_10_*P* decreased by more than four units relative to the lead SNP^114^. This procedure identified 340 GWAS loci that overlapped with at least one significant molQTL. To quantify the contribution of regulatory variants to complex trait, we employed SumHer implemented in LDAK (v6.1)^115^ using meta-GWAS summary statistics to estimate SNP-based heritability for each trait. We then compared the proportion of phenotypic variance explained by molQTL SNPs versus that explained by MAF- and LD-matched background SNPs. Tagging files were constructed from the sheep genotype reference panel under the default heritability model while accounting for local LD structure.

### TWAS of complex traits

To investigate whether the *cis*-genetic components of molecular phenotypes are associated with complex traits in sheep, we performed TWAS based on the 34 meta-GWAS summary statistics. More specifically, we applied single-tissue and multi-tissue TWAS using S-PrediXcan^39^ and S-MultiXcan^40^, respectively, as implemented in the MetaXcan framework (v0.8.0). Briefly, we trained nested cross-validated elastic net models to predict each of the seven types of molecular phenotypes across 51 tissues, using SNPs within ±1□Mb of each gene’s TSS based on the NCBI *Ovis aries* Annotation Release 104 (ARS-UI_Ramb_v2.0, GCF_016772045.1) as predictors and covariate-corrected molecular phenotypes as outcomes. Only predictive models with cross-validated prediction performance exceeding correlation ρ > 0.1 and prediction performance *P* < 0.05 were retained for downstream TWAS analysis. We then applied S-PrediXcan to evaluate gene-trait associations in individual tissues, followed by S-MultiXcan to integrate these results across tissues and identify robust multi-tissue associations. Multiple testing was accounted for using the false discovery rate (FDR) for each trait^40^, with an adjusted *P*□<□0.05 considered statistically significant.

### Colocalization of GWAS and molQTL

To prioritize putative variants, genes, and tissues, we performed systematic colocalization analyses between molQTL and GWAS summary statistics from 34 complex traits. We used the coloc R package (v5.2.3)^107^ to compute posterior probabilities for five hypotheses, focusing on the posterior probability of a shared causal variant (PP_H4_) as evidence of colocalization. We defined a colocalization as significant when PP_H4_□>□0.8. Summary statistics from SNPs within 340 GWAS loci were extracted and matched to the corresponding molQTL summary statistics from the same SNP set. Colocalization analyses were performed independently for each combination of molQTL and tissue. To account for differences in tissue sample size and molGene discovery power, we computed an adjusted proportion of colocalized genes per tissue for prioritizing gene-trait-tissue associations.

### Selective sweep analysis and enrichment of molQTL in selected regions

To identify genomic regions under selection, we calculated the pairwise genetic fixation index (*F*_ST_)^116^ between 710 European (EUR) and 1,104 Central and East Asian (CEA) sheep using 21,542,313 common SNPs (MAF□≥□0.05) from the sheep genotype reference panel. *F*_ST_ was computed in 50-kb sliding windows with a 10-kb step size using VCFtools (v0.1.16)^117^. Genomic intervals falling within the top 1% *F*_ST_ values were considered to be candidate selective sweeps. Adjacent regions were merged using BEDTools (v2.31.0)^101^, and functional annotation of candidate sweeps was performed with ANNOVAR^118^. To support the robustness of selective signals, additional neutrality statistics, including nucleotide diversity (π) and Tajima’s *D*, were also estimated using VCFtools.

To assess the enrichment of regulatory variants in regions under selection, we calculated odds ratio (OR) as:

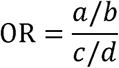

where *a* denotes the number of molQTL in selected regions, *b* denotes the number of non-molQTL SNPs in selected regions, *c* denotes the number of molQTL in non-selected regions, and *d* denotes the number of non-molQTL SNPs in non-selected regions. To ensure that the observed enrichment patterns were driven by the underlying biology of the regulatory variants rather than confounding by local MAF and LD structure, we also performed enrichment analyses using control SNPs matched for both MAF and LD score, following established protocols^119^. Briefly, we used common SNPs (MAF > 0.05) from 2,816 unrelated sheep to compute MAFs and LD scores for all variants using GCTA (v1.94.1)^120^ with default parameters. For each lead or fine-mapped variant (focal SNP), control SNPs were randomly sampled from a pool of 3,346,402 variants (excluding focal SNPs) under two matching criteria: (1) MAF within ±0.02 of the focal SNP; and (2) LD score within ±0.1 standard deviations of the focal SNP.

### Ancient genome data processing

For ancient genome analysis, we compiled 52 previously published ancient sheep genomes (mean coverage ≥ 0.5×) from Europe and Asia^1,54–56^. Single-end reads were adapter-trimmed using Cutadapt (v4.6)^121^, and paired-end reads were processed using AdapterRemoval (v2.3.3)^122^. High-quality clean reads were then aligned to the *Ovis aries* reference genome (ARS-UI_Ramb_v2.0, GCF_016772045.1) using BWA-backtrack (BWA aln) algorithm (v0.7.5a)^123^ with default parameters, and subsequently sorted with SAMtools (v1.18)^90^. PCR duplicates were marked and removed using Picard (v2.21.2) (https://broadinstitute.github.io/picard), and local realignment around indels was performed with GATK (v4.1.4.1). Reads with mapping quality < 30 were excluded using SAMtools. Misincorporation patterns were calculated and visualized using mapDamage (v2.2.2)^124^ (**Supplementary Fig. 32**). Autosomal coverage was estimated using *DepthOfCoverage* function in GATK. Genotype imputation for ancient samples with > 0.5× coverage was performed using GLIMPSE2 (v2.0.0) with the sheep genotype reference panel described above. Imputed genotypes with INFO ≥ 0.80 and MAF ≥ 0.01 were retained for subsequent analyses.

Gene expression of 51 tissues in 52 ancient sheep was predicted using the Predict.py script from the MetaXcan framework (v0.8.0)^40^, based on the trained nested cross-validated elastic net model of eQTL described in **TWAS of complex traits**. Genome-wide genetic differentiation between ancient (> 4,000 YBP) and recent (< 4,000 YBP) domestic sheep was assessed using sliding-window *F*_ST_ (50-kb window, 10-kb step) via VCFtools (v0.1.13). The top 1% of *F*_ST_ windows were defined as candidate selective sweeps and annotated using ANNOVAR. Enrichment of eQTL within these regions was quantified using odds ratio (OR). Gene clustering along time points in ancient samples was performed using Mfuzz (v2.68.0)^125^ in R, based on predicted expression.

### Statistics and reproducibility

No statistical method was used to predetermine sample sizes. Details of data exclusions specific to each analysis are provided in **Methods**. For all boxplots, the center line represents the median; box limits indicate the interquartile range (IQR), from the 25^th^ (Q1) to 75^th^ (Q3) percentile; whiskers extend to the smallest and largest values within 1.5 × IQR from the lower and upper quartiles, respectively. Outliers beyond the whiskers are not shown unless otherwise specified. No randomization was applied, as all datasets were derived from publicly available, observational studies. Investigators were not blinded to sample allocation or outcome assessment, as the data were not generated from randomized controlled experiments.

## Data availability

All the existing data analyzed in this study are publicly available for download without restrictions from the Sequence Read Archive (https://www.ncbi.nlm.nih.gov/sra) database. The raw RNA-seq data newly generated in this study are available in the Sequence Read Archive under accession numbers PRJNA1198671 and PRJNA1304012, while the whole-genome sequencing data can be retrieved under accession number PRJNA1403714. Detailed information on RNA-seq, WGS, molQTL summary, GWAS summary and selection sweep results can be found in Supplementary Tables. All processed data and the full summary statistics of molQTL mapping are available at (https://sheepgtex.farmgtex.org).

## Code availability

All the computational scripts and codes for RNA-seq and WGS analyses, as well as the respective quality control, molecular phenotype normalization, genotype imputation, molQTL mapping, functional enrichment, colocalization, TWAS, selection scans, ancient DNA analyses and figure generation, are publicly available at the FarmGTEx GitHub website (https://github.com/FarmGTEx/SheepGTEx-Pipeline-v0).

## Acknowledgements

This work was supported by grants from the National Key R&D Program of China (2023YFF1001800 and 2022YFF1000100 to Z.P. and Y.J.), the National Natural Science Fund for Excellent Young Scientists Fund Program (Overseas) (2024-HY-01 to Z.P.), the Biological Breeding-National Science and Technology Major Project (2023ZD04051 and 2022ZD04017 to Z.P.), seed-funding from CellFood Hub (AUFF to L.F.), Nanfan Special Program (YBXM2539 to Z.P.), Tianshan Talent Project of Xinjiang Uygur Autonomous Region (2022TSYCCX0045 to W. Wu), the Agricultural Science and Technology Innovation Program of China (ASTIP-IAS13 to M.C.), the China Agriculture Research System of MOF and MARA (CARS-38-02 to M.C.), Strategic Priority Research Program of the Chinese Academy of Sciences (XDA0460405 to Xuemei L., Y.W., J. Lu and M.W.), the National Natural Science Foundation of China (U21A20247 to Y.J.), the Basic Research Project of Yazhouwan National Laboratory (JL23YCKY01 to J. Han), and Central Public-interest Scientific Institution Basal Research Fund (Y2025YC56 to Z.P.).

We thank all the researchers who contributed to the publicly available datasets utilized in this study. We are particularly grateful to the Human GTEx and FarmGTEx Consortia for sharing computational code used in data analysis and figure generation. We also acknowledge the High-Performance Computing resources supported by Northwest A&F University and the Institute of Animal Science, Chinese Academy of Agricultural Sciences.

## Author contributions

L.F., Z.P. and Y.J. conceived and designed the study. Z.P., G. Zhang, Hao L., Z.M., X. Zhang, X. Li, X.D., Y.S., X.H., M. Shan, J.Y. and H.Y. collected samples and generated data. M.G., Z. Zhuang, Xiaoning L., P.F., M. Song, P.S., P.L., G. Zhang and Hao L. performed RNA-seq data processing and bioinformatic analyses. Huanhuan Z. and R.L. conducted whole-genome sequencing analyses. M.G., Z. Zhuang, X.S. and M.T. carried out genotype imputation and molQTL mapping. X.S. and J.S. curated and processed GWAS summary statistics. X.S. performed the integration of molQTL with GWAS. Z. Zhuang, Z.B. and M. Song contributed to cross-species GTEx comparisons. M.G., Z. Zhuang, M. Shan, Yuanyuan Z., Yuwen L., Z.M. and Z.P. conducted validation and functional annotation of molQTL. M.G. led the context-specific molQTL analyses. M.G. and Y.X. performed selective sweep analyses. Y.X. led the analysis of ancient genomes. L.F., D.G., Z.P., Y.J., M.G., J.T., Houcheng L., X. Zhu, Q. Lin, D.Z., B. Lin, X. Liu, Weijie Z., W.G., B.A., Q.Z., X.H., C.L., X.W., B.Z., A.C., R.C., O.M., D.M., J.F.O.G., M.A., G.T., J. Han, F.L., and E.C. contributed to the interpretation of results and manuscript discussions. Y.W., J. Lu, M.W. and Xuemei L. developed the SheepGTEx web portal. D.W., Huicong Z., J. Liu, Z.Y., G.S., M.L., J.G., X.Y., Yanli Z., F.W., Wenxin Z., J. Huang, S.W., J.D., H.T., Z.L., S.F., Yongbin L., Z. Zhang, Y. Zhao, Y. Lin, J. He, J.Z., H.S., Z.W., B. Li, R.X., A.C., W. Wang, J. Han, Q. Liu, J. Li, F.L., Q.Y., G. Zhao, L.J., X. Lan, H. Zhou, G.T., W. Wu, Y.J. and M.C. provided data and computational resources. L.F., M.G., X.S., Y.X. and Huanhuan Z. wrote the manuscript. All authors reviewed, revised and approved the final version.

## Competing interests

The authors declare no competing interests.

## Extended Data Figures and legends

**Extended Data Fig. 1.**
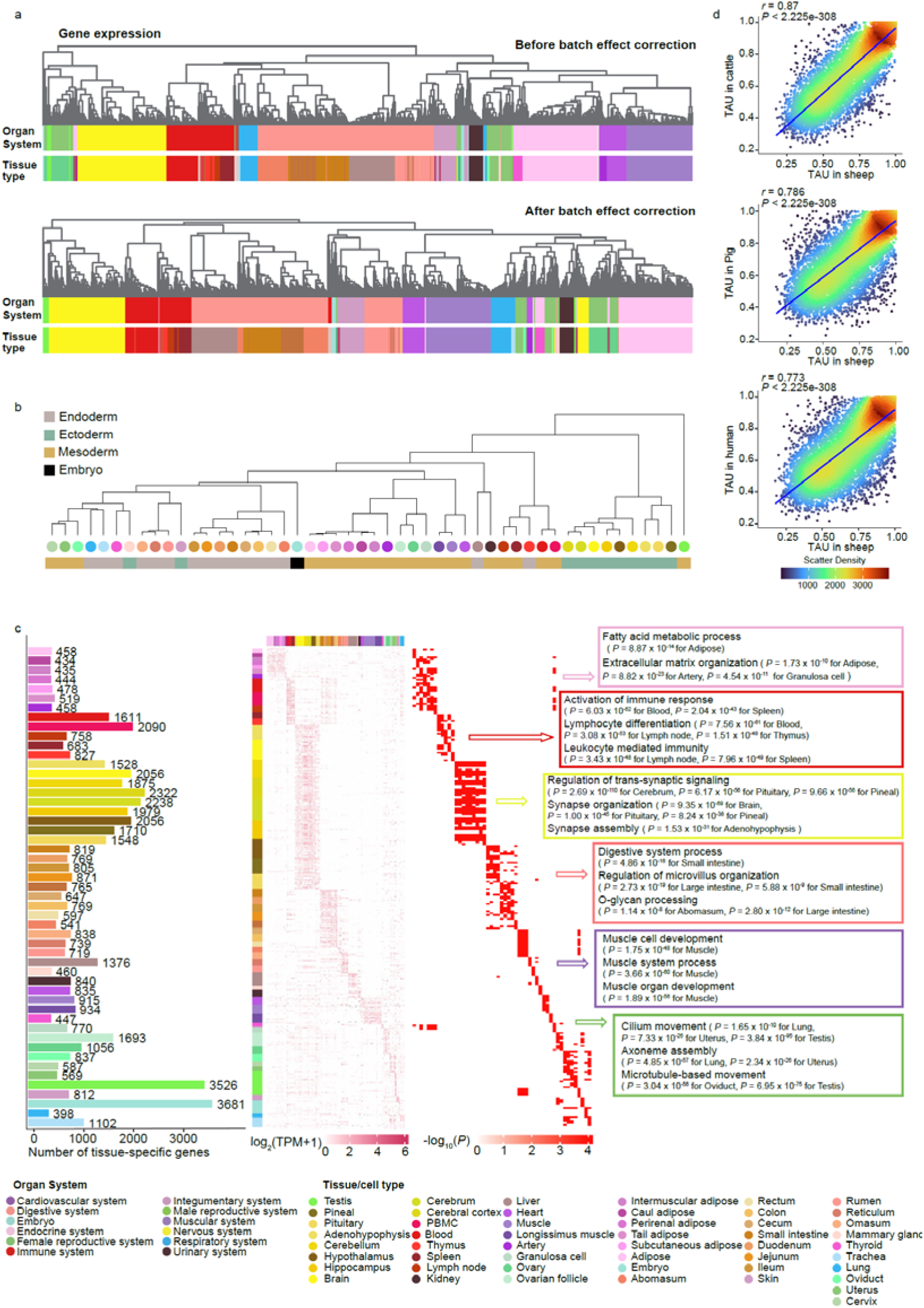
Characterization of gene expression. **a**, Hierarchical clustering of 8,171 RNA-seq samples based on the normalized gene expression (TPM) of the 5,000 most variable genes (∼20% of all genes, ranked by standard deviation), shown before and after batch effect correction. **b**, Phylogeny clustering of 51 tissues based on median gene expression across samples within each tissue, annotated by germ layer origin (endoderm, ectoderm, mesoderm and embryo). **c**, Number of tissue-specific genes (left), and the corresponding expression heatmap across 51 tissues (middle) and enriched Gene Ontology (GO) terms (right). **d**, Pearson’s correlation of tissue specificity (measured by TAU score) between sheep and cattle, pig and human. TPM, transcripts per million.

**Extended Data Fig. 2.**
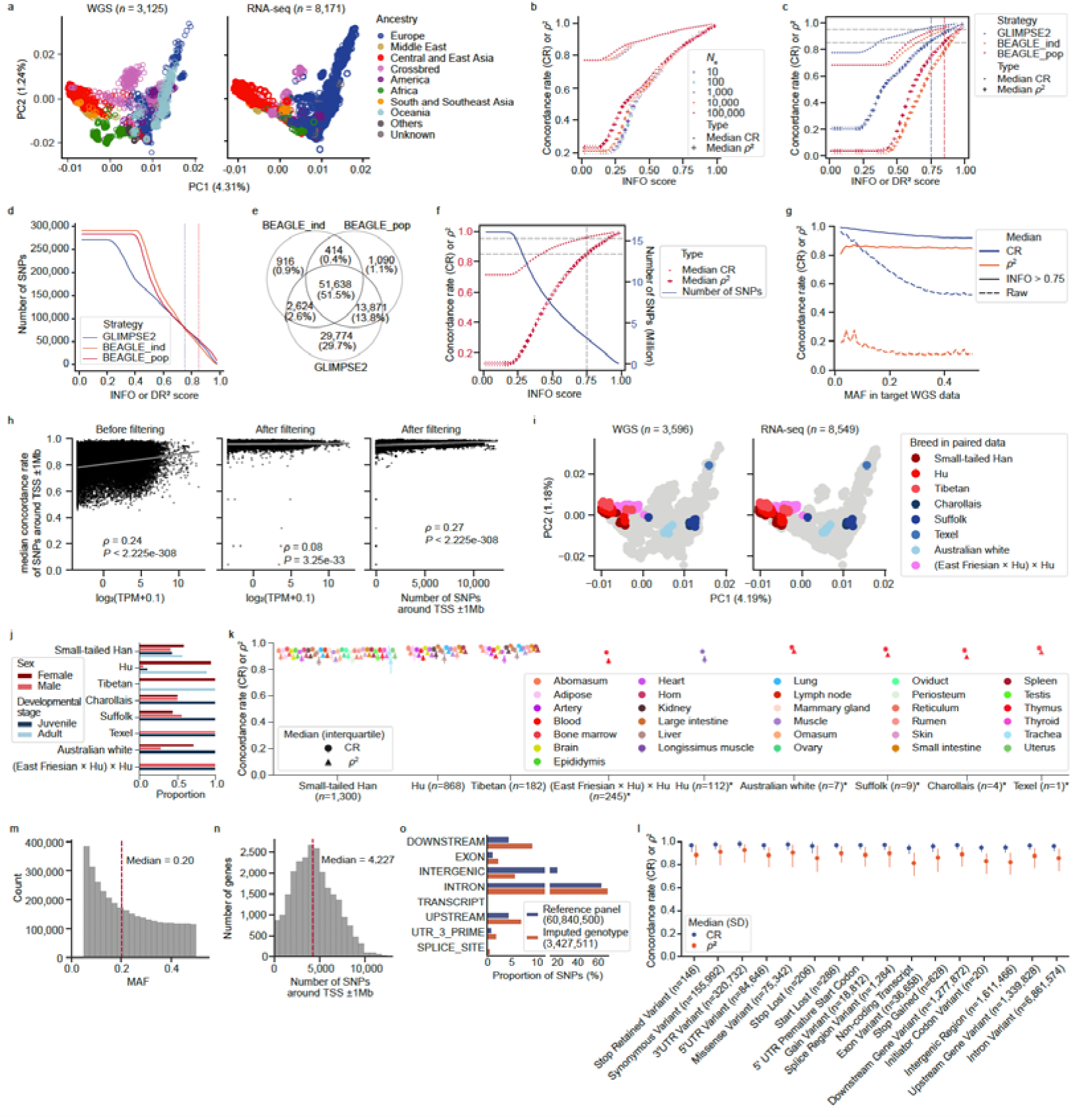
Genotype imputation. **a**, Principal component analysis (PCA) of 3,125 whole-genome sequencing (WGS) samples and 8,171 RNA-seq samples based on 156,560 LD-independent SNPs. The left panel shows WGS samples (*n* = 3,125) from the genotype imputation reference panel, and the right panel shows RNA-seq samples (*n* = 8,171) with successfully imputed genotypes. **b**, Imputation accuracy, measured by concordance rate (CR) and squared Spearman’s correlation (ρ^2^) of genotypes between paired WGS and RNA-seq samples, across different effective population size (*N*_e_) values and INFO thresholds using GLIMPSE2 on chromosome 24 using paired 1,300 RNA-seq samples and 35 WGS Small-tailed Han individuals. **c**, Comparison of imputation accuracy between three imputation strategies: GLIMPSE2, BEAGLE_ind (per-sample), and BEAGLE_pop (population-based). Horizontal lines indicate quality benchmarks (CR ≥ 0.95, ρ^2^ ≥ 0.85). Vertical lines indicate INFO/DR^2^ thresholds that meet these criteria. **d**, Number of SNPs retained under quality controls in **c**. **e**, Venn diagram depicting the overlap of high-quality imputed SNPs (INFO ≥ 0.75 or DR^2^ ≥ 0.85) across strategies. **f**, Effect of INFO thresholds on imputation accuracy (left) and SNP yield (right) on autosomes using GLIMPSE2. **g**, Imputation accuracy across 50 minor allele frequency (MAF) bins before and after INFO filtering. **h**, Spearman’s correlation between CR of *cis*-SNPs [SNPs around□±1□Mb of the transcriptional start site (TSS)] and the corresponding gene expression level, and between CR and the number of high-quality *cis*-SNPs. **i**, PCA of 3,596 WGS samples and 8,549 RNA-seq samples based on 156,560 LD-independent SNPs. The left panel shows WGS samples from the genotype imputation reference panel (*n* = 3,125) and additional individuals with paired RNA-seq data (*n* = 471), while the right panel shows RNA-seq samples with imputed genotypes in discovery data (*n* = 8,171) and validation data independent from the SheepGTEx dataset (*n* = 378). Grey dots denote samples without paired data. **j**, Proportion of sexes and developmental stages across eight breeds of all the paired data. **k**, Genotype imputation accuracy across nine populations, assessed between paired WGS and RNA-seq samples. Asterisks indicate six validation breeds independent of the SheepGTEx discovery cohort. **l**, Imputation accuracy across genomic variant types. **m–n**, MAF distribution and per-gene *cis*-SNP count of 3,427,511 high-quality imputed SNPs in RNA-seq samples. **o**, Distribution of 3,427,511 imputed SNPs in RNA-seq and 60,840,500 SNPs in the reference panel across eight genomic features.

**Extended Data Fig. 3.**
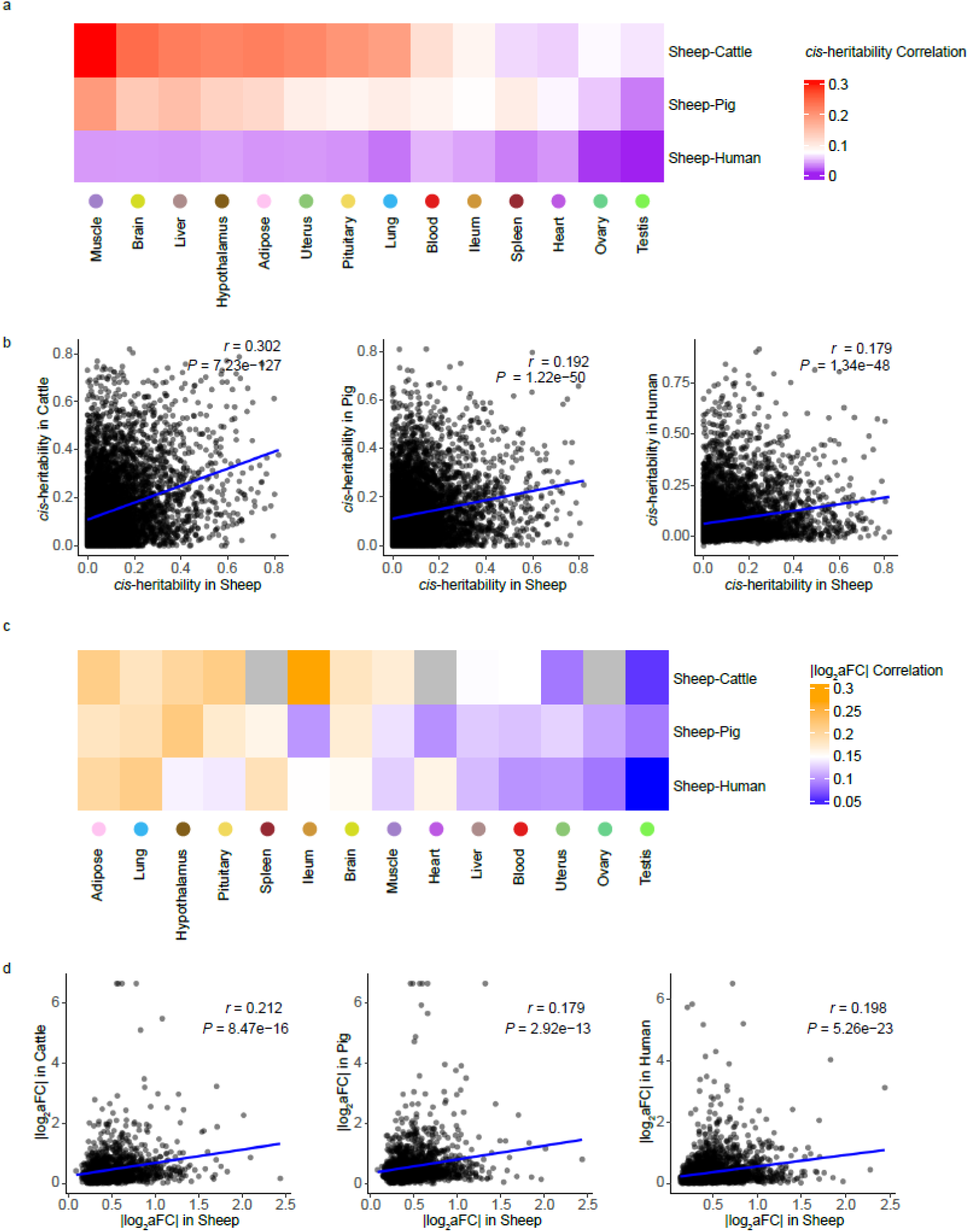
Cross-species comparison of *cis*-heritability and *cis*-eQTL effect sizes. **a–b,** Pearson’s correlation of *cis*-heritability estimates for one-to-one orthologous eGenes between sheep and cattle, sheep and pig, and sheep and human, respectively. **c–d,** Pearson’s correlation of |log_2_aFC| for one-to-one orthologous eGenes between sheep and cattle, pig, and human.

**Extended Data Fig. 4.**
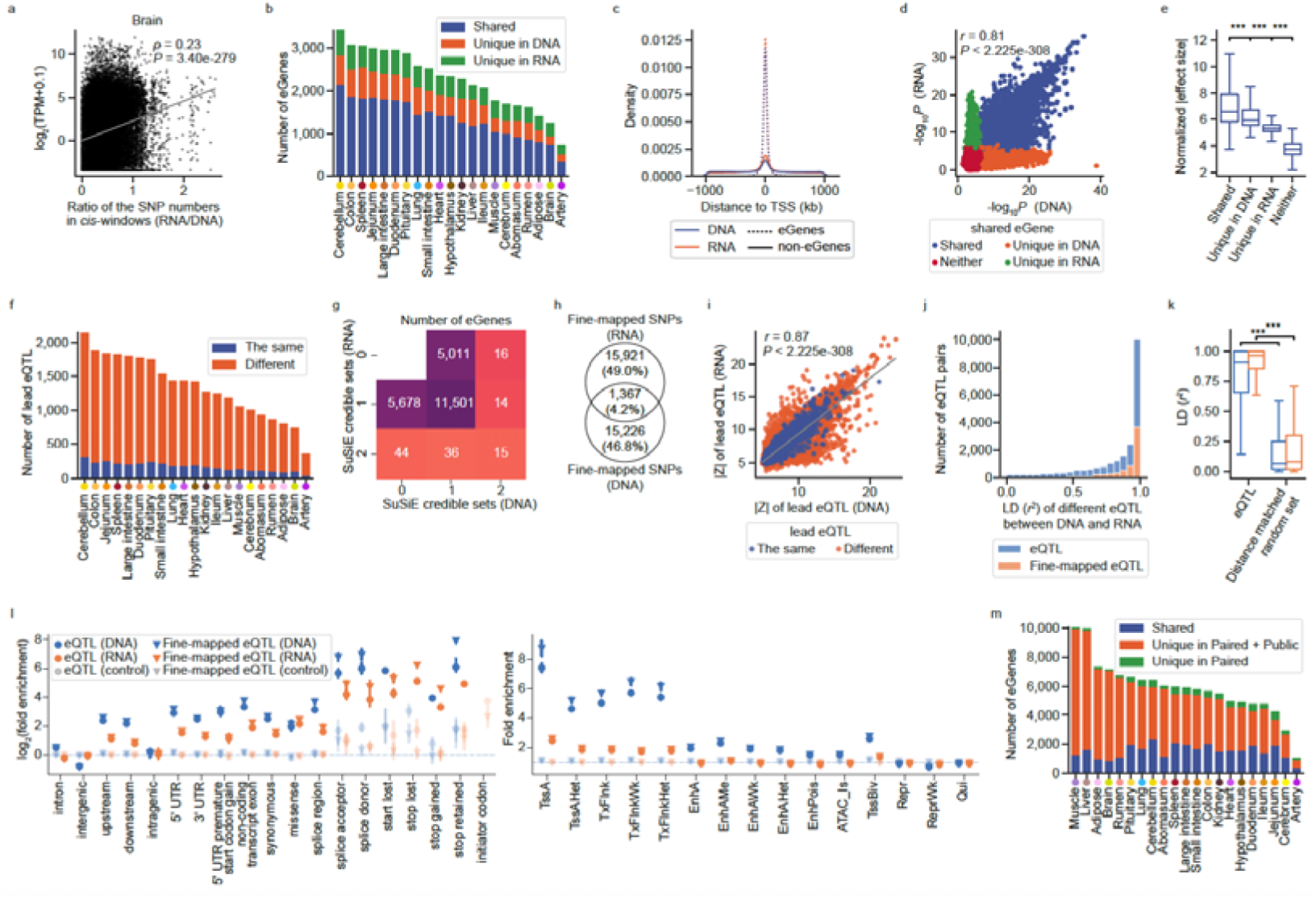
The impact of WGS- and RNA-based genotypes on molQTL detection. **a**, Spearman’s correlation between the ratio of imputed SNPs from RNA-seq data (RNA) to directly observed SNPs from WGS (DNA) within *cis*-windows and the expression levels of the corresponding genes. **b**, Overlap of eGenes identified using DNA-based versus RNA-based genotypes in 93 sheep for which both data types were available. **c**, Genomic distribution of lead variants (relative to TSS) detected using DNA and RNA genotypes, compared between eGenes and non-eGenes. **d**, Pearson’s correlation of significance levels (−log_10_*P*) between lead eQTL of shared eGenes identified using DNA- and RNA-derived genotypes. **e**, Normalized |effect size| (|Z|-scores) of lead eQTL stratified into four categories based on whether the lead variants of target eGenes were shared between RNA- and DNA-based analyses. *P* values were calculated using the two-sided Mann-Whitney U test. ns: non-significant; *: *P* < 0.05; **: *P* < 0.01; ***: *P* < 0.001. **f**, Overlap of lead variants for shared eGenes identified using DNA and RNA genotypes. **g**, Heatmap showing the number of eGenes with varying numbers of fine-mapped credible sets identified using WGS-called genotypes (DNA) and RNA-imputed genotypes (RNA). **h**, Venn diagram illustrating the overlap of lead fine-mapped SNPs for the same eGenes detected using DNA- and RNA-derived genotypes. **i**, Pearson’s correlation of |Z|-scores for lead variants of shared eGenes identified by both methods. “The same”: identical lead eQTL for the same gene. “Different”: different lead eQTL for the same gene. **j–k**, Comparison of linkage disequilibrium (LD; measured as *r*^2^) between lead and fine-mapped eQTL identified by DNA- and RNA-based approaches for the same eGenes. **k** includes a control set of distance-matched random SNP pairs. *P* values were calculated using two-sided Mann-Whitney U tests. ***: *P* < 0.001. **l**, Functional enrichment of lead and fine-mapped (focal) eQTL across 19 sequence features (top) and 15 chromatin states (bottom), comparing WGS-called genotypes with RNA-imputed genotypes. Point, mean; error bar, 95% confidence interval. Lighter-colored points denote control SNPs matched for minor allele frequency (MAF) and LD score. Chromatin state abbreviations: TssA, active promoter; TssAHet, flanking active TSS without ATAC; TxFlnk, transcribed gene; TxFlnkWk, weakly transcribed gene; TxFlnkHet, transcribed without ATAC; EnhA, strong active enhancer; EnhAMe, medium enhancer with ATAC; EnhAWk, weak enhancer; EnhAHet, active enhancer without ATAC (heterochromatic); EnhPois, poised enhancer; ATAC_Is, ATAC island; TssBiv, bivalent/poised TSS; Repr, polycomb-repressed; ReprWk, weakly repressed polycomb; Qui, quiescent. **m,** Overlap of eGenes detected using DNA-based genotype versus RNA-based imputation strategies. DNA-based mapping was limited to 93 paired samples, whereas RNA-based mapping utilized imputed genotypes from thousands of public RNA-seq datasets, significantly boosting molQTL detection power.

**Extended Data Fig. 5.**
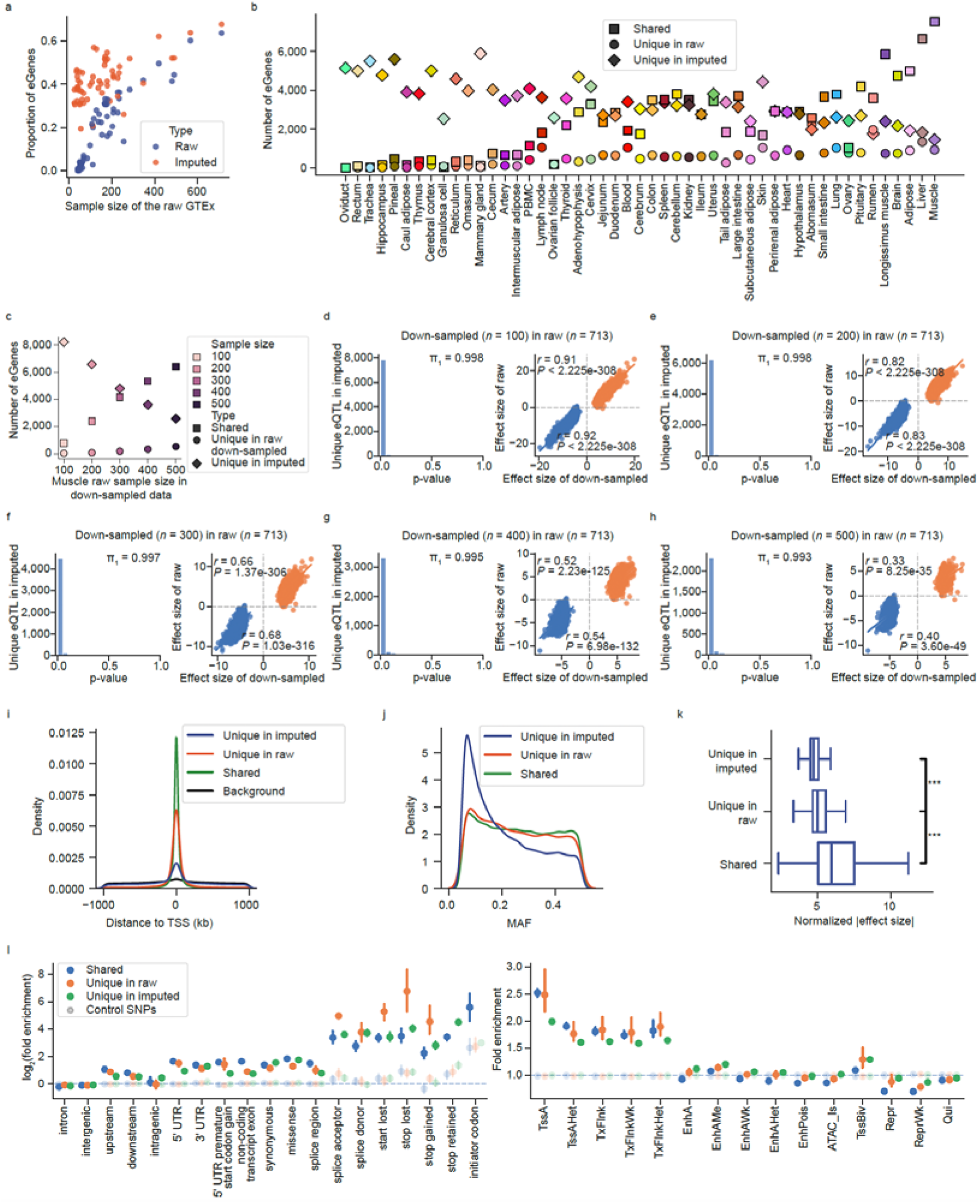
Imputed *cis*-eQTL mapping. **a**, Proportion of eGenes identified using either raw or imputed expression data, shown as a function of raw data sample size. **b**, Number of shared and unique eGenes identified using raw versus imputed expression data across tissues. **c**, Number of shared and unique eGenes identified using raw down-sampled expression data versus imputed expression data in muscle. For the imputed dataset, gene expression imputation models were trained using down-sampled muscle cohorts (*n* = 100–500) and subsequently applied to all available samples (*n* = 1,614) for downstream *cis*-eQTL mapping. **d–h**, Validation of eQTL uniquely discovered in imputed dataset compared to different down-sampled raw data. Storey’s π_1_ statistics and Spearman’s correlation of normalized effect sizeswere used to assess replication of unique eQTL in the “gold standard” full dataset (*n* = 713). **i**, Genomic distribution of lead variants relative to TSS, compared between eGenes shared by both methods and those uniquely identified by one method. **j**, MAF distribution of lead variants, stratified by eGenes shared or unique to each method. **k**, Normalized absolute effect sizes (|Z|-scores) of lead eQTL, compared between shared and method-specific eGenes. *P* values were calculated using the two-sided Mann-Whitney U test. ns: non-significant; *: *P* < 0.05; **: *P* < 0.01; ***: *P* < 0.001. **l**, Enrichment of eQTL in 19 sequence features (top) and 15 chromatin states (bottom), analyzed separately for shared and unique eGenes. Point, mean; error bar, 95% confidence interval. Lighter-colored points indicate control SNPs matched for minor allele frequency (MAF) and linkage disequilibrium (LD) score. Chromatin state abbreviations: TssA, active promoter; TssAHet, flanking active TSS without ATAC; TxFlnk, transcribed gene; TxFlnkWk, weakly transcribed gene; TxFlnkHet, transcribed without ATAC; EnhA, strong active enhancer; EnhAMe, medium enhancer with ATAC; EnhAWk, weak enhancer; EnhAHet, active enhancer without ATAC (heterochromatic); EnhPois, poised enhancer; ATAC_Is, ATAC island; TssBiv, bivalent/poised TSS; Repr, polycomb-repressed; ReprWk, weakly repressed polycomb; Qui, quiescent.

**Extended Data Fig. 6.**
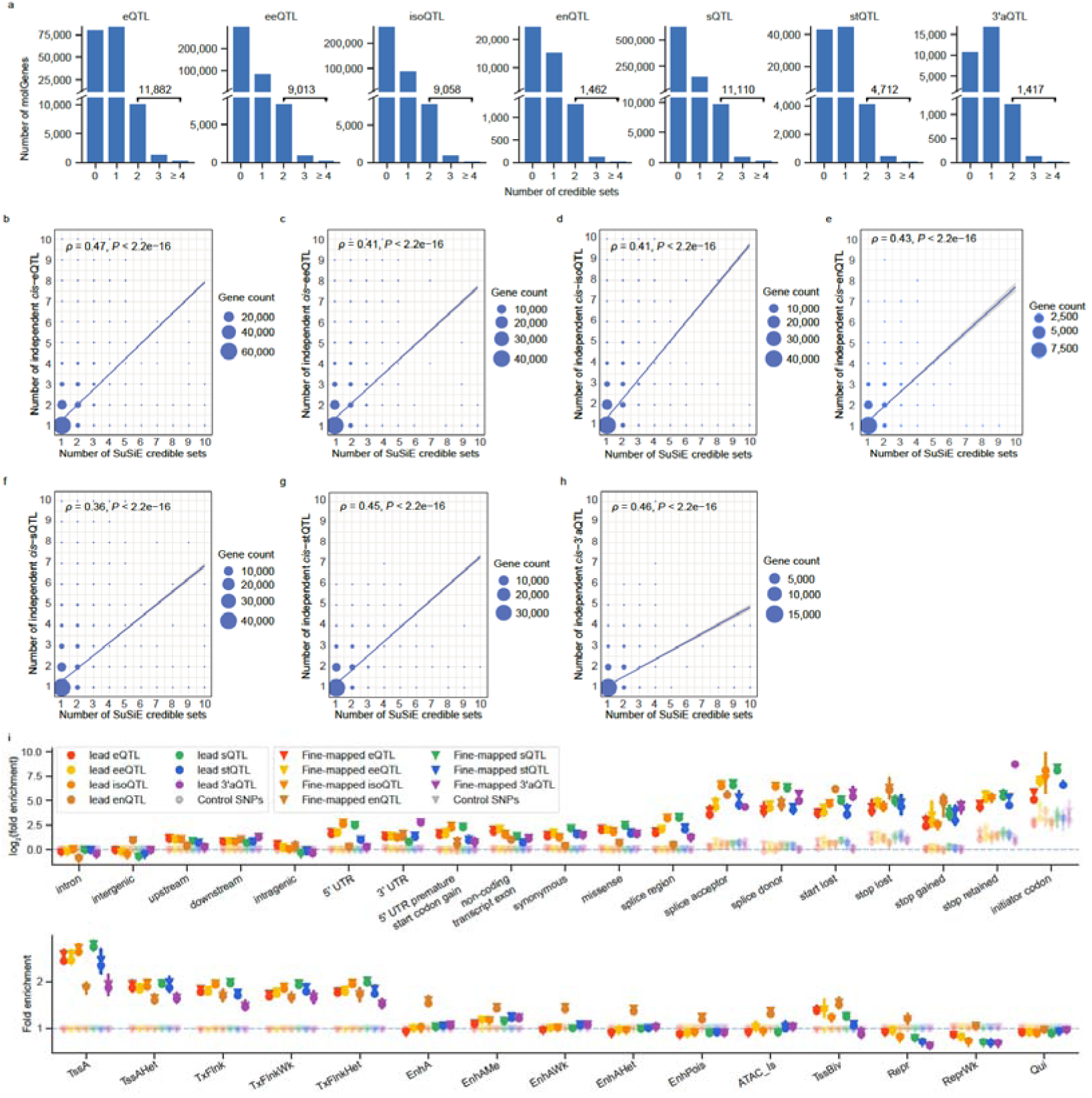
Fine-mapped *cis*-molQTL. **a**, Distribution of molGenes by the number of fine-mapped credible sets. **b–h**, Spearman’s correlation between the number of fine-mapped credible sets and the number of conditionally independent molQTL per gene. Dot size reflects the number of genes. **i**, Enrichment (mean ±95% confidence interval) of seven types of lead and fine-mapped *cis*-molQTL across 19 sequence features (top) and 15 chromatin states (bottom). Lighter-colored points denote control SNPs matched for minor allele frequency (MAF) and linkage disequilibrium (LD) score. Chromatin state abbreviations: TssA, active promoter; TssAHet, flanking active TSS without ATAC; TxFlnk, transcribed gene; TxFlnkWk, weakly transcribed gene; TxFlnkHet, transcribed without ATAC; EnhA, strong active enhancer; EnhAMe, medium enhancer with ATAC; EnhAWk, weak enhancer; EnhAHet, active enhancer without ATAC (heterochromatic); EnhPois, poised enhancer; ATAC_Is, ATAC island; TssBiv, bivalent/poised TSS; Repr, polycomb-repressed; ReprWk, weakly repressed polycomb; Qui, quiescent.

**Extended Data Fig. 7.**
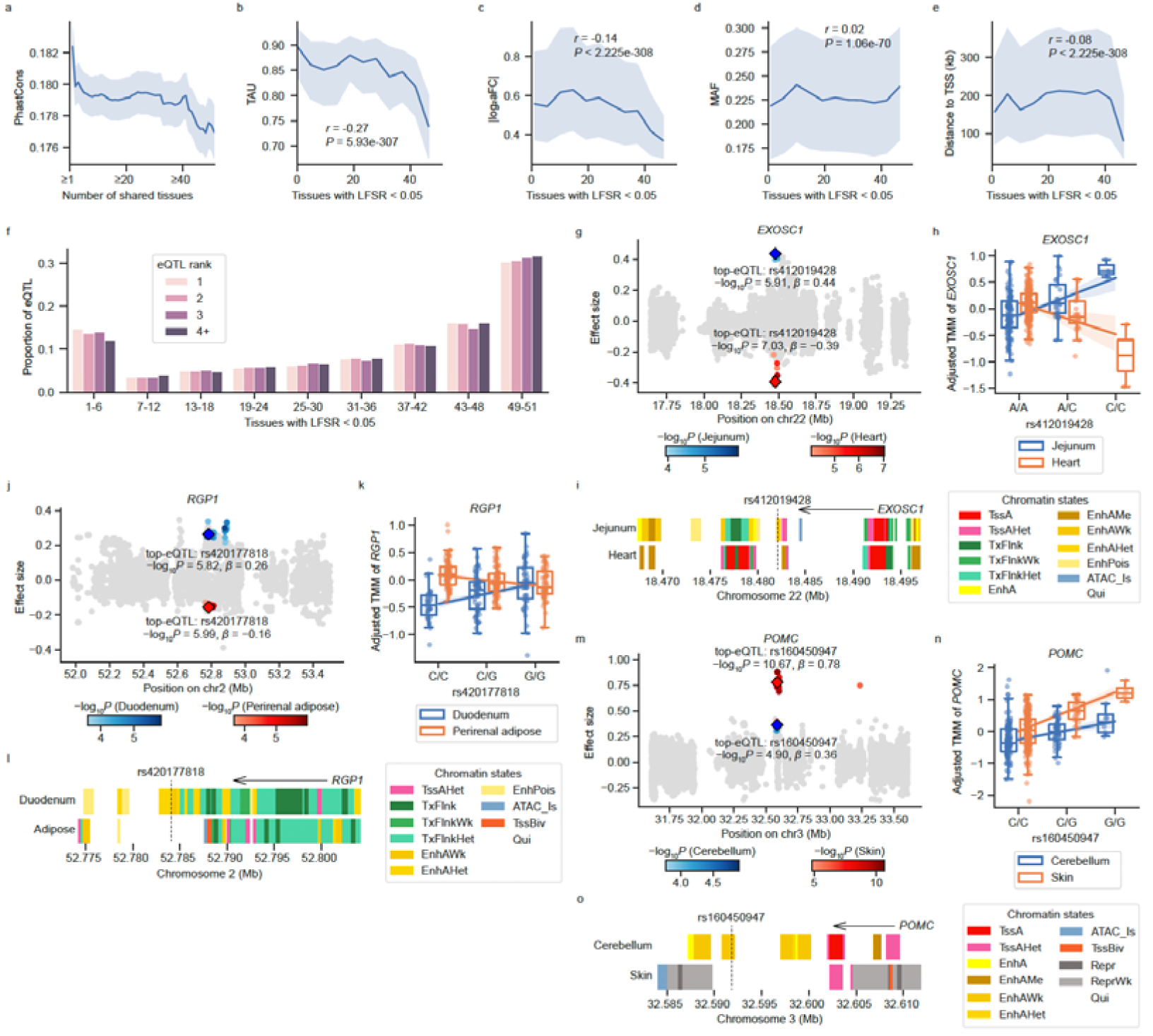
Tissue sharing and specificity properties of *cis*-eQTL. **a**, Sequence conservation (PhastCons score) of eGenes as a function of tissue sharing. The line and shading indicate the mean and standard error, respectively. **b–e**, Relationships between the number of tissues sharing active *cis*-eQTL and their tissue specificity (TAU scores) (**b**) effect sizes (|log_(aFC)|) (**c**), MAF (**d**), distances to the TSS (**e**) and for each gene. **f**, Proportion of primary and non-primary *cis*-eQTL active across different numbers of tissues. **g–i**, *EXOSC1* with opposite eQTL (rs412019428) effects between jejunum and heart. **j–l**, *RGP1* with opposite eQTL (rs420177818) effects betw een duodenum and perirenal adipose. **m–o**, *POMC* with heterogeneous eQTL (rs160450947) effects between cerebellum and skin. Chromatin state abbreviations: TssA, active promoter; TssAHet, flanking active TSS without ATAC; TxFlnk, transcribed gene; TxFlnkWk, weakly transcribed gene; TxFlnkHet, transcribed without ATAC; EnhA, strong active enhancer; EnhAMe, medium enhancer with ATAC; EnhAWk, weak enhancer; EnhAHet, active enhancer without ATAC (heterochromatic); EnhPois, poised enhancer; ATAC_Is, ATAC island; TssBiv, bivalent/poised TSS; Repr, polycomb-repressed; ReprWk, weakly repressed polycomb; Qui, quiescent.

**Extended Data Fig. 8.**
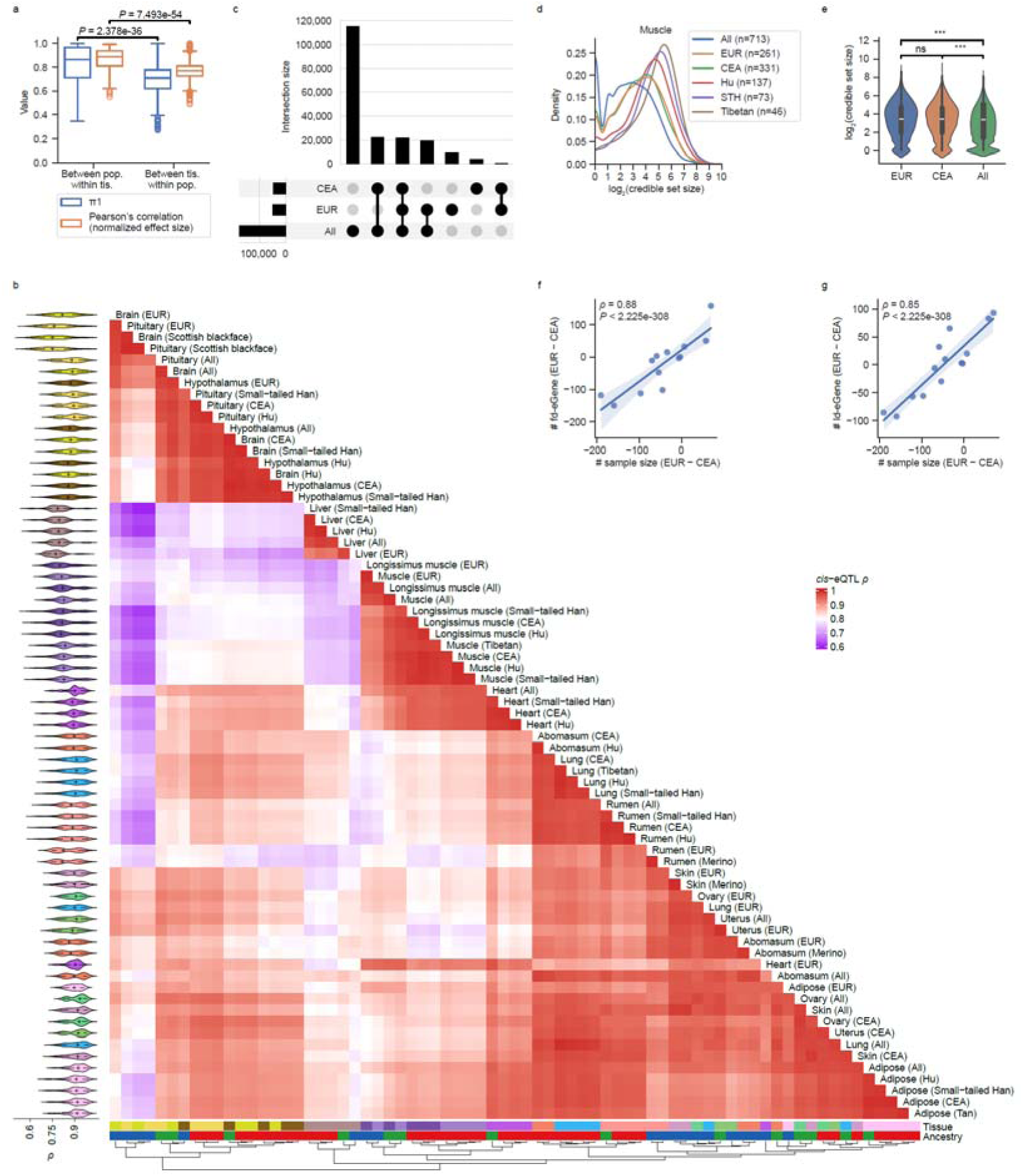
Breed-sharing and specificity of molQTL. **a**, Story’s π_1_ replication statistics and Pearson’s correlations of *cis*-eQTL effect sizes, calculated either between populations within a tissue or between tissues within a population. Populations include all samples (All), Europe (EUR), Central and East Asia (CEA), and breeds with sufficient sample size for QTL mapping (n ≥ 40 per tissue). *P* values are derived from Mann-Whitney U tests. **b**, Clustered heatmap showing pairwise Spearman’s correlation (ρ) of *cis*-eQTL effect sizes across tissues and populations. **c**, Overlap of eGenes identified in EUR, CEA, and all samples across 14 tissues. **d**, Comparison of fine-mapping credible set sizes between populations in muscle. **e**, Violin plot showing significant differences in credible set sizes between ancestry groups, assessed by the Mann-Whitney U test. **f–g**, Number of fd-eGene (**f**) and ld-eGene (**g**) as a function of the sample size differences (EUR minus CEA) across tissues, evaluated using Spearman’s ρ.

**Extended Data Fig. 9.**
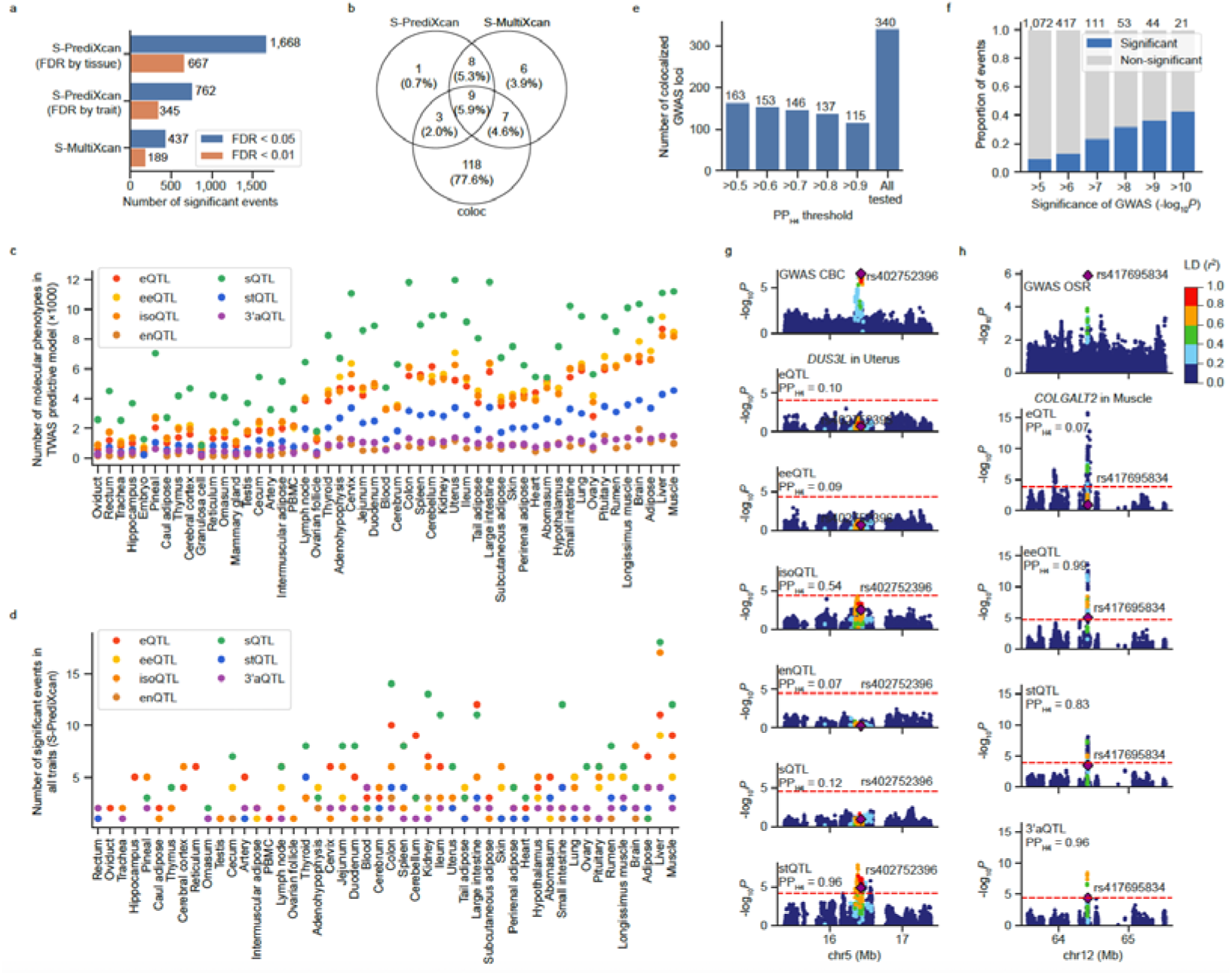
Interpretation of GWAS loci with molQTL. **a**, Number of significant events (trait-tissue-molecular phenotype associations) across different false discovery rate (FDR) thresholds in transcriptome-wide association studies (TWAS), including single-tissue TWAS (S-PrediXcan), and multi-tissue TWAS (S-MultiXcan). **b**, Venn diagram showing the number of GWAS loci linked to at least one *cis*-molQTL by three complementary approaches: colocalization analysis (coloc), S-PrediXcan, and S-MultiXcan. **c**, Number of molecular phenotypes retained in nested cross-validated elastic net prediction models after filtering across tissues and *cis*-molQTL types, ordered by increasing sample size. **d**, Number of molecular phenotypes significantly associated with at least one trait in S-PrediXcan across tissues and cis-molQTL classes, ordered by increasing sample size. **e**, Number of colocalized GWAS loci supported by at least one *cis*-molQTL in at least one tissue under different posterior probability of colocalization (PP_H4_) thresholds. **f**, Proportion of significant colocalizations (PP_H4_ > 0.8) between GWAS loci and *cis*-molQTL across different GWAS significance thresholds. The total number of tested colocalizations at each significance threshold is indicated above each bar. **g,** Cannon bone circumference (CBC) shows colocalization with stQTL of *DUS3L* gene (PP_H4_ = 0.96) at rs402752396 in uterus. **h,** Oil/suint ratio (OSR) shows colocalization with eeQTL (PP_H4_= 0.99) and 3’aQTL (PP_H4_ = 0.96) of *COLGALT2* gene at rs417695834 in muscle.

**Extended Data Fig. 10.**
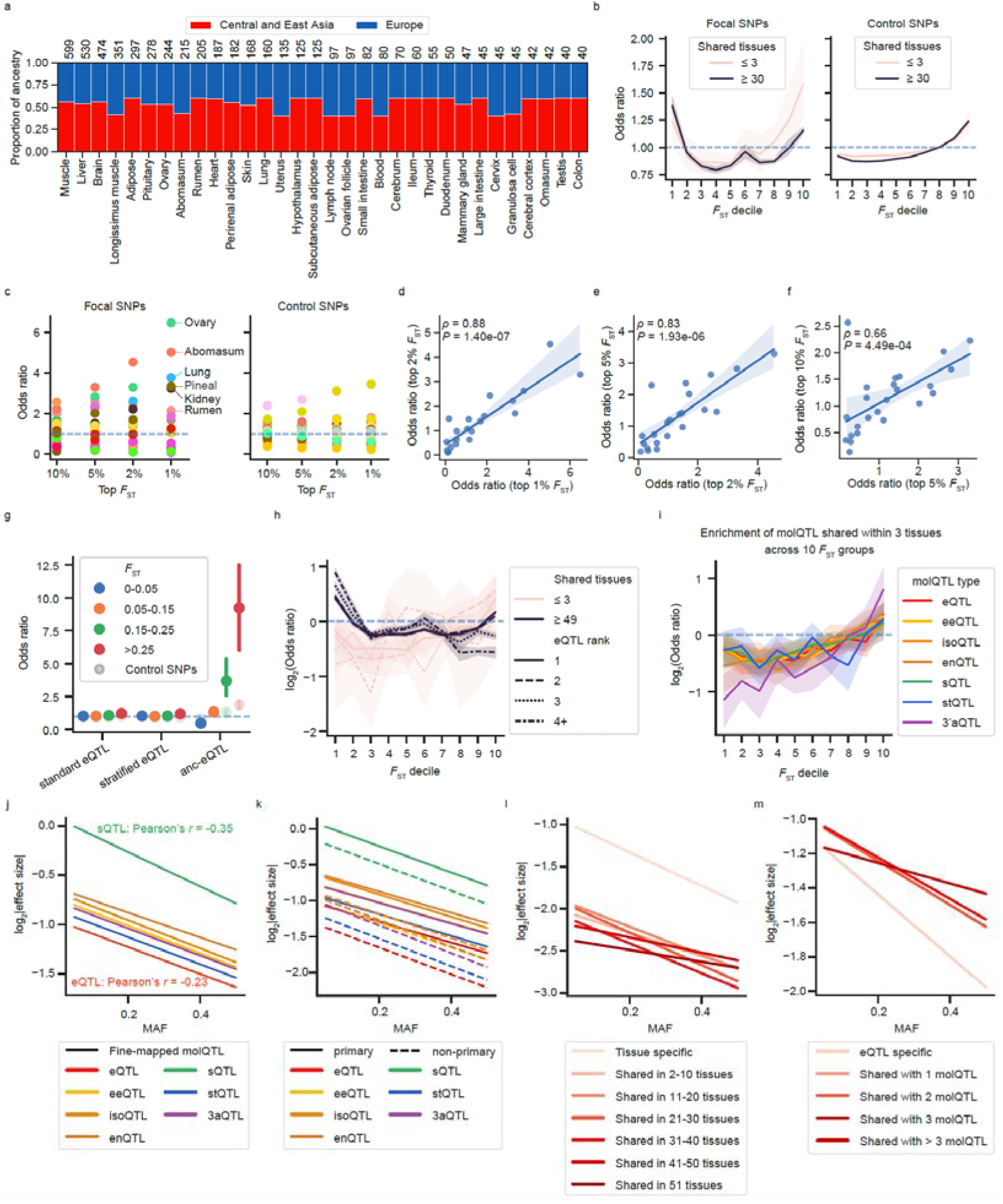
Characterization of molQTL in selection signals. **a**, Proportions of sheep ancestries inferred from geographic origin across 32 tissues, with European and Central/East Asian ancestries balanced (maintaining a ratio between 2:3 and 3:2). Sample sizes for each tissue are indicated above the bars. **b**, Enrichment of *cis*-eQTL shared across ≤ 3 tissues or ≥ 30 tissues across genomic regions stratified by *F*_ST_ deciles, using balanced ancestry groups shown in **a**. Focal *cis*-eQTL SNPs are shown on the left, with control SNPs matched for minor allele frequency (MAF) and linkage disequilibrium (LD) score on the right. **c**, Enrichment of tissue-specific eQTL in different sets of selection signals. Focal *cis*-eQTL SNPs are shown on the left, with MAF- and LD-matched control SNPs on the right. **d–f**, Spearman’s correlations of tissue-specific eQTL enrichment between selection thresholds: top 1% vs. top 2% *F*_ST_ (**d**), top 2% vs. top 5% (**e**), and top 5% vs. top 10% (**f**). **g**, Enrichment of standard eQTL (all samples), stratified eQTL (EUR or CEA ancestry groups), and anc-eQTL in genomic regions under selection. Lighter-colored points indicate control SNPs matched for MAF and LD score. **h**, Enrichment of eQTL shared across ≤ 3 tissues or ≥ 49 tissues in *F*_ST_-ranked genomic deciles, evaluated across different independent eQTL ranks. **i**, Enrichment of molQTL shared within three tissues in *F*_ST_-ranked genomic deciles, across different molQTL types. **j–m**, Selection pressure, as measured by the relationship between MAF and effect size, across fine-mapped molQTL (**j**), primary versus non-primary molQTL (**k**), tissue-sharing patterns of eQTL (**l**), and pleiotropy of eQTL (**m**). Effect sizes were quantified as slopes estimated by TensorQTL (v1.0.9)^25^.

## Notes

https://sheepgtex.farmgtex.org/

### Competing Interest Statement

The authors have declared no competing interest.

### Summary of Updates

we have performed a comprehensive set of additional analyses that have significantly strengthened the rigor and biological insights of our study. Key revisions include: 1) expansion of functional interpretation of molQTL findings by integrating regulatory elements (e.g., enhancer and promoter), and 2) Enhanced methodological rigor particularly for integrative analysis of molQTL and GWAS.

## References

1. Daly, K.G. et al. Ancient genomics and the origin, dispersal, and development of domestic sheep. Science 387, 492–497 (2025).

2. Lv, F.-H. et al. Whole-genome resequencing of worldwide wild and domestic sheep elucidates genetic diversity, introgression and agronomically important loci. Molecular Biology and Evolution (2021).

3. Banstola, A. & Reynolds, J.N.J. The Sheep as a Large Animal Model for the Investigation and Treatment of Human Disorders. Biology (Basel) 11(2022).

4. Yan, Z. et al. A time-resolved multi-omics atlas of transcriptional regulation in response to high-altitude hypoxia across whole-body tissues. Nature Communications 15, 3970 (2024).

5. Hu, Z.-L., Park, C.A. & Reecy, J.M. Bringing the Animal QTLdb and CorrDB into the future: meeting new challenges and providing updated services. Nucleic Acids Research 50, D956–D961 (2021).

6. Albert, F.W. & Kruglyak, L. The role of regulatory variation in complex traits and disease. Nature Reviews Genetics 16, 197–212 (2015).

7. Oliva, M. et al. The impact of sex on gene expression across human tissues. Science 369(2020).

8. Coorens, T.H.H. et al. The human and non-human primate developmental GTEx projects. Nature 637, 557–564 (2025).

9. The GTEx Consortium atlas of genetic regulatory effects across human tissues. Science 369, 1318–1330 (2020).

10. Fang, L. et al. The Farm Animal Genotype–Tissue Expression (FarmGTEx) Project. Nature Genetics 57, 786–796 (2025).

11. Liu, S. et al. A multi-tissue atlas of regulatory variants in cattle. Nature Genetics (2022).

12. Teng, J. et al. A compendium of genetic regulatory effects across pig tissues. Nature Genetics (2024).

13. Guan, D. et al. Genetic regulation of gene expression across multiple tissues in chickens. Nature Genetics (2025).

14. Li, H. et al. An atlas of cell type specific regulatory effects in cattle. bioRxiv, 2025.06.23.661035 (2025).

15. Yuan, Z. et al. Expression quantitative trait loci in sheep liver and muscle contribute to variations in meat traits. Genetics Selection Evolution 53, 8 (2021).

16. Zeng, B. et al. Multi-ancestry eQTL meta-analysis of human brain identifies candidate causal variants for brain-related traits. Nature Genetics 54, 161–169 (2022).

17. Orchard, P. et al. Cross-cohort analysis of expression and splicing quantitative trait loci in TOPMed. medRxiv, 2025.02.19.25322561 (2025).

18. Alkallas, R., Fish, L., Goodarzi, H. & Najafabadi, H.S. Inference of RNA decay rate from transcriptional profiling highlights the regulatory programs of Alzheimer’s disease. Nature Communications 8, 909 (2017).

19. Li, L. et al. An atlas of alternative polyadenylation quantitative trait loci contributing to complex trait and disease heritability. Nat Genet 53, 994–1005 (2021).

20. Li, Y.I. et al. RNA splicing is a primary link between genetic variation and disease. Science 352, 600–604 (2016).

21. Qi, T. et al. Genetic control of RNA splicing and its distinct role in complex trait variation. Nature Genetics 54, 1355–1363 (2022).

22. Li, Y., Ge, X., Peng, F., Li, W. & Li, J.J. Exaggerated false positives by popular differential expression methods when analyzing human population samples. Genome Biology 23, 79 (2022).

23. Lenffer, J. et al. OMIA (Online Mendelian Inheritance in Animals): an enhanced platform and integration into the Entrez search interface at NCBI. Nucleic Acids Res 34, D599–601 (2006).

24. Davenport, K.M. et al. An improved ovine reference genome assembly to facilitate in-depth functional annotation of the sheep genome. Gigascience 11(2022).

25. Taylor-Weiner, A. et al. Scaling computational genomics to millions of individuals with GPUs. Genome Biology 20, 228 (2019).

26. Antonio P. Camargo, Adrielle A. Vasconcelos, Mateus B. Fiamenghi, Gonçalo A. G. Pereira & Marcelo F. Carazzolle. tspex: a tissue-specificity calculator for gene expression data. Research Square (2020).

27. Langfelder, P. & Horvath, S. WGCNA: an R package for weighted correlation network analysis. BMC Bioinformatics 9, 559 (2008).

28. Viñas, R. et al. Hypergraph factorization for multi-tissue gene expression imputation. Nature Machine Intelligence 5, 739–753 (2023).

29. Shen, W.K. et al. AnimalTFDB 4.0: a comprehensive animal transcription factor database updated with variation and expression annotations. Nucleic Acids Res 51, D39–d45 (2023).

30. Brotman, S.M. et al. Adipose tissue eQTL meta-analysis highlights the contribution of allelic heterogeneity to gene expression regulation and cardiometabolic traits. Nature Genetics (2025).

31. Li, H. et al. The enhanced multi-tissue atlas of regulatory effects in cattle. bioRxiv, 2026.03.18.712441 (2026).

32. Chen, L. et al. Large-scale ruminant genome sequencing provides insights into their evolution and distinct traits. Science 364, eaav6202 (2019).

33. Han, B. et al. A multi-tissue single-cell expression atlas in cattle. Nature Genetics (2025).

34. Mizuno, A. & Okada, Y. Biological characterization of expression quantitative trait loci (eQTLs) showing tissue-specific opposite directional effects. European Journal of Human Genetics 27, 1745–1756 (2019).

35. Hensel, K.O. et al. Differential Expression of Mucosal Trefoil Factors and Mucins in Pediatric Inflammatory Bowel Diseases. Scientific Reports 4, 7343 (2014).

36. Kachuri, L. et al. Gene expression in African Americans, Puerto Ricans and Mexican Americans reveals ancestry-specific patterns of genetic architecture. Nature Genetics 55, 952–963 (2023).

37. Haslin, E. et al. Genome-Wide Association Studies of Live Weight at First Breeding at Eight Months of Age and Pregnancy Status of Ewe Lambs. Genes (Basel) 14(2023).

38. Lu, Z. et al. Genome-Wide Association Study of Body Weight Traits in Chinese Fine-Wool Sheep. Animals (Basel) 10(2020).

39. Barbeira, A.N. et al. Exploring the phenotypic consequences of tissue specific gene expression variation inferred from GWAS summary statistics. Nat Commun 9, 1825 (2018).

40. Barbeira, A.N. et al. Integrating predicted transcriptome from multiple tissues improves association detection. PLoS Genet 15, e1007889 (2019).

41. Scott, I.S., Chattopadhyay, A. & Ansorge, O. Development and Microscopic Anatomy of the Pituitary Gland. in Endotext (eds. Feingold, K.R. et al.) (MDText.com, Inc. Copyright © 2000-2025, MDText.com, Inc., South Dartmouth (MA), 2000).

42. Song, C. et al. Molecular cloning, spatial and temporal expression analysis of CatSper genes in the Chinese Meishan pigs. Reprod Biol Endocrinol 9, 132 (2011).

43. Garcia-Rudaz, C. et al. FXYD1, a modulator of Na,K-ATPase activity, facilitates female sexual development by maintaining gonadotrophin-releasing hormone neuronal excitability. J Neuroendocrinol 21, 108–22 (2009).

44. Yang, J. et al. Collagen β(1-O) galactosyltransferase 2 deficiency contributes to lipodystrophy and aggravates NAFLD related to HMW adiponectin in mice. Metabolism 120, 154777 (2021).

45. Tvaroška, I. Glycosylation Modulates the Structure and Functions of Collagen: A Review. Molecules 29(2024).

46. Kalds, P. et al. Genetics of the phenotypic evolution in sheep: a molecular look at diversity-driving genes. Genetics Selection Evolution 54, 61 (2022).

47. Song-Song XU, M.-H.L. Recent advances in understanding genetic variants associated with economically important traits in sheep (*Ovis aries*) revealed by high-throughput screening technologies. Front. Agr. Sci. Eng. 4, 279–288 (2017).

48. Meurens, F. et al. Expression of mucosal chemokines TECK/CCL25 and MEC/CCL28 during fetal development of the ovine mucosal immune system. Immunology 120, 544–55 (2007).

49. Etchevers, L. et al. Exogenous ACTH stimulus during the preovulatory period alters patterns of leukocyte recruitment in the ovary of dairy cows. Theriogenology 195, 176–186 (2023).

50. Hao, Y. et al. Relationship between CCL25/CCR9 Levels in Follicular Fluid and High Ovarian Response in Patients with Polycystic Ovary Syndrome. Int J Endocrinol 2024, 2449037 (2024).

51. Gaspa, G., Cesarani, A., Pauciullo, A., Peana, I. & Macciotta, N.P.P. Genomic Analysis of Sarda Sheep Raised at Diverse Temperatures Highlights Several Genes Involved in Adaptations to the Environment and Heat Stress Response. Animals (Basel) 14(2024).

52. Mei, X. et al. RIPK1 regulates starvation resistance by modulating aspartate catabolism. Nat Commun 12, 6144 (2021).

53. Sohrabi, Y. & Reinecke, H. RIPK1 targeting protects against obesity and atherosclerosis. Trends Endocrinol Metab 32, 420–422 (2021).

54. Larsson, M.N.A. et al. Ancient Sheep Genomes Reveal Four Millennia of North European Short-Tailed Sheep in the Baltic Sea Region. Genome Biol Evol 16(2024).

55. Rossi, C. et al. Exceptional ancient DNA preservation and fibre remains of a Sasanian saltmine sheep mummy in Chehrābād, Iran. Biol Lett 17, 20210222 (2021).

56. Kaptan, D. et al. The Population History of Domestic Sheep Revealed by Paleogenomes. Mol Biol Evol 41(2024).

57. Li, X. et al. Whole-genome resequencing of wild and domestic sheep identifies genes associated with morphological and agronomic traits. Nature Communications 11(2020).

58. Han, Z.P. et al. Population structure and selection signal analysis of indigenous sheep from the southern edge of the Taklamakan Desert. BMC Genomics 25, 681 (2024).

59. Bolormaa, S. et al. A conditional multi-trait sequence GWAS discovers pleiotropic candidate genes and variants for sheep wool, skin wrinkle and breech cover traits. Genetics Selection Evolution 53, 58 (2021).

60. Wu, D. et al. Identification and Expression Patterns of Critical Genes Related to Coat Color in Cashmere Goats. Genes (Basel) 16(2025).

61. Liu, S., Wang, S., Cui, H. & Chen, G. Genome-Wide Association Study on Chinese Merino Sheep Alopecia. Pakistan Journal of Zoology 56, 87 (2024).

62. Qiao, G. et al. Genetic Basis of Dorper Sheep (Ovis aries) Revealed by Long-Read De Novo Genome Assembly. Front Genet 13, 846449 (2022).

63. Reissmann, M. & Ludwig, A. Pleiotropic effects of coat colour-associated mutations in humans, mice and other mammals. Semin Cell Dev Biol 24, 576–86 (2013).

64. Wilkins, A.S., Wrangham, R.W. & Fitch, W.T. The “Domestication Syndrome” in Mammals: A Unified Explanation Based on Neural Crest Cell Behavior and Genetics. Genetics 197, 795–808 (2014).

65. Taylor, D.J. et al. Sources of gene expression variation in a globally diverse human cohort. Nature 632, 122–130 (2024).

66. Liu, S. et al. Adaptive Selection of Cis-regulatory Elements in the Han Chinese. Molecular Biology and Evolution 41(2024).

67. Jagoda, E., et al. Detection of Neanderthal Adaptively Introgressed Genetic Variants That Modulate Reporter Gene Expression in Human Immune Cells. Molecular Biology and Evolution 39(2021).

68. Rinker, D.C. et al. Neanderthal introgression reintroduced functional ancestral alleles lost in Eurasian populations. Nature Ecology & Evolution 4, 1332–1341 (2020).

69. Luo, L.Y. et al. Telomere-to-telomere sheep genome assembly identifies variants associated with wool fineness. Nat Genet 57, 218–230 (2025).

70. Li, R. et al. A sheep pangenome reveals the spectrum of structural variations and their effects on tail phenotypes. Genome Research 33, 463–477 (2023).

71. Clark, E.L. et al. A high resolution atlas of gene expression in the domestic sheep (Ovis aries). PLoS Genet 13, e1006997 (2017).

72. Zhao, B. et al. A Developmental Gene Expression Atlas Reveals Novel Biological Basis of Complex Phenotypes in Sheep. Genomics Proteomics Bioinformatics (2025).

73. Ran, F.A. et al. Genome engineering using the CRISPR-Cas9 system. Nature Protocols 8, 2281–2308 (2013).

74. Liu, S. et al. Systematic identification of regulatory variants associated with cancer risk. Genome Biology 18, 194 (2017).

75. Connally, N.J. et al. The missing link between genetic association and regulatory function. Elife 11(2022).

76. Boye, C., Nirmalan, S., Ranjbaran, A. & Luca, F. Genotype□×□environment interactions in gene regulation and complex traits. Nature Genetics 56, 1057–1068 (2024).

77. Avsec, Ž., et al. AlphaGenome: advancing regulatory variant effect prediction with a unified DNA sequence model. bioRxiv, 2025.06.25.661532 (2025).

78. Chen, S., Zhou, Y., Chen, Y. & Gu, J. fastp: an ultra-fast all-in-one FASTQ preprocessor. Bioinformatics 34, i884–i890 (2018).

79. Dobin, A. et al. STAR: ultrafast universal RNA-seq aligner. Bioinformatics 29, 15–21 (2013).

80. Liao, Y., Smyth, G.K. & Shi, W. featureCounts: an efficient general purpose program for assigning sequence reads to genomic features. Bioinformatics 30, 923–30 (2014).

81. Pertea, M., Kim, D., Pertea, G.M., Leek, J.T. & Salzberg, S.L. Transcript-level expression analysis of RNA-seq experiments with HISAT, StringTie and Ballgown. Nat Protoc 11, 1650–67 (2016).

82. R Core Team, R. R: A language and environment for statistical computing. (Citeseer, 2020).

83. van der Maaten, L. & Hinton, G. Visualizing high-dimensional data using t-sne. 2008. Journal of Machine Learning Research 9, 2579.

84. Zhou, H.J., Li, L., Li, Y., Li, W. & Li, J.J. PCA outperforms popular hidden variable inference methods for molecular QTL mapping. Genome Biology 23, 210 (2022).

85. Ritchie, M.E. et al. limma powers differential expression analyses for RNA-sequencing and microarray studies. Nucleic Acids Research 43, e47–e47 (2015).

86. Yu, G. Using ggtree to Visualize Data on Tree-Like Structures. Current Protocols in Bioinformatics 69, e96 (2020).

87. Xu, S. et al. ggtreeExtra: Compact Visualization of Richly Annotated Phylogenetic Data. Molecular Biology and Evolution 38, 4039–4042 (2021).

88. Bolger, A.M., Lohse, M. & Usadel, B. Trimmomatic: a flexible trimmer for Illumina sequence data. Bioinformatics 30, 2114–20 (2014).

89. Li, H. Aligning sequence reads, clone sequences and assembly contigs with BWA-MEM. arXiv (2013).

90. Danecek, P. et al. Twelve years of SAMtools and BCFtools. GigaScience 10(2021).

91. McKenna, A. et al. The Genome Analysis Toolkit: a MapReduce framework for analyzing next-generation DNA sequencing data. Genome Research 20, 1297–303 (2010).

92. Browning, B.L., Tian, X., Zhou, Y. & Browning, S.R. Fast two-stage phasing of large-scale sequence data. Am J Hum Genet 108, 1880–1890 (2021).

93. Browning, B.L., Zhou, Y. & Browning, S.R. A One-Penny Imputed Genome from Next-Generation Reference Panels. The American Journal of Human Genetics 103, 338–348 (2018).

94. Rubinacci, S., Hofmeister, R.J., Sousa da Mota, B. & Delaneau, O. Imputation of low-coverage sequencing data from 150,119 UK Biobank genomes. Nature Genetics 55, 1088–1090 (2023).

95. Asiimwe, R. & Dobin, A. STAR+WASP reduces reference bias in the allele-specific mapping of RNA-seq reads. 2024.01.21.576391 (2024).

96. Patro, R., Duggal, G., Love, M.I., Irizarry, R.A. & Kingsford, C. Salmon provides fast and bias-aware quantification of transcript expression. Nature Methods 14, 417–419 (2017).

97. Li, Y.I. et al. Annotation-free quantification of RNA splicing using LeafCutter. Nature Genetics 50, 151–158 (2018).

98. Munro, D. et al. Multimodal analysis of RNA sequencing data powers discovery of complex trait genetics. Nature Communications 15, 10387 (2024).

99. Zou, X. et al. Using population-scale transcriptomic and genomic data to map 3′ UTR alternative polyadenylation quantitative trait loci. STAR Protocols 3, 101566 (2022).

100. Feng, X., Li, L., Wagner, E.J. & Li, W. TC3A: The Cancer 3′ UTR Atlas. Nucleic Acids Research 46, D1027–D1030 (2017).

101. Quinlan, A.R. & Hall, I.M. BEDTools: a flexible suite of utilities for comparing genomic features. Bioinformatics 26, 841–2 (2010).

102. Purcell, S. et al. PLINK: a tool set for whole-genome association and population-based linkage analyses. American Journal of Human Genetics 81, 559–75 (2007).

103. Robinson, M.D. & Oshlack, A. A scaling normalization method for differential expression analysis of RNA-seq data. Genome Biol 11, R25 (2010).

104. Battle, A., Brown, C.D., Engelhardt, B.E. & Montgomery, S.B. Genetic effects on gene expression across human tissues. Nature 550, 204–213 (2017).

105. Zou, Y., Carbonetto, P., Wang, G. & Stephens, M. Fine-mapping from summary data with the “Sum of Single Effects” model. PLOS Genetics 18, e1010299 (2022).

106. Wang, G., Sarkar, A., Carbonetto, P. & Stephens, M. A simple new approach to variable selection in regression, with application to genetic fine mapping. J R Stat Soc Series B Stat Methodol 82, 1273–1300 (2020).

107. Giambartolomei, C. et al. Bayesian test for colocalisation between pairs of genetic association studies using summary statistics. PLoS Genet 10, e1004383 (2014).

108. Wu, Y. et al. Joint analysis of GWAS and multi-omics QTL summary statistics reveals a large fraction of GWAS signals shared with molecular phenotypes. Cell Genomics 3, 100344 (2023).

109. Urbut, S.M., Wang, G., Carbonetto, P. & Stephens, M. Flexible statistical methods for estimating and testing effects in genomic studies with multiple conditions. Nature Genetics 51, 187–195 (2019).

110. Vavrek, M.J. Fossil: palaeoecological and palaeogeographical analysis tools. Palaeontologia electronica 14, 16 (2011).

111. Alexander, D.H., Novembre, J. & Lange, K. Fast model-based estimation of ancestry in unrelated individuals. Genome Res 19, 1655–64 (2009).

112. Gao, B. & Zhou, X. MESuSiE enables scalable and powerful multi-ancestry fine-mapping of causal variants in genome-wide association studies. Nature Genetics (2024).

113. Willer, C.J., Li, Y. & Abecasis, G.R. METAL: fast and efficient meta-analysis of genomewide association scans. Bioinformatics 26, 2190–1 (2010).

114. Bouwman, A.C. et al. Meta-analysis of genome-wide association studies for cattle stature identifies common genes that regulate body size in mammals. Nature Genetics 50, 362–367 (2018).

115. Speed, D., Holmes, J. & Balding, D.J. Evaluating and improving heritability models using summary statistics. Nat Genet 52, 458–462 (2020).

116. Weir, B.S. & Cockerham, C.C. Estimating f-statistics for the analysis of population structure. Evolution 38, 1358–1370 (1984).

117. Danecek, P. et al. The variant call format and VCFtools. Bioinformatics 27, 2156–2158 (2011).

118. Wang, K., Li, M. & Hakonarson, H. ANNOVAR: functional annotation of genetic variants from high-throughput sequencing data. Nucleic Acids Research 38, e164–e164 (2010).

119. Mostafavi, H., Spence, J.P., Naqvi, S. & Pritchard, J.K. Systematic differences in discovery of genetic effects on gene expression and complex traits. Nature Genetics 55, 1866–1875 (2023).

120. Yang, J., Lee, S.H., Goddard, M.E. & Visscher, P.M. GCTA: a tool for genome-wide complex trait analysis. Am J Hum Genet 88, 76–82 (2011).

121. Martin, M. Cutadapt removes adapter sequences from high-throughput sequencing reads. EMBnet. journal 17, 10–12 (2011).

122. Schubert, M., Lindgreen, S. & Orlando, L. AdapterRemoval v2: rapid adapter trimming, identification, and read merging. BMC Res Notes 9, 88 (2016).

123. Li, H. & Durbin, R. Fast and accurate short read alignment with Burrows-Wheeler transform. Bioinformatics 25, 1754–60 (2009).

124. Jonsson, H., Ginolhac, A., Schubert, M., Johnson, P.L. & Orlando, L. mapDamage2.0: fast approximate Bayesian estimates of ancient DNA damage parameters. Bioinformatics 29, 1682–4 (2013).

125. Kumar, L. & M, E.F. Mfuzz: a software package for soft clustering of microarray data. Bioinformation 2, 5–7 (2007).

