## Supplementary_note for "A multi-tissue atlas of genetic regulatory effects in sheep"

The supplementary information contains:

**Supplementary Note**

**Supplementary Figures 1–33**

**Supplementary References**

**Supplementary Tables 1–26** (in a separate Excel file)

**Supplementary Note**

**Tissue sample collection**

In this study, 10 juvenile and six adult males, along with 11 juvenile and 44 adult females of Small-tailed Han sheep from Yuncheng County, Shandong Province (China), were selected. In addition, 30 adult male Sunite sheep from the Sonid Grassland, Inner Mongolia Autonomous Region (China), and one adult male and 13 adult female Tibetan sheep from Damxung County, Tibet Autonomous Region (China) were included. All animals were maintained with balanced nutrition, adequate drinking water, and routine health management. After a 12-hour fasting period, the animals were humanely euthanized, and 1–42 tissue samples per individual were rapidly dissected and collected. Tissues included: heart; abomasum; large intestine (cecum, colon, rectum); liver; omasum; reticulum; rumen; small intestine (duodenum, ileum, jejunum); adipose depots (intermuscular, perirenal, subcutaneous, tail); thyroid; mammary gland; ovary; oviduct; uterus (cervix); bone marrow; lymph node; spleen; thymus; soft horn; skin; epididymis; testis; longissimus muscle; brain regions (brainstem, cerebellum, cerebral cortex, hippocampus, hypothalamus, medulla oblongata, optic chiasm, pineal, pituitary); lung; trachea; periosteum; and kidney. At the end of the experiment, all samples were properly disposed of, and the environment was thoroughly cleaned and disinfected.

**RNA extraction and sequencing**

Total RNA was extracted using TRIzol reagent (Invitrogen, CA, USA) following the manufacturer’s protocol. RNA purity and concentration were assessed with a NanoDrop 2000 spectrophotometer (Thermo Fisher Scientific, USA), and RNA integrity was evaluated using an Agilent 2100 Bioanalyzer (Agilent Technologies, Santa Clara, CA, USA). Strand-specific RNA-seq libraries were prepared using the VAHTS Universal V10 RNA-seq Library Prep Kit (Premixed Version), according to the manufacturer’s instructions. Library construction and transcriptome sequencing were performed by OE Biotech Co., Ltd. (Shanghai, China). Paired-end 150-bp reads were generated on the DNBSEQ-T7 platform. In total, we generated 1,900 RNA-seq datasets from sheep across 47 tissues types (**Supplementary Table 1**), including 1,777 samples (1–42 tissues per individual) from 71Small-tailed Han sheep, 61 samples (1–4 tissues per individual) from 30 Sunite sheep, and 62 samples (2–6 tissues per individual) from 14 Tibetan sheep. Raw sequence data are available for download at the Sequence Read Archive under accessions PRJNA1198671 and PRJNA1304012.

**DNA extraction and sequencing**

Of the 1,900 newly generated RNA-seq samples, 1,300 had matched whole-genome sequencing (WGS) data from 35 Small-tailed Han sheep (**Supplementary Table 2**). Genomic DNA was extracted from whole blood using the CWE9600 Magbead Blood DNA Kit following a magnetic bead-based protocol. DNA quality and integrity were assessed using standard quality control procedures provided by BGI. For DNB-seq library preparation, ≥ 1 μg of genomic DNA was randomly fragmented to ~300–350 bp using Covaris^TM^ ultrasonication, followed by end repair, A-tailing, and adapter ligation. Size selection was performed using NadPrep® SP Beads, and libraries were amplified by PCR and purified. Final libraries were quantified using Qubit 2.0 and evaluated for insert size using the Agilent Bioanalyzer. Qualified libraries were pooled and sequenced on the DNBSEQ-T7 platform (MGI) using paired-end 150 bp reads (PE150). Sequencing was based on DNA nanoball (DNB) technology, and base calling was performed to generate raw reads for downstream analyses. Raw sequence data are available at the Sequence Read Archive under accession PRJNA1403714.

**Decomposition of gene expression variance**

To identify key contributors to gene expression variability, we used the variancePartition package (v1.30.2)^1^ to quantify the proportion of expression variance attributable to tissue, individual, sex, developmental stage, genetic background, and batch effect (BioProject). We first computed the coefficient of variation (CV) of transcripts per million (TPM) values for each gene, and stratified genes based on log-transformed CV into the top 2,000 most and least variable genes, respectively. Gene Ontology (GO) enrichment analysis, performed with the *enrichGO* function from clusterProfiler (v4.8.3)^2^ using the org.Hs.eg.db (v3.16.0) database, revealed that highly variable genes were enriched for tissue-specific functions, while genes with low variability were associated with fundamental cellular processes (**Supplementary Fig. 3a; Supplementary Table 3**). We subsequently applied variancePartition using a linear mixed model to quantify the contribution of each variable to gene expression variance. The input included log-transformed gene expression values [log(TPM + 0.25)] and metadata such as individual ID, tissue, breed, sex, developmental stage, and BioProject accession (retrieved from NCBI). Batch effects were estimated as expression principal components (PCs) within tissue using PCAForQTL package^3^ and removed with the *removeBatchEffect* function from the limma package (v3.56.2)^4^. As expected, tissue type accounted for the largest proportion of expression variance across genes. Notably, the contribution of BioProject (a proxy for batch effects) was substantially reduced following batch correction (**Supplementary Fig. 3b**).

**Tissue-specificity and co-expression analysis of gene expression**

We assessed tissue-specific gene expressions across 51 tissues, each containing at least 40 RNA-seq samples. Tissue specificity was quantified using the TAU value, implemented in the tspex package (v0.6.3)^5^. Differential expression analysis was performed by comparing each tissue to all others using the Wilcoxon rank-sum test^6^, based on trimmed mean of M value (TMM)-normalized expression values. Highly expressed genes were classified as tissue-specific if they met the following criteria: FDR corrected *P*-value < 0.05 and Log_2_FoldChange > 2 relative to all other tissues. Functional enrichment analysis of tissue-specific genes was conducted within tissue for Gene Ontology (GO) and Biological Process (BP) terms using clusterProfiler (v4.8.3)^2^. We also examined the expression patterns of tissue-specific genes previously associated with Mendelian traits, curated from the Online Mendelian Inheritance in Animals (OMIA) database^7^.

Gene co-expression networks were constructed within each tissue using weighted gene co-expression network analysis (WGCNA, v1.73)^8^. Before network construction, gene expression levels were adjusted for hidden confounders by regressing out expression PCs using PCAForQTL package. Annotation of genes to GO terms was obtained via the biomaRt R package (v2.60.1)^9^. GO-based functional enrichment was again performed using clusterProfiler, and co-expression networks were visualized with Gephi (v0.10.1)^10^.

**Variant calling and genotyping from WGS data**

WGS data from 93 individuals with matched RNA-seq samples (**Supplementary Table 2**) was processed using a standardized pipeline based on BaseNumber (v1.0.3)^11^ for downstream analyses, including validation of genotype imputation from matched RNA-seq data and *cis*-eQTL mapping. Raw reads were quality-trimmed with fastp (v0.20.0)^12^ to remove adapter sequences and low-quality bases prior to alignment to the sheep reference genome (ARS-UI_Ramb_v2.0, GCA_016772045.1) using BWA-MEM (v0.7.17)^13^. Aligned reads were sorted with SAMtools (v1.9)^14^, and PCR duplicates were marked and removed using Picard (v2.20.1) (<https://broadinstitute.github.io/picard>). Variant calling was performed with GATK (v4.1.5)^15^. Per-sample gVCFs were generated using HaplotypeCaller, merged via CombineGVCFs, and jointly genotyped with GenotypeGVCFs, including non-variant sites (--include-non-variant-sites). SNP filtering was applied with VariantFiltration using thresholds: QD < 2.0, FS > 60.0, MQRankSum < −12.5, ReadPosRankSum < −8.0, SOR > 3.0, MQ < 40.0, and ExcessHet > 54.69. High-confidence variants were retained using BCFtools (v1.16)^14^ with criteria of (1) maximum alleles ≤ 2; (2) missing rate < 0.1; and (3) read depth between 10× and 60×.

**Evaluation of genotype imputation from RNA-seq data**

We performed genotype phasing and imputation for the filtered variants called from all 8,171 RNA-seq alignments using GLIMPSE2 (v2.0.0)^16^, leveraging a multi-breed sheep reference panel constructed from 3,125 WGS individuals (**Supplementary Table 5**). Variants with minor allele frequency (MAF) < 0.05 and IMPUTE info quality score (INFO) < 0.75 were excluded, resulting in 3,427,511 high-confidence SNPs used for subsequent molecular quantitative trait loci (molQTL) mapping. To evaluate the representativeness of the reference panel for RNA-seq-based imputation, we pruned imputed SNPs in high linkage disequilibrium (LD) using PLINK (v1.90b7)^17^, specifying the parameters --indep-pairwise 50 5 0.2, yielding 156,560 LD-independent SNPs. Principal component analysis (PCA) of 3,125 WGS reference individuals and the 8,171 RNA-seq samples revealed highly similar population structures, confirming the appropriateness of the reference panel for RNA-seq-based imputation (**Extended Data Fig. 2a**).

To assess best practices for RNA-seq genotype imputation, we benchmarked imputation accuracy on chromosome 24 using 1,300 RNA-seq samples with matched WGS data from 35 Small-tailed Han sheep, which were not included in the reference panel (**Supplementary Table 2**). Imputation accuracy was evaluated using genotype concordance rate (CR) and genotype correlation (*ρ*^2^) between imputed RNA-seq and known WGS genotypes. We compared performance across a range of effective population sizes (*N*_e_) and INFO score thresholds. Imputation accuracy was largely consistent across *N*_e_ values for variants with high INFO scores (≥ 0.75) (**Extended Data Fig. 2b**). Based on this observation, we set *N*_e_ = 1,000 for all subsequent imputation analyses, in accordance with the PigGTEx pipeline^18^.

As SNP calling with GATK followed by phasing and imputation using BEAGLE has been widely adopted in previous FarmGTEx studies^18-20^, we next benchmarked three imputation strategies on chromosome 24 using the 1,300 RNA-seq data with matched WGS described above: (1) BEAGLE_ind, in which SNPs were individually called per sample using GATK (v4.3.0.0)^15^, and phased and imputed using BEAGLE (v5.4)^21,22^; (2) BEAGLE_pop, in which SNPs were jointly called across individuals using GATK and then phased and imputed with BEAGLE; and (3) GLIMPSE2, using population-based imputation directly from BAM files. For GATK-based pipelines, low-quality variants were filtered using the criteria FS > 30.0, QD < 2.0 or DP < 5.0. BEAGLE-imputed variants were further filtered by model-based imputation accuracy (DR^2^), while GLIMPSE2-imputed variants were filtered by INFO score. Both BEAGLE_pop and GLIMPSE2 achieved satisfactory performance (CR ≥ 0.95 and *ρ*^2^ ≥ 0.85) when applying thresholds of INFO ≥ 0.75 for GLIMPSE2 and DR^2^ ≥ 0.85 for BEAGLE-based methods (**Extended Data Fig. 2c**). However, the number of retained SNPs was substantially higher for GLIMPSE2 (*n* = 81,016) compared to BEAGLE_ind (*n* = 44,019) and BEAGLE_pop (*n* = 54,092) (**Extended Data Fig. 2d**). Notably, GLIMPSE2 recovered > 98.8% of the SNPs identified by the other two strategies and uniquely identified 29.7% (*n* = 29,774) additional SNPs (**Extended Data Fig. 2e**). Based on these results, we adopted GLIMPSE2 for genome-wide imputation in this study and applied final filters of MAF ≥ 0.05 and INFO ≥ 0.75, yielding 3,427,511 high-confidence SNPs for downstream analyses, with a median CR of 96.0% and *ρ*^2^ of 85.1% (**Extended Data Fig. 2f**). Additional evaluations of imputation accuracy stratified by MAF and restricted to SNPs within *cis*-windows (±1 Mb from the transcription start site (TSS), the interval used for *cis*-molQTL mapping) further supported the robustness of the approach (**Extended Data Fig. 2g,h**).

To further assess RNA-seq-based genotype imputation accuracy, we analyzed 2,350 samples from three breeds independent of the reference panel (including the Small-tailed Han population described above) within the SheepGTEx discovery cohort (**Supplementary Table 2**), together with 378 validation samples from six additional populations/breeds outside the SheepGTEx resource. These included 245 (East Friesian × Hu) × Hu crossbreeds, 112 Hu sheep^23^, four Charollais, nine Suffolk, one Texel and seven Australian white sheep. All samples had matched WGS data independent of the reference panel. Using GLIMPSE2 and the multi-breed reference panel, we imputed the 3,427,511 high-confidence SNPs from RNA-seq to sequence level. Population structure analyses confirmed that these validation samples align closely with the broader SheepGTEx cohort, and that their matched WGS data are well-represented within our global reference panel (**Extended Data Fig. 2i**), supporting the ancestry representativeness of the dataset. In addition, the distributions of sex and developmental stages in the validation dataset are broad and balanced across the included breeds (**Extended Data Fig. 2j**), reflecting the diversity of the full atlas. We evaluated imputation performance across all nine populations by comparing RNA-seq-imputed genotypes with WGS-called “gold-standard” genotypes. Across 3,427,511 high-quality SNPs, we observed a median concordance rate of 94.7% and a genotype correlation of 90.2% (**Extended Data Fig. 2k**). Similar accuracy was observed across genomic features (**Extended Data Fig. 2l**).

Collectively, these results provide evidence that our genotype imputation accuracy is sufficient and generalizes effectively across the diverse ancestry groups, sexes, and developmental stages represented in the SheepGTEx resource.

**Sex, developmental stage and breed prediction**

We inferred sex labels for SheepGTEx samples using both genome coverage- and gene expression-based approaches (**Supplementary Fig. 6a**). For the coverage-based method, we adapted scripts from ancient goat genome analyses^24^, calculating the mean SNP depth on autosomes, the X chromosome, and single-copy regions of the Y chromosome^25^. Relative X- and Y-coverage (X-rate and Y-rate) were computed, and a threshold of log_10_(Y-rate/X-rate) = 0.05, determined from 2,440 WGS-validated samples, was used to distinguish males from females (**Supplementary Fig. 6b,c**). For the expression-based method, we trained a multi-layer perceptron classifier on the TPM matrix using the same WGS-validated set, selecting the top 20 ANOVA-ranked features in scikit-learn, following the FarmGTEx metadata prediction pipeline (<https://github.com/FarmGTEx/metadata-prediction-v1>). Cross-validation (1-50 folds) achieved mean accuracy > 99.7% (**Supplementary Fig. 6d**), and the two methods showed 97.0% concordance (**Supplementary Fig. 6e**). Discordant samples were manually curated using sex-specific tissue information (e.g., testis or mammary gland) when available; otherwise, the coverage-based result was retained.

We adapted a previously established framework for *Arabidopsis thaliana* RNA-seq data^26^, applying a Random Forest-based approach to predict the missing developmental stage labels of bulk RNA-seq samples in sheep (**Supplementary Fig. 7**). Briefly, the TPM-normalized expression matrix was reformatted to a sample-by-gene layout, and available age annotations were integrated, with missing values labeled as NA. A Random Forest classifier (1,500 trees, min_samples_leaf = 3, max_features = ‘sqrt’, max_depth = 48, bootstrap = False) was trained on 70% of age-annotated samples, with performance assessed on the remaining 30% using overall accuracy, per-class precision, recall, and F1-scores, and the confusion matrix implemented in sklearn.metrics package. The trained model was then applied to unannotated samples to generate developmental stage predictions, which were merged with sample metadata to produce the final output. Notably, this framework implicitly incorporates Median Ratio Normalization through TPM input and uses parameter settings previously optimized for transcriptome-based trait prediction.

We subsequently inferred missing breed labels from RNA-seq-imputed genotypes following established approaches^27,28^ (**Supplementary Fig. 8**). The dataset comprised a total of 8,171 samples with 943 unknown labels. After quality control, including outlier removal via principal component analysis (PCA) and exclusion of breeds with ≤ 5 samples, 6,284 samples remained. To minimize redundancy and improve prediction efficiency, breed-informative SNPs were selected based on fixation index (*F*_ST_) and average Euclidean distance (AED). We evaluated the impact of SNP set size on classification accuracy and compared four machine-learning algorithms (K-nearest neighbors, random forest, support vector machine and XGBoost) using cross-validation. The optimal performance was achieved with 50K SNPs selected by *F*_ST_ and classified using SVM, which was subsequently applied for final breed assignment.

**Detection of duplicate RNA-seq samples within each tissue**

To remove duplicate samples originating from the same individuals within each tissue, we calculated identity-by-state (IBS) distances using PLINK (v1.90b7)^17^ and genetic relationship matrices (GRM) using GCTA (v1.94.1)^29^ for all 8,171 RNA-seq samples based on 3,427,511 high-confidence SNPs. IBS and GRM values from imputed genotypes showed > 99% concordance with those derived from matched WGS data across six data resources described in **Evaluation of genotype imputation from RNA-seq data** (**Supplementary Fig. 9a,c**). To verify detection accuracy, we compared pairwise IBS and GRM values for 2,350 RNA-seq samples from 93 known individuals (1,300 from 35 Small-tailed Han sheep, 868 from 48 Hu sheep, and 182 from 10 Tibetan sheep), revealing distinct distributions between duplicate and unrelated samples, with optimal separation at IBS = 0.9 and GRM = 0.5 (**Supplementary Fig. 9b,d**). Clustering analysis further confirmed that duplicates grouped together under these cutoffs (**Supplementary Fig. 9e,f**). Accordingly, among the remaining 5,821 samples, any pair with IBS ≥ 0.9 and GRM ≥ 0.5 was flagged as a duplicate, and the replicate with the highest sequencing depth was retained. After filtering, 51 tissues with sufficient sample sizes (n ≥ 40) were retained for molQTL mapping, ranging from 40 samples (rectum, oviduct) to 713 (muscle) (**Supplementary Figs. 1o and 9g; Supplementary Table 6**).

**Covariate analysis**

To account for hidden technical and biological confounders, we computed phenotype principal components (PCs) for each tissue using the PCAForQTL package, with the number of PCs determined automatically by the *runElbow* function (**Supplementary Fig. 11a**). Given that previous GTEx studies inferred hidden variables^18-20,30^ using the Probabilistic Estimation of Expression Residuals (PEER) method^31,32^, we compared the gene expression variance explained by PEER factors with that explained by phenotype PCs. The number of PEER factors per tissue was chosen based on the Human GTEx study design^30^. Across tissues with large sample sizes (n > 70), the variance explained by PEER factors closely matched that explained by expression PCs (**Supplementary Fig. 11b–d**). In contrast, tissues with smaller sample sizes (n < 70) showed weaker correlations, likely reflecting the high collinearity among PEER factors reported previously^3^ (**Supplementary Fig. 11e–g**). Given the comparable or superior performance of PCA, we adopted phenotype PCs, rather than PEER factors, as covariates in subsequent analyses. To account for population structure, we computed genotype PCs for each tissue using PLINK (v1.90b7) on imputed genotypes. The top 10 genotype PCs captured most of the variance in population structure (**Supplementary Fig. 11h–j**). We included the inferred phenotype PCs, the top five genotype PCs for tissues with sample sizes < 200 (or the top ten for ≥ 200), and sex and age as covariates for heritability estimation and molQTL mapping.

**Heritability estimation of seven molecular phenotypes**

To quantify the genetic contribution to variation in each molecular phenotype, we estimated *cis*- and *trans*-heritability (*h*^2^) per tissue using OmiGA (v1.0.1)^33^ (**Supplementary Fig. 12a,b**), incorporating all covariates described above. Additive GRM was computed from *cis*- or *trans*-variants, and heritability was estimated using a linear mixed model with Average Information Restricted Maximum Likelihood (AI-REML) method^34^. For gene expression, *cis*-*h*^2^ was positively correlated with expression level but negatively correlated with gene network connectivity, measured by the absolute module eigengene-based connectivity (|kME|) (**Supplementary Fig. 12c**). Across the seven types of molecular phenotypes, an average of 11.5% and 20.4% of molecular phenotypes exhibited significant *cis*- and *trans*-*h*^2^ (one-sided Wald-test *P* < 0.05), respectively (**Supplementary Fig. 13a**). The proportion of significant heritability estimates was strongly influenced by tissue sample size (**Supplementary Fig. 13b**). Molecular phenotypes with significant heritability showed broadly similar distribution patterns, but with generally higher *h*^2^ estimates (**Supplementary Fig. 13c**), compared to analyses based on all *h*^2^ estimates (**Supplementary Fig. 12a**).

As expected, molecular phenotypes with significant *cis*-molQTL (i.e., molGenes) showed higher *cis*-*h*^2^ than those without significant *cis*-molQTL (non-molGenes; **Supplementary Fig. 14a**). Taking gene expression as an example, 76.1% of eGenes displayed significant *cis*-*h*^2^, compared with only 9.8% of non-eGenes (**Supplementary Fig. 14b**). To further evaluate whether filtering genes based on significant *cis*-heritability (*cis*-*h*^2^) is appropriate for molQTL mapping, we conducted a simulation study using a dataset of 300 individuals from our previous study^33^, which includes 3,556 known eGenes and 1,879 non-eGenes. We first estimated *cis*-*h*^2^ for all simulated genes using the Average Information Restricted Maximum Likelihood (AI-REML) method implemented in OmiGA^33^. Across 50 simulation replicates, 42.71% and 0.29% of known eGenes and non-eGenes showed significant *cis*-*h^2^* estimates, respectively. In addition, 99.6% of genes with significant *cis*-*h*^2^ estimates were indeed true eGenes, while 51.3% of genes with non-significant *cis*-*h*^2^ estimates were also detected as true eGenes (**Supplementary Fig. 14c**). Therefore, we did not filter molecular phenotypes based on their *cis*-*h*^2^ significance for downstream molQTL mapping.

***cis*-eQTL evaluation and validation**

We characterized *cis*-eQTL across 51 sheep tissues. As expected, genes without detectable *cis*-eQTL (non-eGenes) were more tissue-specific, harbored more rare variants, showed lower expression levels, were located further from the TSS, and exhibited reduced *cis*-*h*^2^ (**Supplementary Fig. 16a–f**). Highly expressed non-eGenes were typically enriched for tissue-specific functional pathways, whereas lowly expressed non-eGenes were enriched for fundamental GO categories (**Supplementary Fig. 16g**). Consistent with previous findings^19^, the power to detect molQTL (particularly those with small effect sizes) increased with tissue sample size (**Fig. 2a; Supplementary Fig. 16h**). Notably, molQTL with smaller effect sizes tended to map closer to the TSS, yet their target genes were less central in co-expression networks (**Supplementary Fig. 16i,j**), suggesting more context-dependent regulatory functions.

To assess the impact of genetic relatedness on *cis*-molQTL mapping, we compared results from TensorQTL (v1.0.9)^35^, which uses a linear regression model (LM), and OmiGA (v1.0.1)^33^, which implements a linear mixed model (LMM). The sets of eGenes identified by the two approaches showed substantial overlap (**Supplementary Fig. 17a–c**). Furthermore, key summary statistics, including Z-scores (slope/slope_se), effect sizes, and significance levels (−log_10_*P*), were highly correlated between LM and LMM (Pearson’s *r* = 0.994, 0.993 and 0.985 for Z-score, effect size and significance level, respectively; **Supplementary Fig. 17d,e**), indicating that both approaches effectively captured and accounted for spurious population genetic effects.

We performed internal validation of *cis*-eQTL in 33 tissues with sample sizes > 100. For each tissue, samples were randomly split into two equal groups, and *cis*-eQTL mapping was conducted separately in each group using TensorQTL (**Supplementary Fig. 18a**). Validation rates were quantified using Storey’s π_1_ statistic implemented in the qvalue package (v2.32.0)^36^. The π_1_ statistic was defined as the proportion of *cis*-eQTL in one group that were also significant in the other^37^. In addition, we calculated Pearson’s correlations of normalized effect sizes (Z-scores) between groups.

Allele-specific expression (ASE) analysis was further performed to validate *cis*-eQTL by assessing the concordance of allelic fold change (aFC). *cis*-eQTL aFC was estimated using aFC.py (v0.3)^38^ based on genotypes, normalized molecular phenotypes, and the same covariates used in *cis*-eQTL mapping, with 95% confidence intervals derived from 100 bootstraps. For ASE analysis, haplotypes were phased using phASER (v1.1.1)^39^ with options: --mapq 255 --baseq 10 --gw_phase_vcf 1 --pass_only 0, applied to WASP-filtered alignments to remove reads with allelic mapping bias (see **Methods**). Genomic regions with high mapping error rates (mappability < 0.5) in the sheep reference genome were masked, as estimated by GenMap (v1.3.0)^40^ with parameters: -K 75 -E 2. Gene-level haplotype counts were generated with phaser_gene_ae.py (v1.2.0) and aggregated across samples using phaser_expr_matrix.py (v0.1.0). ASE-based aFC was then calculated per tissue using phaser_cis_var.py (v0.1.0) for variants present in ≥ 10 individuals with ASE data and ≥ 8 reads per individual. Spearman’s correlations revealed strong concordance between ASE-derived and eQTL-derived aFC (**Supplementary Fig. 18b–d**).

To compare the impact of WGS- and RNA-based genotypes on molQTL detection, we used 2,350 RNA-seq samples in 21 tissues with 93 matched WGS, as described above. WGS genotypes were directly called and RNA-seq genotypes were imputed as described above. For each dataset, *cis*-eQTL mapping was performed using TensorQTL (v1.0.9), and results were compared between imputed RNA-seq genotypes and WGS-called genotypes (**Extended Data Fig. 4**).

For external/independent validation, we used muscle RNA-seq data from 112 Hu sheep with matched WGS data. RNA-seq genotypes were imputed using GLIMPSE2 (v2.0.0)^16^ based on the sheep reference panel, whereas WGS genotypes were directly called from the alignments. *cis*-eQTL mapping was performed with TensorQTL using either imputed RNA-seq genotypes or WGS-called genotypes. Validation rates (quantified using the π_1_ statistic) and Spearman’s correlation were computed by comparing *cis*-eQTL discovered in SheepGTEx with those in the two validation datasets (**Supplementary Fig. 18e–g**).

**Sequence conservation of molQTL**

To assess the evolutionary conservation of molQTL, we downloaded PhastCons scores for 100 vertebrate species from UCSC (<https://hgdownload.cse.ucsc.edu/goldenpath/hg38/phastCons100way/hg38.100way.phastCons>). Wiggle files were converted to BED format using convert2bed (v1.6) from BEDOPS^41^, and coordinates were lifted over from the human genome (GRCh38) to the sheep genome (ARS-UI_Ramb_v2.0) using LiftOver^42^ with parameters “-minMatch=0.2 -minBlocks=0.5”. For each gene, the mean PhastCons score across its sequence was used as the conservation metric. Gene-level conservation was quantified as the mean PhastCons score across the aligned sequence, retaining only genes with ≥ 50% of their length successfully mapped in LiftOver for downstream analysis.

**Functional enrichment analyses of molQTL**

To explore the molecular mechanisms underlying the detected regulatory variants, we assessed their enrichment across multiple biological features, including 19 sequence ontology categories annotated by SnpEff (v5.2a)^43^ and 15 chromatin states inferred from 43 tissues. Enrichment was quantified using two metrics: odds ratio (OR) and fold enrichment (FE).

OR was calculated as:

$$\mathrm{OR}=\frac{a/b}{c/d}$$

where a is the number of molQTL in functional regions, b is the number of other SNPs in functional regions, c is the number of molQTL in non-functional regions, and d is the number of other SNPs in non-functional regions.

FE was calculated as:

$$\mathrm{FE}=\frac{C/A}{B/D}$$

where C is the number of molQTL in functional regions, A is the total number of molQTL, B is the number of SNPs in functional regions, and D is the total number of SNPs.

To ensure that the observed enrichment patterns reflected the underlying biology of regulatory variants rather than confounding by local MAF or LD structure, we repeated all molQTL enrichment analyses using control SNPs matched for both MAF and LD score, following established protocols^44^. Briefly, we used common SNPs (MAF > 0.05) from 2,816 unrelated sheep to calculate MAFs and LD scores for all variants using GCTA (v1.94.1)^29^ with default parameters. For each lead or fine-mapped variant (focal SNP), control SNPs were randomly sampled from a pool of 3,346,402 variants (excluding focal SNPs) under two matching criteria: (1) MAF within ±0.02 of the focal SNP; and (2) LD score within ±0.1 standard deviations of the focal SNP.

**Gene expression imputation and imputed *cis*-eQTL mapping**

We applied HYFA (hypergraph factorization)^45^, a parameter-efficient graph representation learning framework, for joint imputation of multi-tissue gene expression in sheep (**Supplementary Fig. 20a**). For each test individual, measured expression from source tissues was used to predict expression in the remaining target tissues, with performance evaluated by Pearson’s correlation against observed values. To determine the minimum training sample size, we used three accessible tissues (blood, skin, and subcutaneous adipose) as sources to impute expression in other tissues, based on TMM-normalized, inverse-normal–transformed data from 2,904 non-redundant individuals. Node features were initialized with demographic information (age and sex). Down-sampling experiments showed stable imputation performance when the number of training individuals exceeded 25 (**Supplementary Fig. 20b–d**). We therefore excluded tissues with < 25 individuals and individuals with expression data from only one tissue, yielding 1,614 individuals of 7,915 samples across 57 tissues for downstream analysis. Benchmarking six imputation approaches demonstrated that HYFA consistently outperformed alternatives, particularly when leveraging all available source tissues (**Supplementary Fig. 20e–h**). The final imputed gene expression matrix comprised 91,998 samples across 57 tissues.

Using the same pipeline as for observed data, we mapped *cis*-eQTL from the HYFA-imputed dataset and compared them with results from the original dataset to quantify the gain in regulatory variant discovery. Using imputed expression data (*n* = 1,614 across all tissues), we initially observed increased statistical power for eGene discovery compared to analyses based on raw expression data (**Extended Data Fig. 5a**). Consistently, in tissues with larger raw sample sizes, the number of shared eGenes increased, whereas the number of eGenes unique to imputed data decreased (**Extended Data Fig. 5b**). This pattern suggests that at least part of the additional eGenes identified from imputed data may reflect limited statistical power in the raw datasets for gene expression prediction, although false positives cannot be completely excluded. To further address this concern, we implemented a down-sampling strategy.

We utilized muscle tissue, which had the largest raw sample size (*n* = 713), as a “gold standard” truth set. Individuals were randomly down-sampled to training sets of varying sample sizes (*n* = 100, 200, 300, 400 and 500) to build expression prediction models using HYFA (hypergraph factorization)^45^. Gene expression was then imputed for the remaining individuals, followed by *cis*-eQTL mapping. We compared eGenes identified from the imputed datasets with those detected in the corresponding raw down-sampled datasets (**Extended Data Fig. 5c**). As training sample size increased, (1) the number of eGenes shared between raw and imputed datasets increased substantially, whereas (2) the number of eGenes uniquely identified in the imputed datasets progressively decreased.

To further evaluate the validity of these additional discoveries, we calculated the replication rate (Storey’s π_1_ statistic) of *cis*-eQTL uniquely identified in the imputed datasets relative to the corresponding raw down-sampled datasets, using the full muscle dataset (*n* = 713) as the gold-standard reference. We also assessed the Pearson’s correlation of normalized effect sizes between the unique *cis*-eQTL identified in the imputed datasets and those estimated from the gold-standard dataset. We observed high replication rates (π_1_ > 0.993) for the uniquely discovered imputed *cis*-eQTL, and their effect sizes were strongly correlated with those estimated in the gold-standard dataset at smaller training sample sizes (**Extended Data Fig. 5d–h**). Although the presence of a small number of false-positive associations, which decreased the effect-size correlation decreased as sample size became larger, the vast majority of additional *cis*-eQTL detected through imputation were supported by the gold-standard dataset. Together, these results demonstrate that expression imputation predominantly increases power rather than inflating false positives, with the greatest benefit observed in tissues with limited sample sizes.

Crucially, to address the biological validity of the “imputation-only” signals, we analyzed the functional enrichment of eQTL identified exclusively through imputed expression data. These “novel” signals exhibited significant enrichment near TSS, within active chromatin states (promoters and enhancers), and in conserved sequence features, mirroring the patterns observed for raw eQTL (**Extended Data Fig. 5l**). The fact that these novel signals reside in canonical regulatory regions argues against them being random false positives.

In summary, these analyses demonstrate that imputed expression data substantially enhanced discovery power while maintaining a high level of signal validity and biological interpretability.

***trans*-eQTL mapping**

We evaluated genomic control inflation factors (λ) comparing linear regression models implemented in TensorQTL (v1.0.9)^35^ and linear mixed models in OmiGA (v1.0.1)^33^ across 51 tissues, revealing minimal genomic inflation in association tests of *trans*-regions under the linear mixed model framework (**Supplementary Fig. 21a**). Subsequently, *trans*-eQTL mapping was performed in 15 tissues with >200 individuals. To reduce false positives, stringent filtering of variants and genes was applied following previous studies^18,30^. For variants, genome-wide SNP mappability was calculated using GenMap (v1.3.0)^40^, and SNPs with 75-mer mappability < 1 were removed. Variants located in repeat regions (UCSC RepeatMasker track: <https://hgdownload.soe.ucsc.edu/hubs/GCF/016/772/045/GCF_016772045.1/GCF_016772045.1.repeatMasker.out.gz>)^42^ were also excluded. Only SNPs with MAF > 0.05 were retained. For genes, exon mappability was computed using a 75-mer setting and 3’UTR mappability with a 36-mer setting. Cross-mappability between genes was then assessed using the crossmap (v1.2)^46^ pipeline, by aligning ambiguous k-mers from genes with mappability < 1 to other genes. Only genes with mappability ≥ 0.8 were included. *trans*-eQTL mapping was conducted using a linear mixed model implemented in OmiGA (v1.0.1)^33^, incorporating a genomic relationship matrix and the same covariates as in *cis*-eQTL mapping. For each gene, we excluded SNPs on the same chromosome or within ±1 Mb of any cross-mappable gene. Multiple testing correction followed the approach used in pig GTEx^18,30^. Specifically, the most extreme genome-wide *P*-value across all tested genes was multiplied by 10^6^, consistent with the estimated effective number of genome-wide tests. We then applied Benjamini-Hochberg correction at the gene level, defining *trans*-eGenes as those with FDR < 0.05. For each *trans*-eGene, variant-level *P*-values were further adjusted by Benjamini-Hochberg, and variants with FDR < 0.05 were considered significant *trans*-eQTL. GO enrichment analysis of the 135 *trans*-eGenes in 79 hotspots of lead *trans*-eQTL that affected multiple *trans*-eGenes was performed using *enrichGO* function from clusterProfiler package (v4.14.6)^2^ with the org.Hs.eg.db (v3.20.0) database.

***cis*-eQTL contribution to *trans*-eQTL**

To assess whether *trans* effects are influenced by *cis*-regulated genes, we first tested the enrichment of *trans*-eGenes with corresponding *cis*-eQTL by estimating the odds that their lead variants are also identified as *cis*-eQTL for any gene in the same tissue, following the approach described in the *Functional enrichment analyses of molQTL* section.

We further evaluated mediation for lead *trans*-eQTL that also exhibited a *cis*-eQTL in the matched tissue. Following a previous study^30^, we performed *trans*-eQTL mediation analysis using two-stage least squares (TSLS). Briefly, we first applied ridge regression (*α* = 100) to estimate the effect of genetic variation on the *cis*-eGene. We then used the resulting coefficients to predict *cis*-eGene expression for each individual and performed a second regression to estimate the causal effect of the predicted *cis*-eGene expression on the observed *trans*-eGene expression. To assess significance, we generated matched *β*_TSLS_ statistics using permuted *trans*-eGene expression levels (100 permutations per trio). False discovery rate (FDR < 0.05; Benjamini-Hochberg correction) was determined based on these empirical *P* values.

To further characterize downstream regulatory relationships, we identified putative *trans*-associations for *cis*-mediating genes using colocalization analysis with the coloc (v5.2.3) package^47^ (*coloc.abf* function). Using the *cis*-mediating eGenes identified above, we tested all inter-chromosomal *trans*-eGene pairs within each tissue, considering a ±1Mb window around the TSS of each gene. Candidate *trans*-associations were retained if the posterior probability of colocalization (PP_H4_) exceeded 0.8 and the corresponding *trans*-eGenes exhibited an average mappability > 0.8.

**Tissue-sharing patterns of *cis*-eQTL** **based on the down-sampled dataset**

To empirically evaluate the robustness of our conclusions, we performed a sensitivity analysis by down-sampling all tissues with large sample sizes (*n* > 100) to a fixed threshold of 100 individuals. We then re-ran the *cis*-eQTL mapping and tissue-sharing analysis using *mashr* (v0.2.79)^48^. The characteristic “U-shaped” distribution of tissue sharing (where most eQTL are either highly tissue-specific or broadly shared) remained highly consistent between the full and down-sampled datasets (**Supplementary Fig. 26a,b**). While we observed a modest increase in the proportion of tissue-specific eQTL in the down-sampled data (**Supplementary Fig. 26b**), this is the expected result of reduced statistical power, which tends to favor the detection of large-effect, tissue-specific signals over subtle, broadly shared effects. In addition, The cross-tissue correlation structure of effect sizes was well-preserved after down-sampling (Pearson’s *r* = 0.81; **Supplementary Fig. 26c**), indicating that the global regulatory architecture is not driven by sample-size outliers.

In summary, the high concordance between our full and down-sampled analyses demonstrated that our conclusions regarding tissue-sharing patterns of regulatory effects are robust to sample size variation.

**Breed-sharing patterns of *cis*-eQTL**

To investigate *cis*-eQTL sharing across populations, we analyzed 14 tissues with ≥ 40 samples in both European (EUR) and Central/East Asian (CEA) populations. Within each tissue, samples were stratified by ancestry and breed (*n* ≥ 40), and fine-mapped *cis*-eQTL analysis was performed separately for each group using TensorQTL (v1.0.9)^35^ and the *susie_rss* function^49^ from the susieR package (v0.12.35)^50^, following the procedures described in **Methods**. Replication of *cis*-eQTL between populations within a tissue was quantified using Storey’s π_1_ statistic and Pearson’s correlation of normalized eQTL effect sizes and compared to replication between tissues within a population. To further characterize sharing patterns across populations, we conducted a meta-analysis of *cis*-eQTL across tissues and population groups using *mashr* (v0.2.79)^48^, following the same pipeline used for tissue-sharing patterns of molQTL (**Methods**).

**Sex- and developmental stage-biased gene expression analysis**

Genes with low expression (counts ≥ 6 and TPM ≥ 0.1 in ≥ 80% of samples) were retained. Following strategies adopted in previous studies of sex-biased gene expression in humans^51^, batch effects were corrected for each tissue using SmartSVA (v0.1.3)^52^ to extract surrogate variables (SVs), which were included as covariates in the *removeBatchEffect* function of limma (v3.56.2)^4^. For sex-biased gene (sbGene) expression analysis, we retained tissues containing ≥ 10 samples from each sex. For developmental stage-biased gene (dbGene) expression analysis, stages with < 10 samples were excluded, and only tissues with ≥ 2 remaining stages were analyzed. Pairwise comparisons were conducted using the Wilcoxon rank-sum test^6^ on the corrected TMM-normalized expression values. Genes with Bonferroni-adjusted *P* < 0.05 were considered significantly biased, and those with |log_2_FoldChange| > 1 were classified as large-effect biased genes. For dbGene analysis, directionality was determined from the sign and magnitude of log_2_ fold-change, and only tissues with > 50 biased genes were retained for downstream analysis. Gene set enrichment analysis (GSEA) was performed using clusterProfiler (v4.8.3)^2^ within tissue restricted to genes identified as sex- or developmental stage-biased. Enrichment results across tissues were integrated via meta-analysis and visualized in Cytoscape (v3.10.2)^53^.

To evaluate the impact of different covariate adjustment strategies based on expression PCs versus SVs, we independently corrected expression matrices in each tissue using either SVs or expression PCs. The resulting corrected expression matrices were highly correlated between the two approaches (**Supplementary Fig. 30a,b**). Likewise, the numbers of identified sbGenes and dbGenes were also strongly correlated across tissues (**Supplementary Fig. 30c,d**). However, SV-adjusted analyses consistently identified a larger number of biased genes than PC-adjusted analyses, suggesting that PC-based correction imposes a stronger adjustment that may also partially remove biological variation correlated with sex or developmental stage.

**Evaluation of genotype imputation from ancient genome data**

To evaluate the performance of GLIMPSE2 (v2.0.0) in imputing ancient sheep genomes, we used two high-coverage samples previously published (ASTF002 and tps062; 10.89× and 5.87×, respectively)^54^ and systematically down-sampled their sequencing data to generate datasets across a range of coverage levels. The two high-coverage BAM files were aligned to the sheep reference genome (ARS-UI_Ramb_v2.0^55^, GCF_016772045.1). Variants for the two high-coverage samples were called using BCFtools (v1.17). Genotype likelihoods were first computed using BCFtools *mpileup* with the parameters: -q 30 -Q 30 -a “FORMAT/DP,FORMAT/AD,INFO/AD”^56^, followed by genotype calling using BCFtools *call*, restricted to the 60,840,500 high-quality SNPs in sheep genotype reference panel described above. Only biallelic SNPs were retained. Genotypes were further filtered by applying a genotype quality threshold (GQ > 30) and a depth filter requiring the observed depth to fall within one-third to three times the expected depth for each coverage level^57^. The resulting high-confidence variant set from the two high-coverage samples was treated as the “gold-standard” genotype sets for downstream accuracy assessment. We then down-sampled the two high-coverage BAM files independently to target depths of 0.1×, 0.2×, 0.3×, 0.4×, 0.5×, 0.75×, and 1× using SAMtools (v1.18) *view*. Prior to imputation of the down-sampled datasets, the reference genome was divided into 6-Mb chunks with a 1-Mb buffer using GLIMPSE2_chunk. Each down-sampled BAM file was then imputed independently using GLIMPSE2 based on the predefined chunk files. The resulting imputed BCF files were ligated using GLIMPSE2_ligate and converted to VCF format with BCFtools *convert*. Imputation accuracy, measured by concordance rates (CR) and Spearman’s correlation (*ρ*^2^), was assessed by comparison with the “gold-standard” set.

We first evaluated imputation accuracy separately for transitions and transversions, given their distinct error profiles in low-coverage and ancient DNA data. SNPs were classified as transitions (A↔G, C↔T) or transversions (all other substitutions), and imputation accuracy was calculated independently for each class. This analysis confirmed that transversions consistently exhibited higher imputation accuracy than transitions (**Supplementary Fig. 31b**), consistent with previous observations in ancient DNA imputation studies^56,57^. We further assessed site-level accuracy as a function of INFO score thresholds and imputation accuracy across all samples and observed high imputation accuracy (mean CR = 0.90 and mean *ρ*^2^ = 0.83) when applying an INFO threshold of ≥ 0.80, resulting in 27,866,951 SNPs with MAF > 0.01 retained for downstream analyses (**Supplementary Fig. 31c**). As expected, imputation performance showed a positive correlation with sequencing depth and MAF (**Supplementary Fig. 31d,e**).

**Comparative analysis between sheep and cattle, pig and human**

*Genome comparison and inference of ancestral alleles*

To reconstruct ancestral alleles among sheep, cattle, pig, and human, we generated pairwise whole-genome alignments using LAST (v1454)^58^, following the unified pipeline in Ruminant Genome Database^59^ ([http://animal.omics.pro/code/index.php/RGD/loadByGet?address[]=RGD/Documents/Pipeline.php](http://animal.omics.pro/code/index.php/RGD/loadByGet?address%5b%5d=RGD/Documents/Pipeline.php)) with minor modifications. The sheep reference genome (ARS-UI_Ramb_v2.0) was indexed with lastdb (-P0 -uNEAR -R01), and cross-species alignments were performed with last-train and lastal, followed by maf-swap, last-split and maf-sort to produce optimized one-to-one MAF files, which were further converted to PSL format with maf-convert. Target genomes were converted to 2bit format using faToTwoBit, and chain files were generated using axtChain (-linearGap=medium) from kentUtils (v302.1.0)^42^. Chromosome-level MAF files were prepared by renaming species identifiers, splitting alignments with mafSplit, and merging non-chromosomal scaffolds. Variants shared between the sheep reference panel and outgroups (cattle, pig, human) were extracted with cyvcf2^60^ in Python (v3.11.4), allele frequencies were calculated (get_ref_info; vectorized via NumPy^61^), and data were partitioned into 30 subsets for input into est-sfs (v2.04)^62^. Ancestral states were assigned based on posterior probability thresholds (p > 0.8: major allele ancestral; p < 0.2: minor allele ancestral; otherwise ambiguous), and results were merged across chromosomes for standardized reporting.

*eGene categorization by orthology*

To investigate the conservation of gene regulation, eGenes identified in each of the four species were partitioned into two distinct categories. Orthologous eGenes were defined as those with a one-to-one ortholog between sheep and the given species, based on Ensembl (v112) orthology annotations. Species-specific eGenes were defined as eGenes lacking a corresponding one-to-one ortholog in sheep.

*Cross-species comparison of eQTL effect size and heritability using ancestral alleles*

We established a high-confidence set of conserved regulatory loci by selecting variants that were significant, orthologous lead eQTL shared across human, cattle, and pig, and that carried the identical ancestral allele. For each locus, the corresponding SNP with the same ancestral allele was identified in sheep. Comparison of effect sizes (|log_2_(aFC)|) and heritability of these ancestral alleles across all four species facilitated the determination of the conservation of regulatory effects across a subset of mammalian species.

**Figures and Legends**

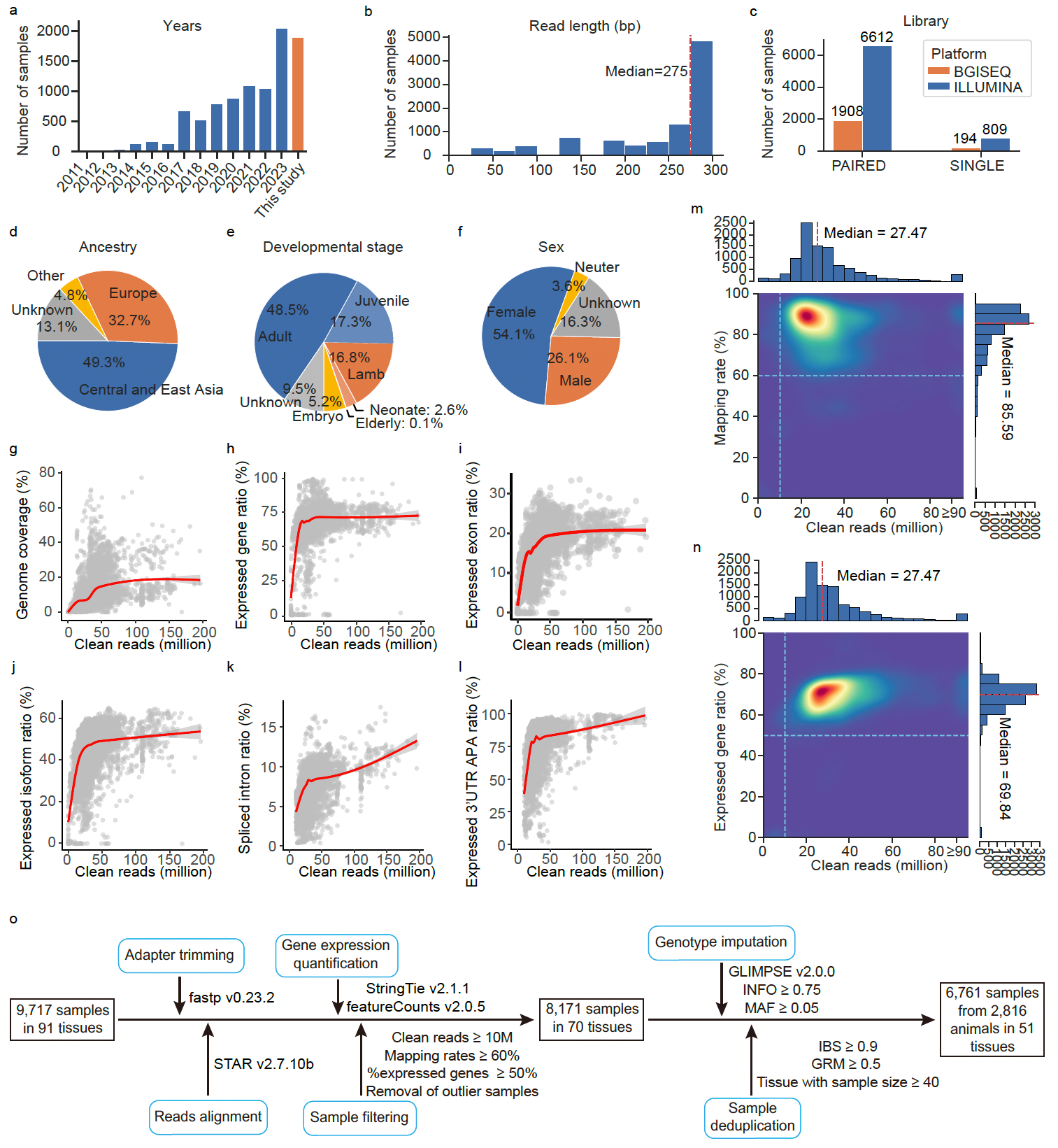

**Supplementary Fig. 1 | Summary of data from the pilot phase of the SheepGTEx project. a**, Number of public RNA-seq samples available annually from 2011 to 2023, compared with newly generated samples in this study. **b–f**, Overview of the 9,717 RNA-seq samples, including read length (**b**), sequencing platform and strategy (**c**), ancestry group (**d**), developmental stage (**e**) and sex (**f**). **g–l**, Smoothed lines (fitted using the *geom_smooth* function in ggplot2) between the number of clean reads and the ratio of genome coverage (**g**), expressed genes (Transcripts Per Million, TPM > 0.1) (**h**), expressed exons (TPM > 0.1) (**i**), expressed isoforms (TPM > 0.1) (**j**). spliced introns (**k**) and expressed 3’UTR APA (PDUI not missing) (**l**). **m–n**, Relationship between the distributions of clean read count and mapping rate (**m**) and expressed gene ratio (**n**). Dashed blue lines indicate the quality control thresholds. **o**, Workflow of RNA-seq sample processing and filtering. A total of 9,717 bulk RNA-seq samples collected across 91 tissues were processed through a unified analytical pipeline, including adapter trimming with fastp (v0.23.2)^12^, two-pass alignment to the sheep reference genome using STAR (v2.7.10b)^63^, and gene-level quantification using StringTie (v2.1.1)^64^ and featureCounts (v2.0.5)^65^. Samples were then subjected to stringent quality control, retaining only those with ≥ 10 million clean reads, uniquely mapping rates ≥ 60%, and detectable expression (TPM ≥ 0.1) in at least 50% of annotated genes, followed by removal of outlier samples based on expression clustering. This resulted in 8,171 high-quality RNA-seq samples across 70 tissues for molecular phenotype quantification. To construct the molQTL mapping cohort, RNA-seq samples were further assigned to individuals and deduplicated using genotype-based identity-by-state (IBS ≥ 0.9) and genomic relationship matrix (GRM ≥ 0.5) criteria, with only tissues containing at least 40 samples retained for downstream analyses. Genotypes were subsequently imputed using GLIMPSE2 (v2.0.0)^16^, and variants were filtered at INFO ≥ 0.75 and minor allele frequency (MAF) ≥ 0.05. The final dataset comprised 6,761 RNA-seq samples from 2,816 individuals across 51 tissues for molQTL mapping. Sample numbers retained after each filtering step are indicated.

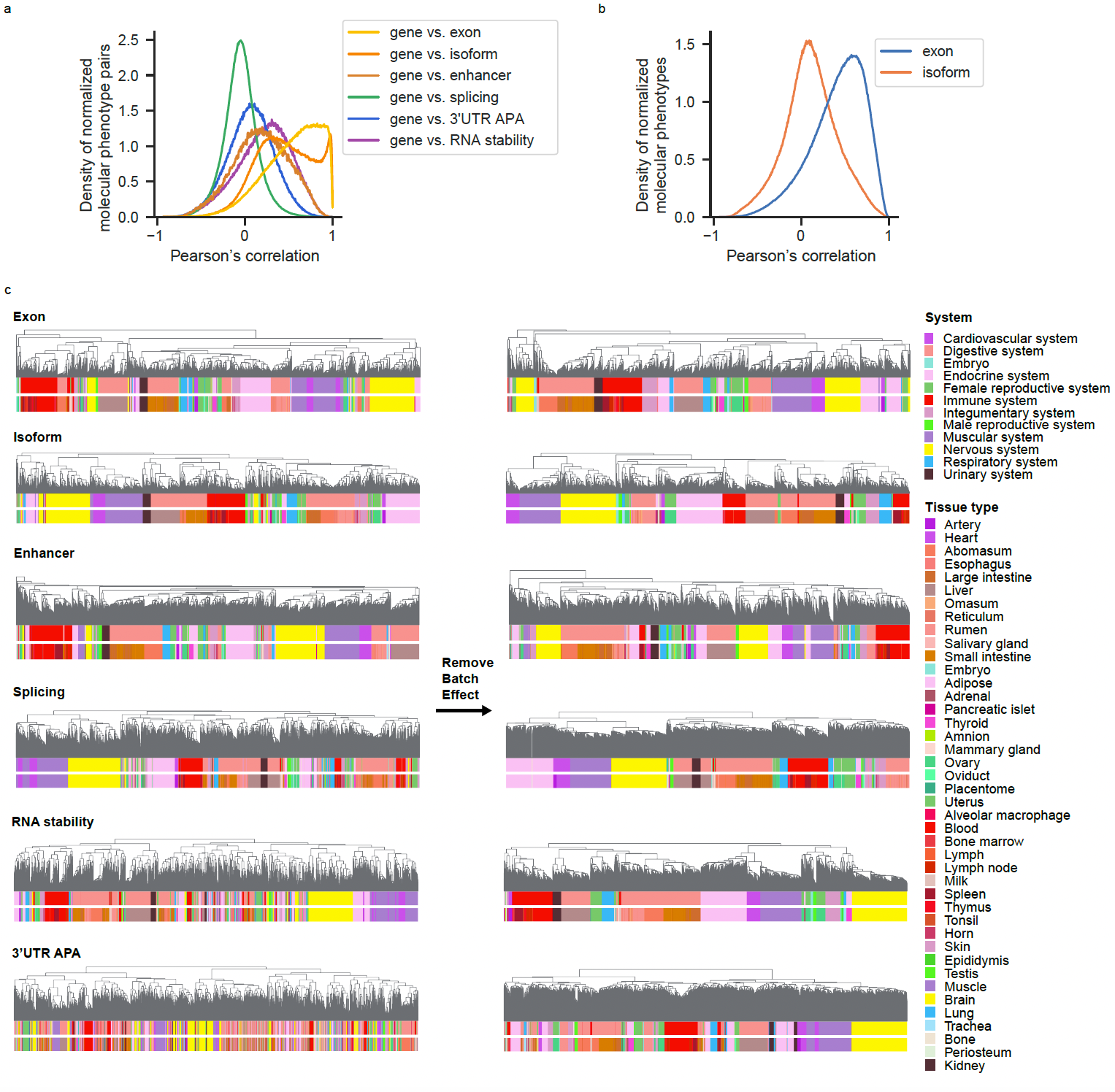

**Supplementary Fig. 2 | Characterization of molecular phenotypes.** **a**, Pearson’s correlation between gene expression and each of the other molecular phenotypes across samples, calculated for each gene within each tissue. **b**, Pearson’s correlations of exon- or isoform-level expression within each gene and tissue, computed across samples for all pairwise combinations of exons or isoforms belonging to the same gene. **c**, hierarchical clustering of six molecular phenotypes based on the expression levels of the ~20% most variable features (ranked by standard deviation). Clustering was performed using the *hclust* function in R (v4.3.0), with distances defined as (1 − *r*), where *r* denotes the Pearson’s correlation coefficient. For abundance-based molecular phenotypes, input values were log_2_(TPM + 0.25); for structure-based phenotypes, normalized values were used.

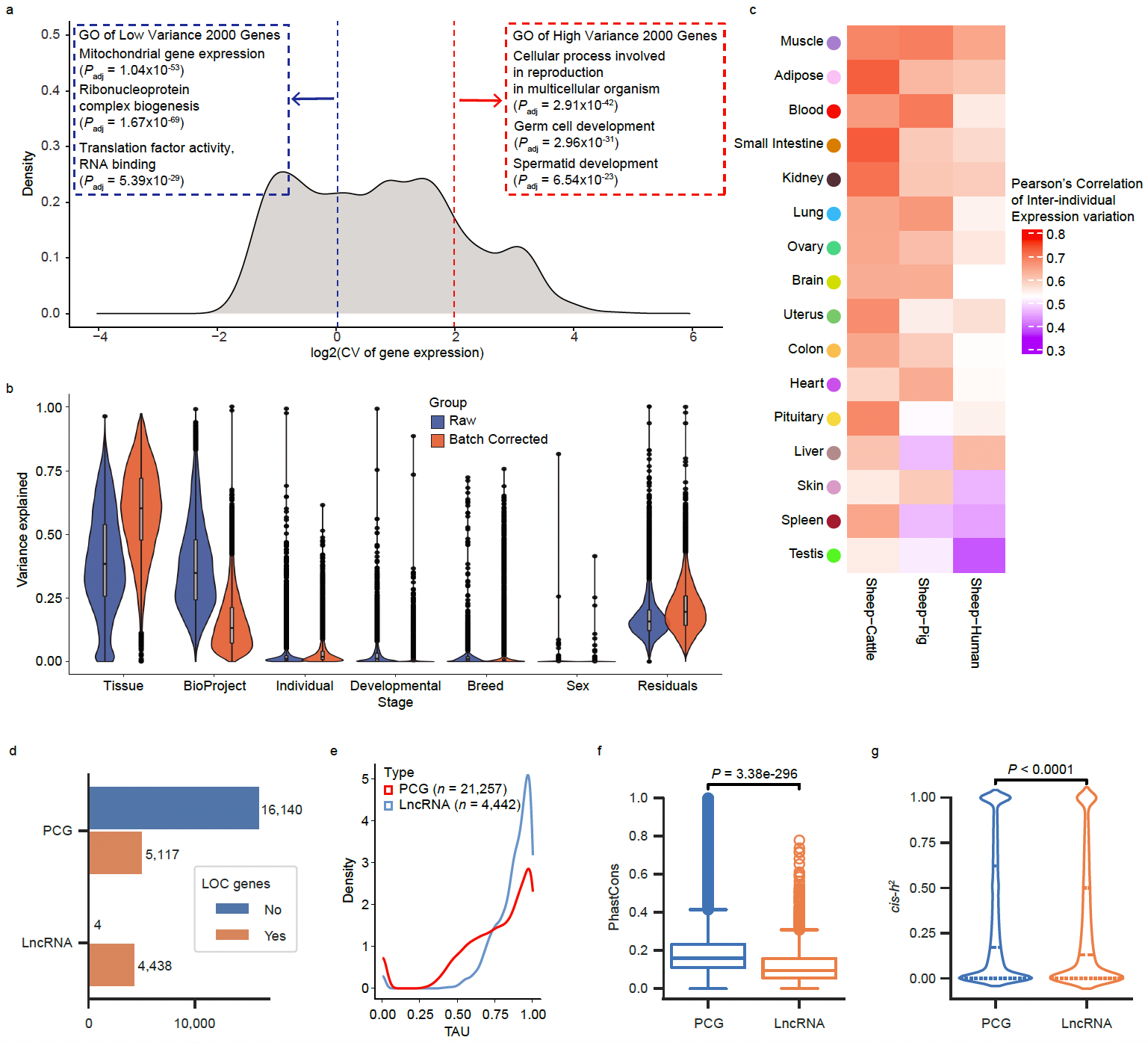

**Supplementary Fig. 3 | Decomposition of gene expression variance. a**, Distribution of the log-transformed coefficient of variation (CV) of gene expression, along with Gene Ontology (GO) terms enriched among the top 2,000 genes with the highest and lowest expression variance. GO enrichment analysis was performed using *enrichGO* function from clusterProfiler package (v4.8.3)^2^ with the org.Hs.eg.db (v3.16.0) database. **b**, Proportion of gene expression variance explained by different sources, before and after batch effect correction. Variance components were estimated using a linear mixed model (LMM) implemented in the variancePartition package (v1.30.2)^1^. **c**, Pearson’s correlation of inter-individual gene expression variability between sheep and cattle, sheep and pig, and sheep and human. **d**, Numbers of annotated genes and uncharacterized (LOC) genes in NCBI *Ovis aries* Annotation Release 104 (ARS-UI_Ramb_v2.0^55^, GCF_016772045.1). **e**, Tissue-specific expression of protein-coding genes (PCGs) and long non-coding RNA genes (lncRNAs), as measured by the TAU score. **f**, Evolutionary constraint of PCGs and lncRNAs, as measured by PhastCons scores. **g**, *cis*-heritability (*cis*-*h*^2^) of gene expression for PCGs and lncRNAs.

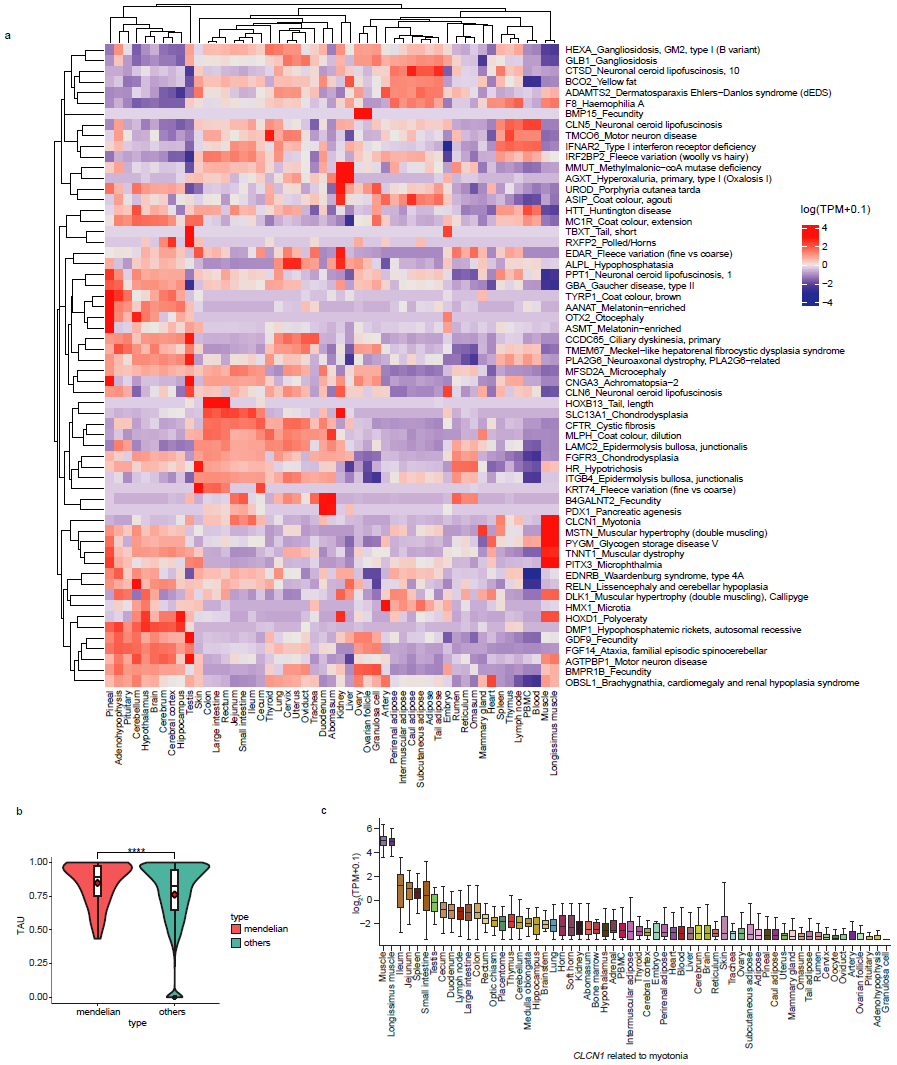

**Supplementary Fig. 4 | Expression of genes associated with Mendelian traits in sheep.** Mendelian traits annotations were obtained from Online Mendelian Inheritance in Animals (OMIA)^7^. **a**, Clustered heatmap showing the tissue-specific expression patterns of Mendelian trait-associated genes across different traits. **b**, Comparison of tissue specificity (measured by TAU scores) between Mendelian genes and all other genes. **c**, Box plot illustrating an example of the *CLCN1* gene with relatively high expression in tissues relevant to its associated phenotype.

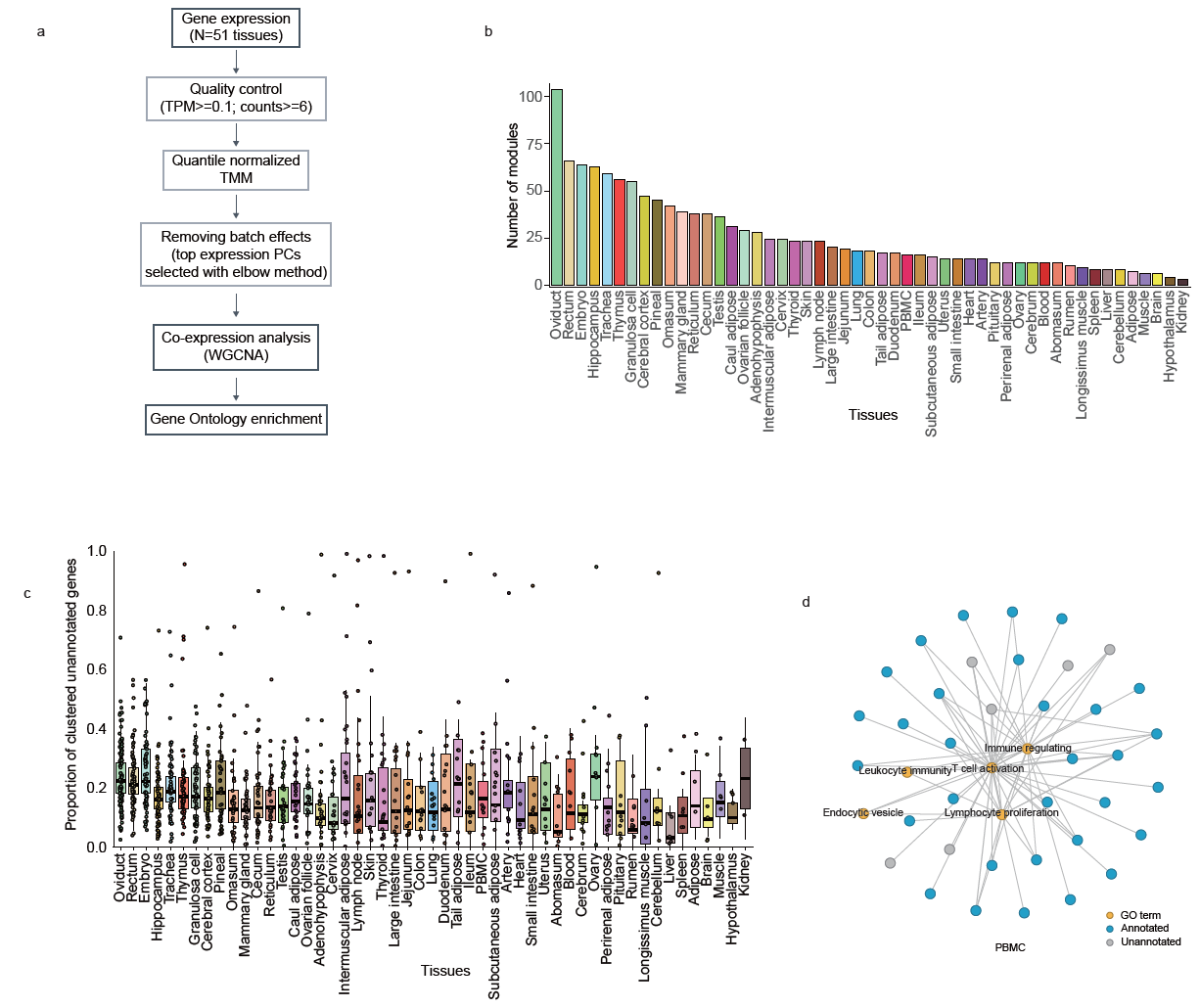

**Supplementary Fig. 5 | Gene co-expression** **analysis. a**, Overview of the workflow for gene co-expression analysis. **b**, Number of co-expression modules identified across 51 tissues. **c**, Proportion of genes within co-expression modules lacking functional annotation in the Gene Ontology (GO) database across tissues. **d**, Representative gene co-expression module identified in peripheral blood mononuclear cells (PBMC).

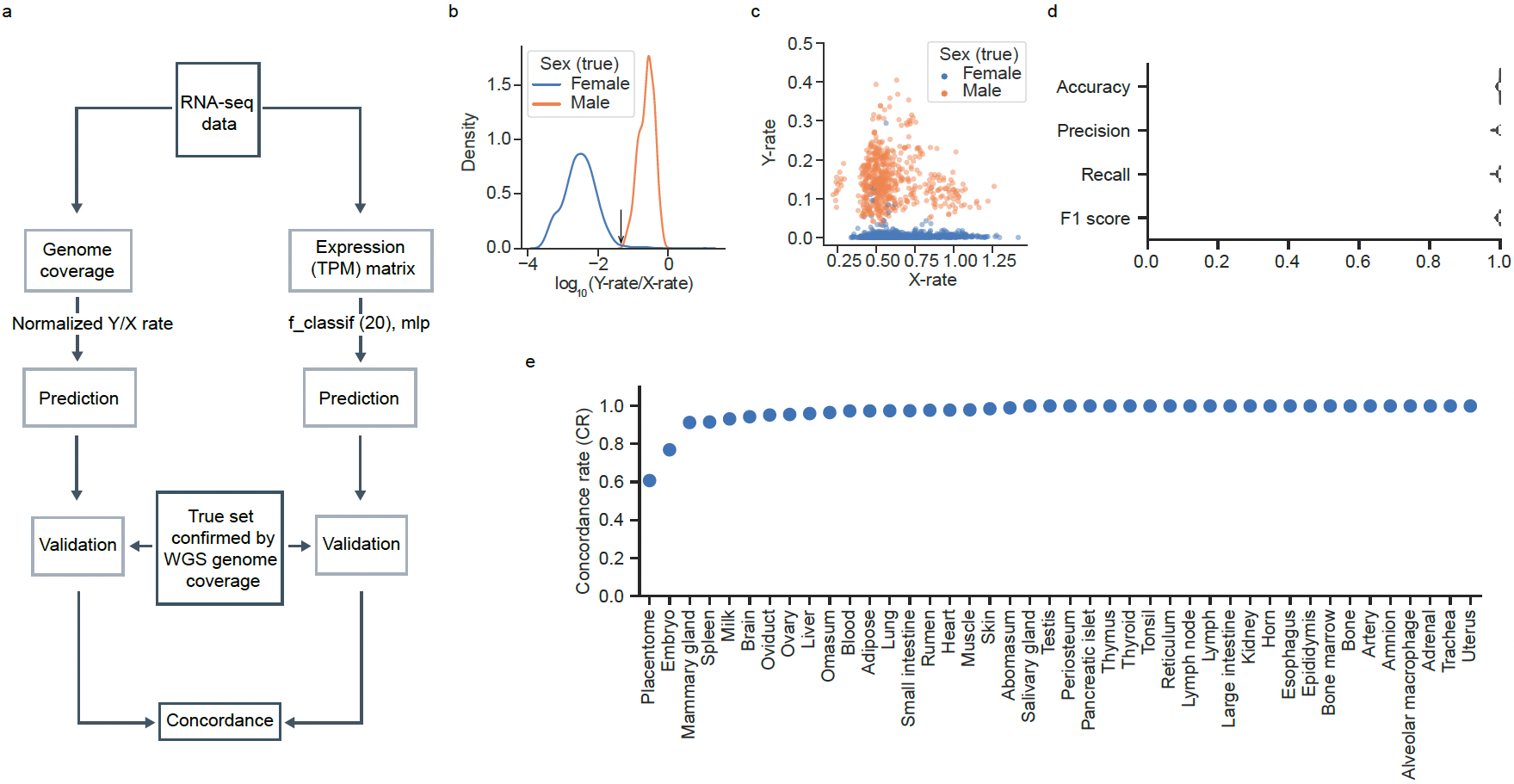

**Supplementary Fig. 6 | Sex prediction of RNA-seq samples. a**, Overview of the sex prediction workflow. **b**, Density plot showing the distribution of log_10_-transformed X-rate/Y-rate in samples with known sex. X-rate and Y-rate were calculated as the ratio of the mean sequencing depth of SNPs on the chromosome X or Y to that on the autosomes. **c**, Distribution of the X-rate and Y-rate in the true sex-labeled dataset. **d**, Performance of the gene expression-based sex prediction method, evaluated using accuracy, precision, recall, and F1 score. **e**, Concordance rate between genome coverage-based and gene expression-based methods across tissues.

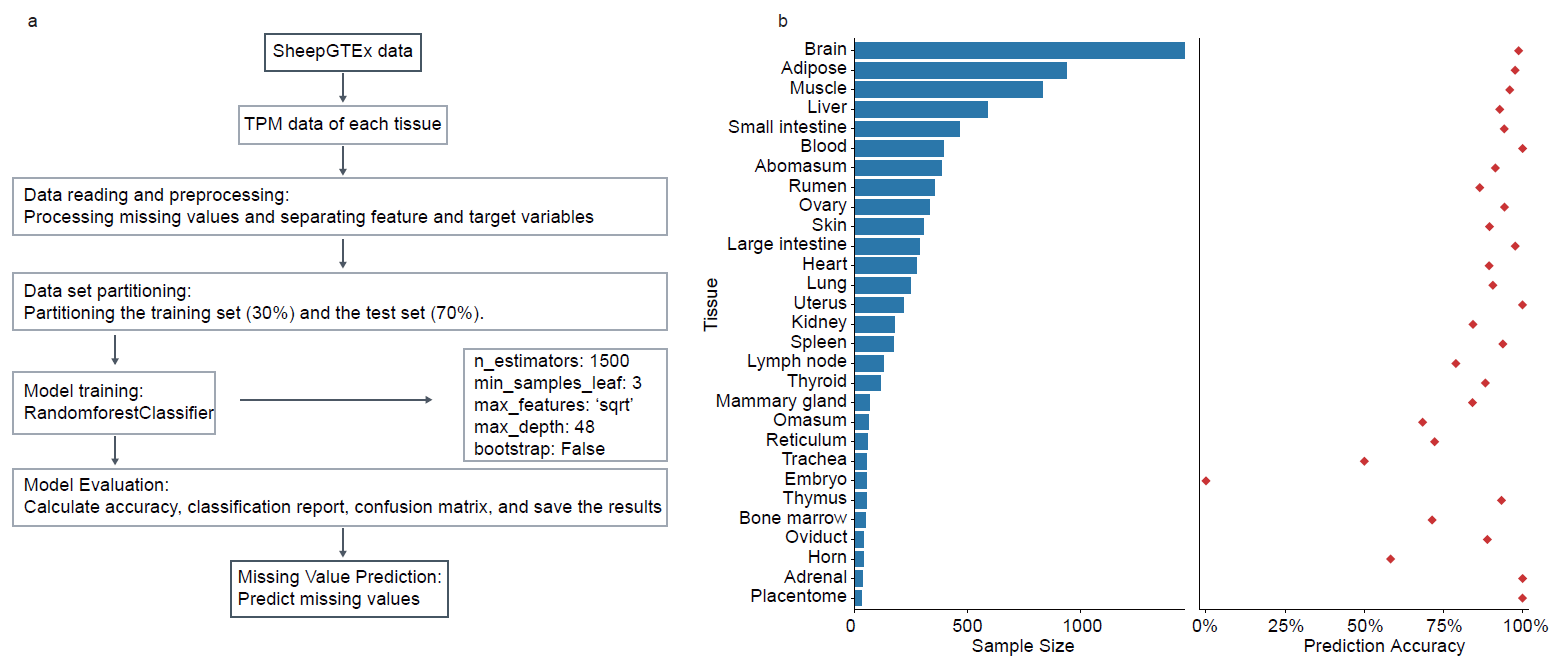

**Supplementary Fig. 7 |** **Developmental stage prediction. a**, Workflow for predicting the developmental stage of RNA-seq samples using gene expression profiles. **b**, Sample size and corresponding prediction accuracy across tissues.

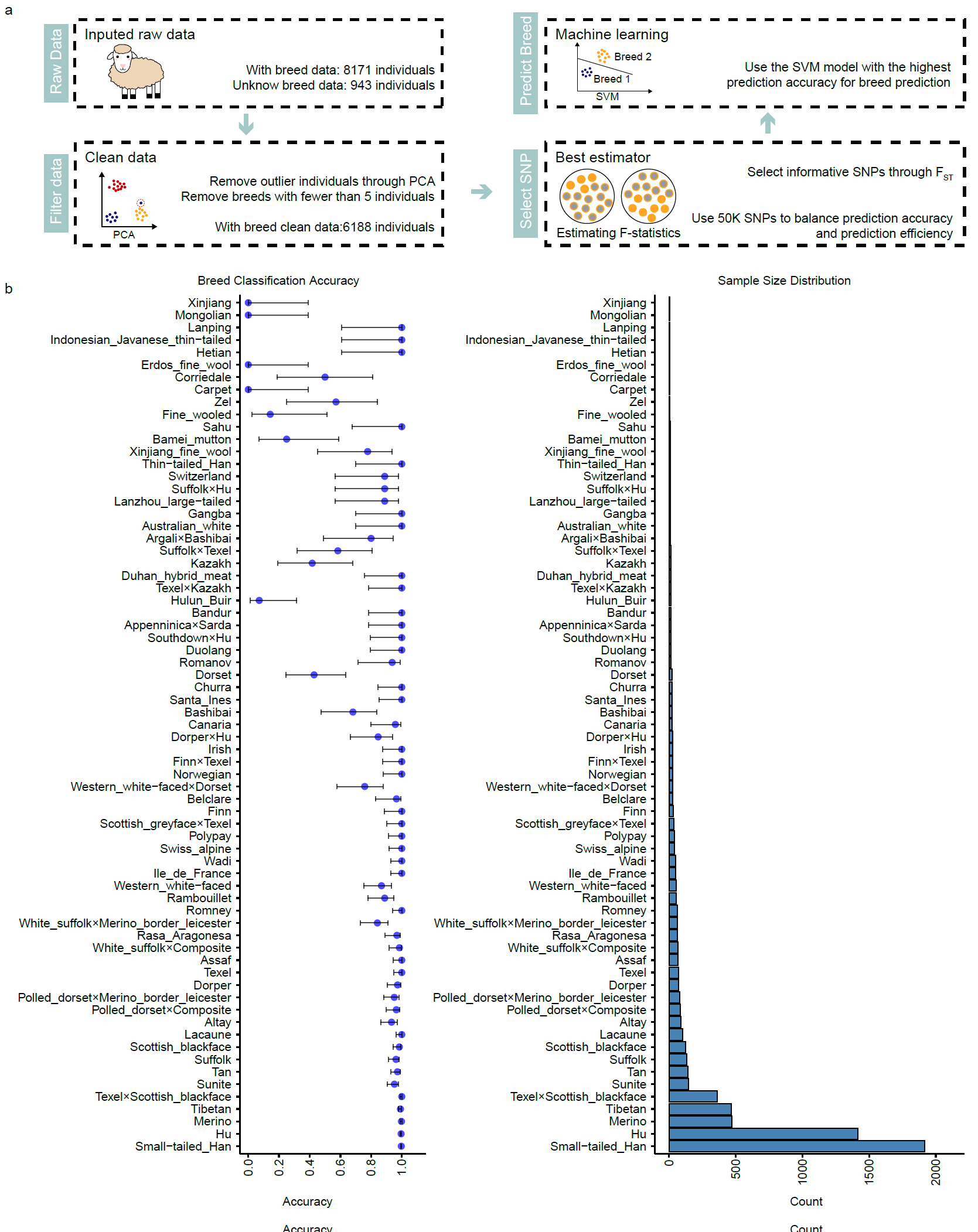

**Supplementary Fig. 8 | Breed prediction. a**, Workflow for predicting sheep breeds based on genotypes imputed from RNA-seq data. **b**, Sample size and corresponding prediction accuracy across breeds. Point, mean; error bar, 95% confidence interval.

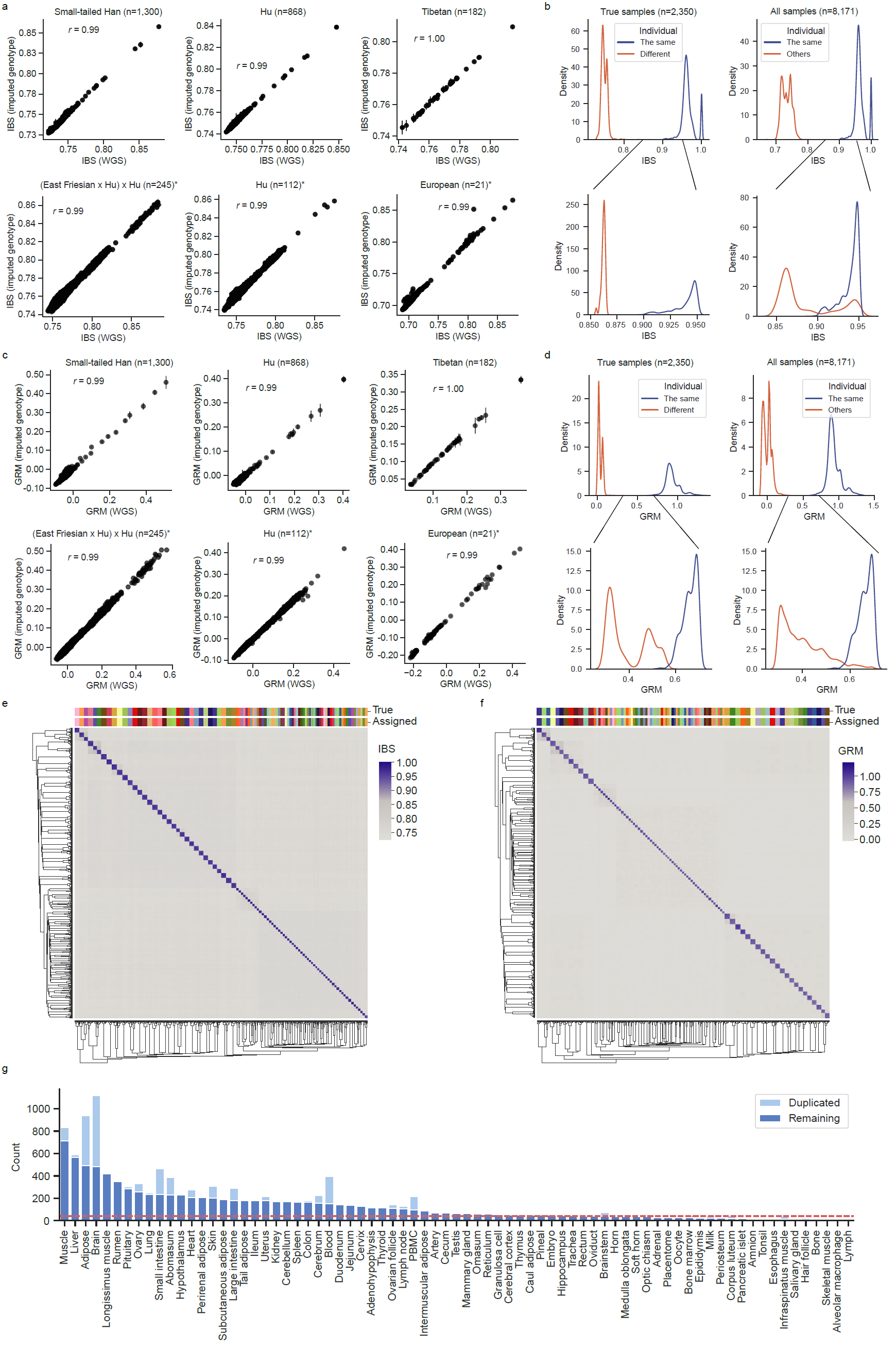

**Supplementary Fig. 9 | Identification of duplicate samples and assignment of RNA-seq samples to individuals. a**, Pearson’s correlation between identity-by-state (IBS) values derived from imputed RNA-seq genotypes and called WGS genotypes across six datasets. **b**, Distribution of IBS scores among RNA-seq samples with matched WGS individuals (left) and all RNA-seq samples (right). **c**, Pearson’s correlation between genetic relationship matrices (GRM) derived from imputed RNA-seq genotypes and called WGS genotypes across six datasets. **d**, Distribution of GRM values in matched RNA-WGS pairs (left) and all RNA-seq samples (right). **e–f**, Clustered heatmap of IBS (**e**) and GRM (**f**) for 2,350 RNA-seq samples across 93 individuals. Top color bars indicate true individual labels (True) and clustering-based assignments (Assigned). **g**, Number of duplicated and remaining individuals across 75 tissues. Sample pairs with IBS ≥ 0.9 and GRM ≥ 0.5 were classified as duplicates. The red dashed line indicates the minimum sample size (*n* = 40) required for molQTL mapping.

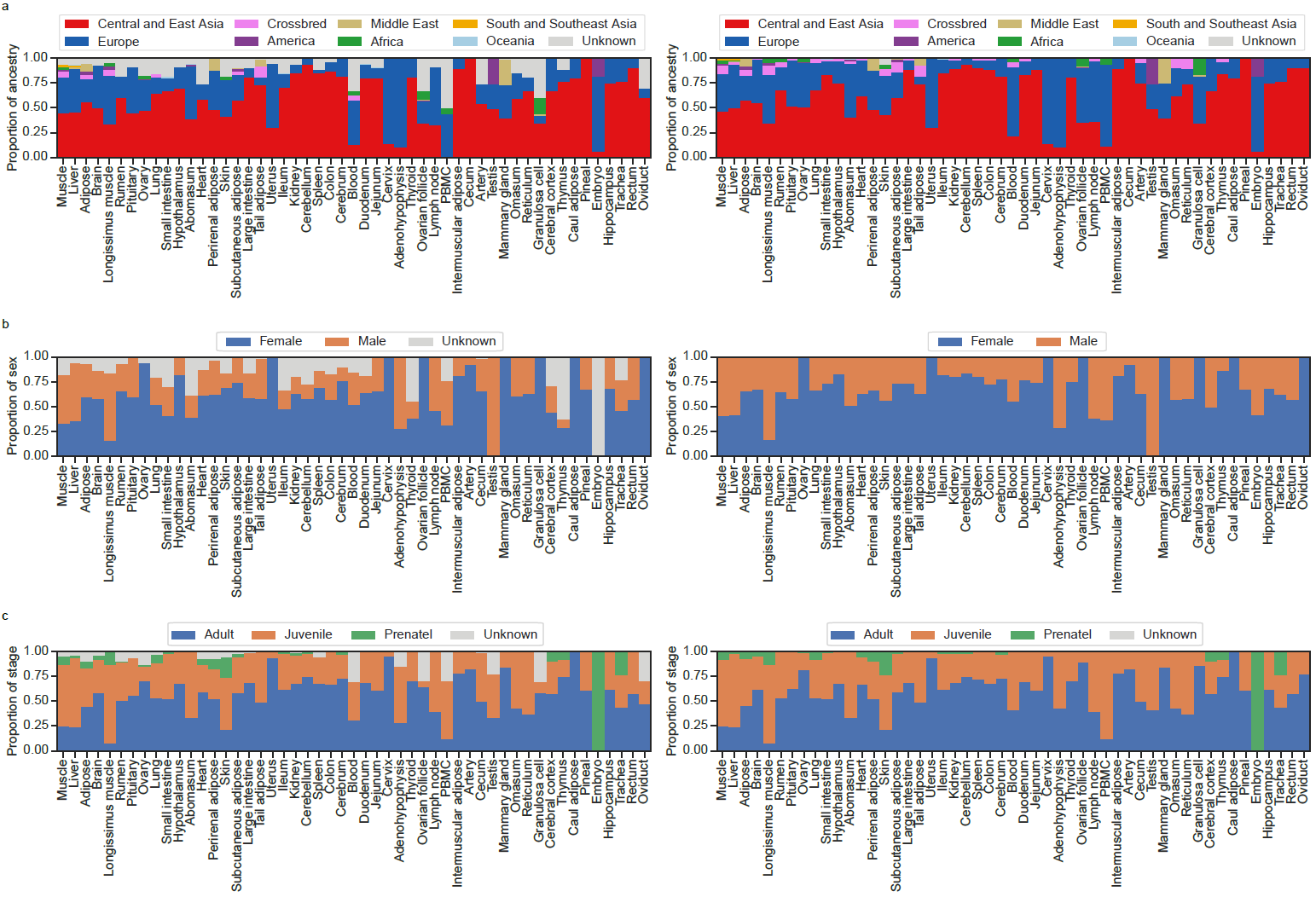

**Supplementary Fig. 10 | Summary of ancestry, sex and developmental stage of all 2,816 RNA-seq individuals across tissues. a**, Proportions of ancestries inferred based on geographic origins across 51 tissues before (left) and after (right) imputation of missing data. **b**, Proportions of sexes across 51 tissues before (left) and after (right) imputation of missing data. **c**, Proportions of developmental stages across 51 tissues before (left) and after (right) imputation of missing data.

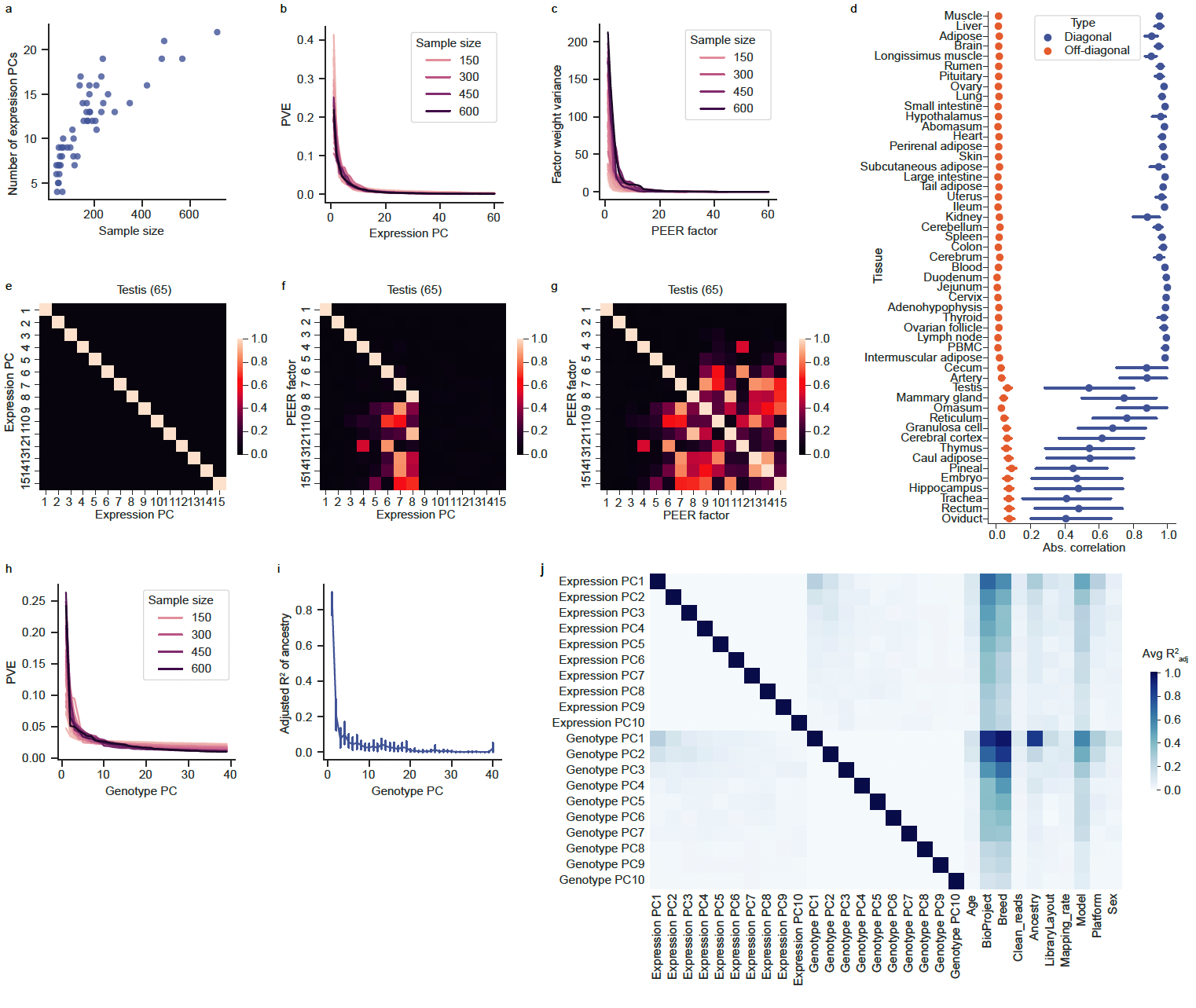

**Supplementary Fig. 11 |** **Confounding factors in RNA-seq data. a**, Number of estimated gene expression principal components (PCs) as a function of sample size. **b**, Proportion of variance explained (PVE) by each expression PC across 51 tissues. **c**, Variance of factor weights for probabilistic estimation of expression residual (PEER) factors across 51 tissues. **d**, Absolute Pearson’s correlation between PEER factors and expression PCs across 51 tissues, with matrix entries classified as diagonal or off-diagonal. **e–g**, Correlation matrices between expression PCs and PEER factors in Testis (n=65). **h**, PVE by genotype PCs across 51 tissues. **i**, Proportion of variance (adjusted R^2^) of ancestry (see **Extended Data Fig. 2a**) explained by genotype PCs across 51 tissues, estimated using linear modeling in R. **j**, Mean adjusted R^2^ of the top 10 expression and genotype PCs, as well as known batch effects (e.g., age and sex) across 51 tissues.

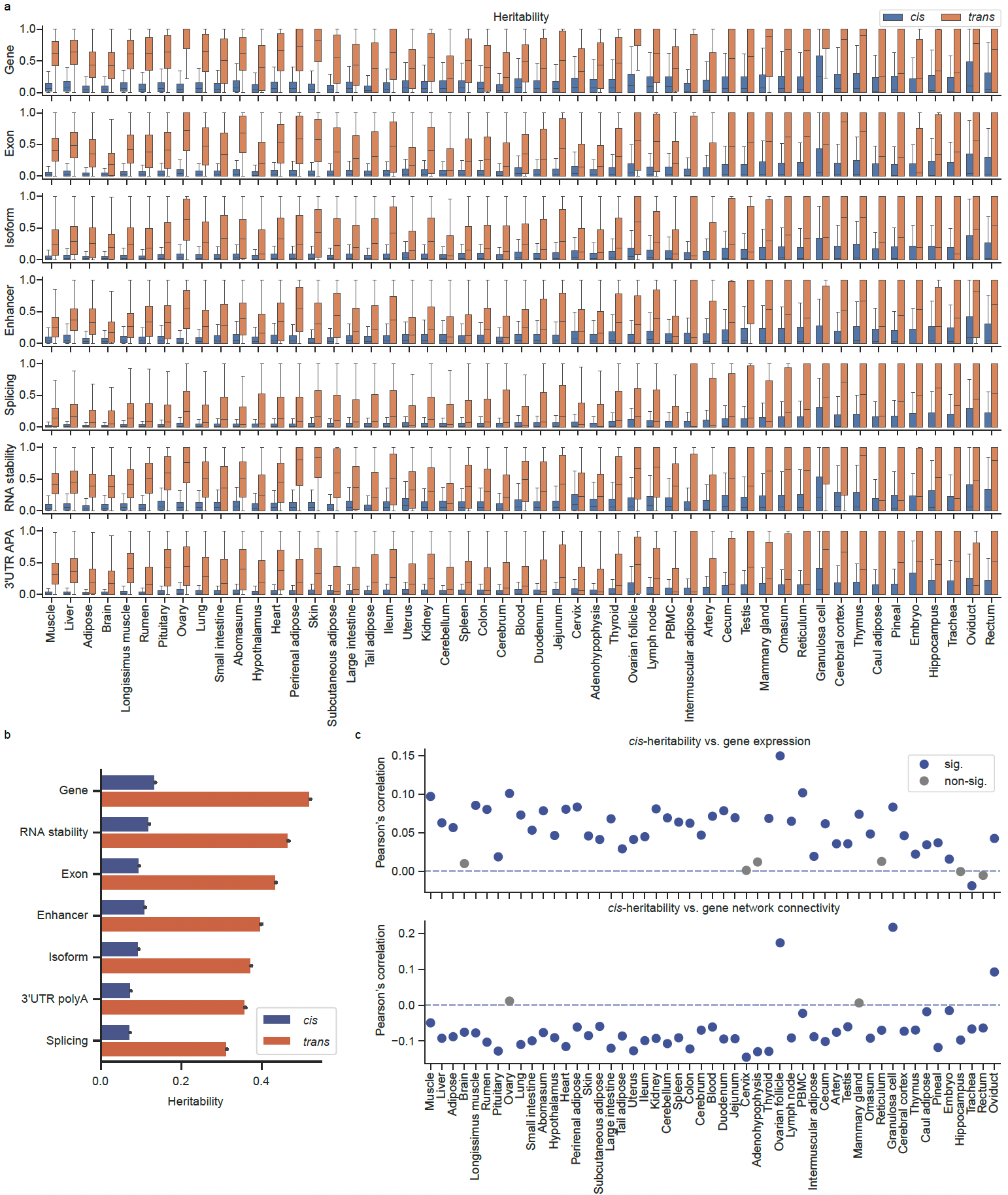

**Supplementary Fig. 12 | Heritability of seven molecular phenotypes across 51 sheep tissues. a–b**, Distributions of estimated *cis*- and *trans*-heritability across seven molecular phenotype classes and tissues. Tissues in **a** are ordered by decreasing sample size. **c**, Pearson’s correlations between gene *cis*-heritability and gene expression level (top) and gene network connectivity (bottom), the latter measured by the absolute value of module eigengene-based connectivity (|kME|). Statistical significance was defined as *P* < 0.05.

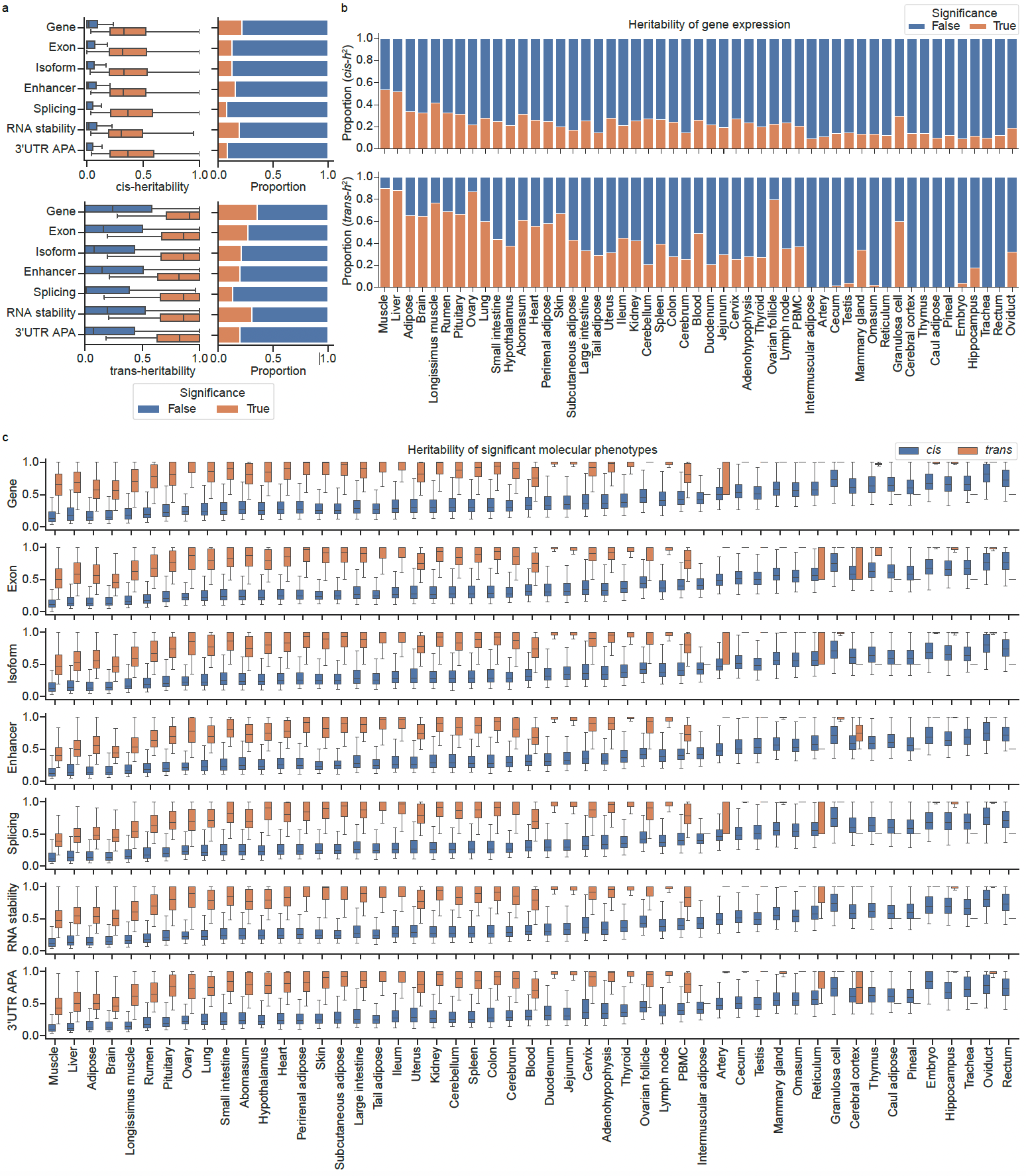

**Supplementary Fig. 13 | Comparison of heritability estimates of significant molecular phenotypes across 51 sheep tissues. a**, Distributions and proportions of significant and non-significant *cis*- and *trans*-heritability across seven types of molecular phenotypes. The statistical significance of heritability was determined by one-sided Wald-test with *P* < 0.05. **b**, Proportion of significant and non-significant *cis*- and *trans*-heritability for gene expression across tissues, ordered by decreasing sample size. **c**, Distributions of significant *cis*- and *trans*-heritability across seven molecular phenotypes and 51 tissues.

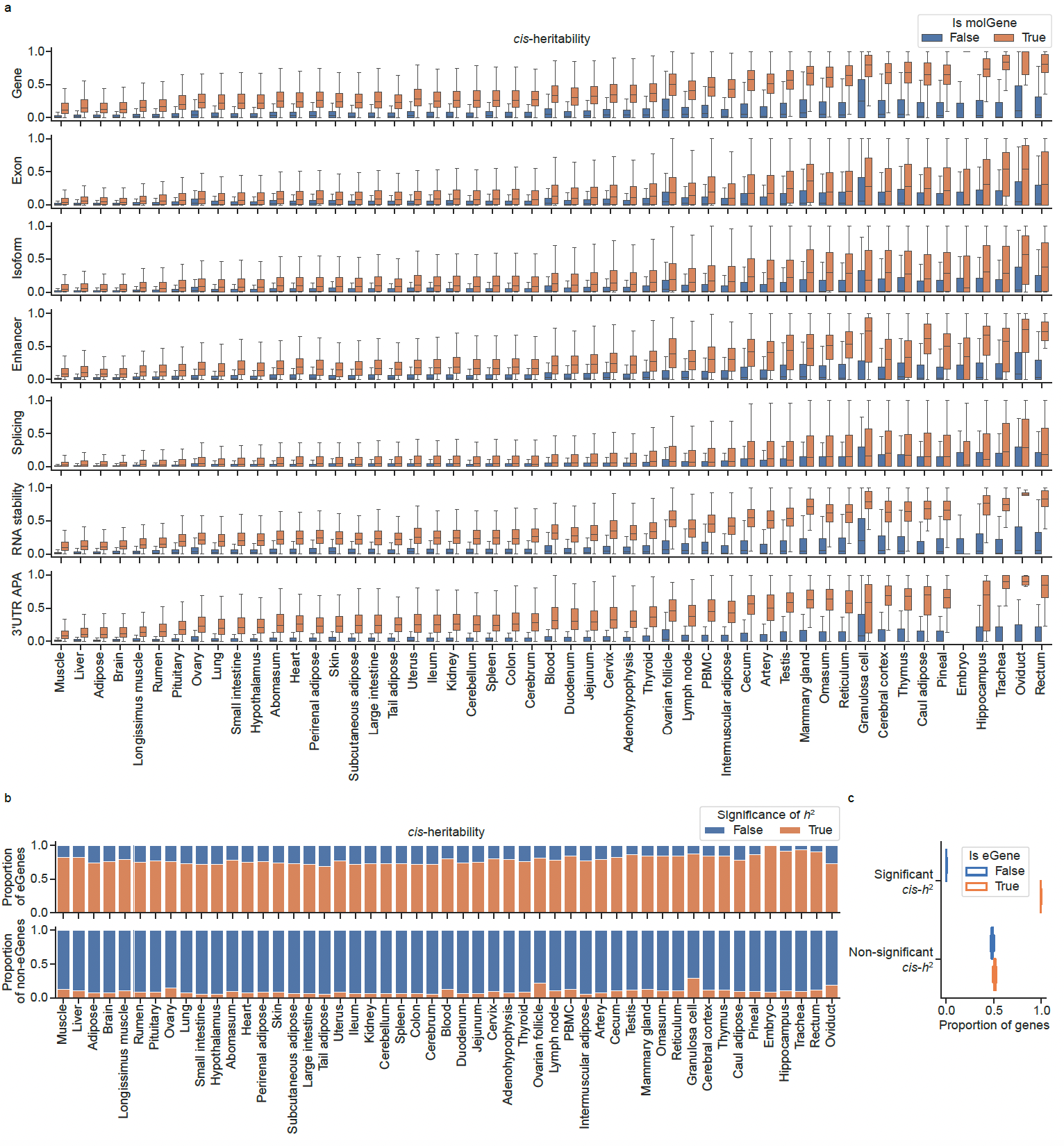

**Supplementary Fig. 14 | Comparison of heritability estimates between molGenes and non-molGenes across 51 sheep tissues. a**, Distributions of estimated *cis*-heritability for seven types of molGenes and non-molGenes across tissues. **b**, Proportions of estimated significant and non-significant *cis*-heritability for eGenes and non-eGenes across tissues. **c**, Proportion of true eGenes and non-eGenes with significant or non-significant *cis*-heritability based on 50 simulations using a dataset of 300 individuals from a previous study^33^.

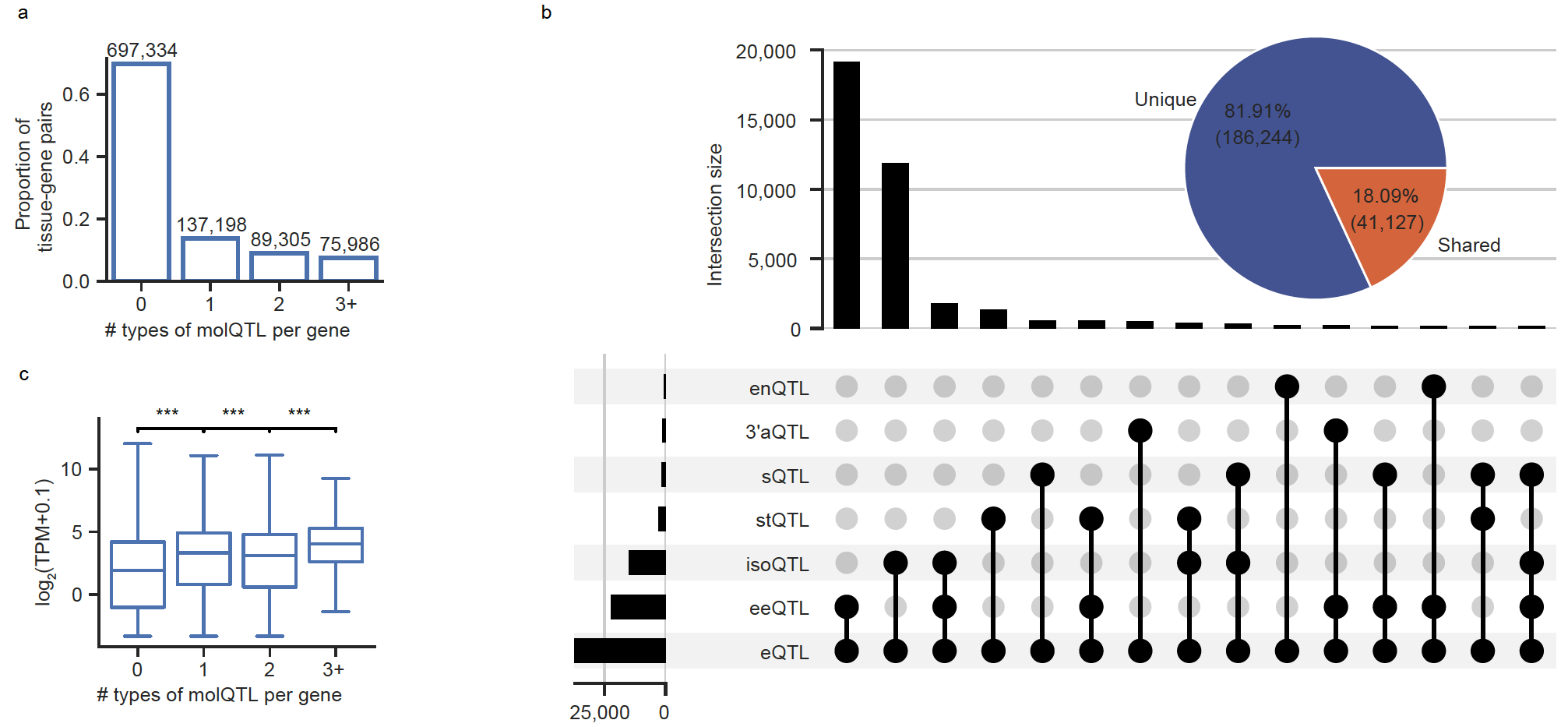

**Supplementary Fig. 15 | Shared and unique regulatory effects of different molecular phenotypes in the same genes. a**, Proportion of genes associated with different numbers of molQTL types. **b**, Number of lead eQTL that are shared with or unique from other types of lead molQTL within the same genes. The pie chart summarizes the overall proportion of shared vs. unique lead eQTL-molQTL pairs, while the upset plot illustrates the top 15 most frequent combinations of shared lead variants. **c**, Boxplots showing median gene expression levels of each tissue stratified by the number of associated molQTL types. *P* values were calculated using the two-sided Mann-Whitney U test. ns: non-significant; *: *P* < 0.05; **: *P* < 0.01; ***: *P* < 0.001.

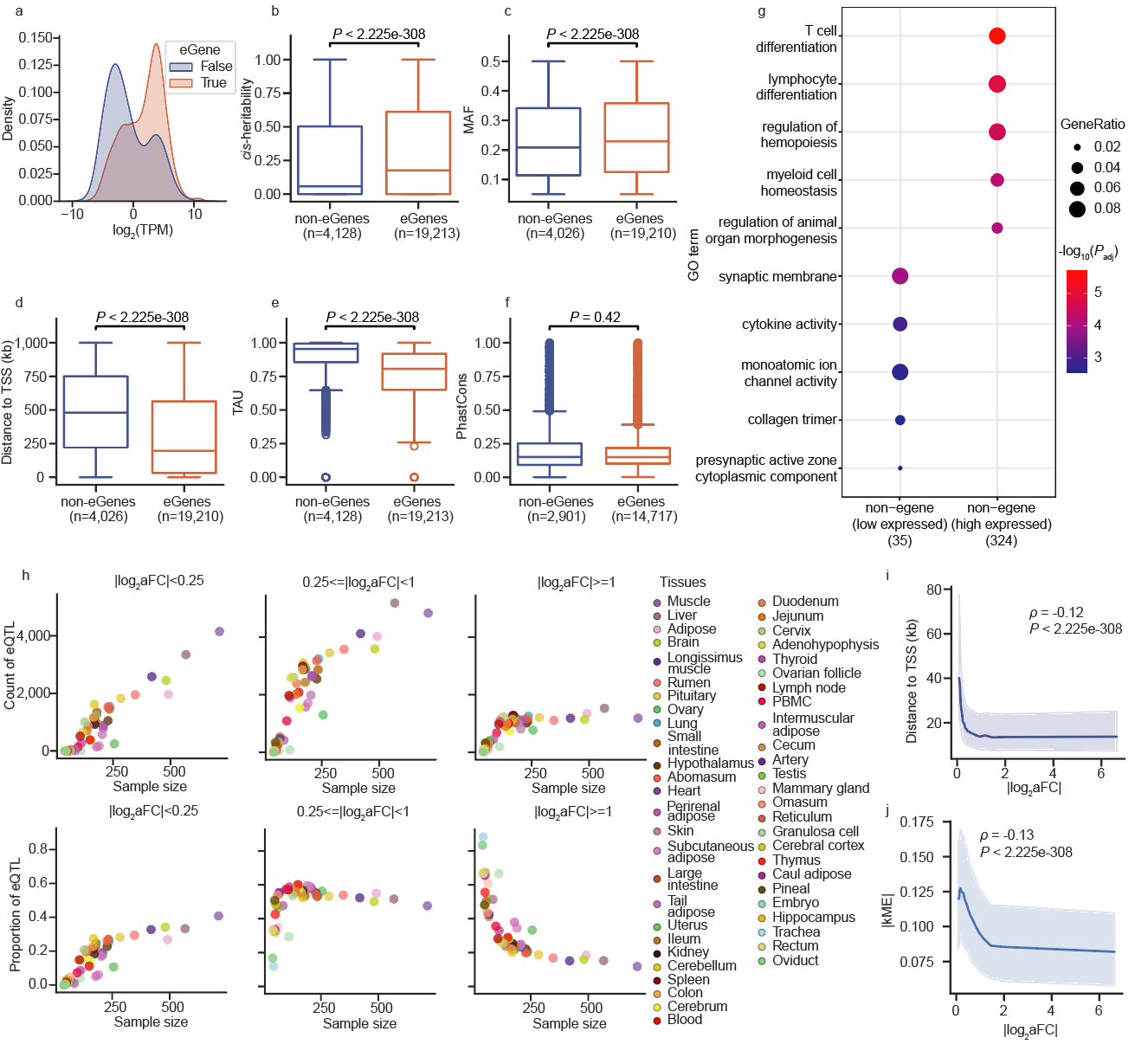

**Supplementary Fig. 16 | Characterization of *cis*-eQTL across 51 sheep tissues. a**, Gene expression levels of eGenes (genes with at least one significant eQTL in any tissue) compared to non-eGenes across all 51 tissues. **b–f**, Comparison between eGenes and non-eGenes for: **b**, *cis*-heritability; **c**, minor allele frequency (MAF) of lead variants; **d**, distance of lead variants to transcription start sites (TSS); **e**, tissue specificity (measured by TAU score); **f,** sequence constraint (PhastCons score). Statistical significance was assessed using the two-sided Mann-Whitney U test. **g**, Top GO term enrichment among non-eGenes corresponding to the two expression peaks observed in (**a**), where genes within ±1.5 log_2_(TPM) of each peak were included in the analysis (280 genes for the high-expression peak and 242 genes for the low-expression peak). **h**, Number (top) and proportion (bottom) of detected eQTL across bins of increasing effect size (measured as absolute log₂ allelic fold change, |log_2_(aFC)|) as a function of tissue sample size. **i**, Spearman’s correlation between effect sizes (|log_2_(aFC)|) and distances to TSS of lead eQTL. **j**, Spearman’s correlation between absolute effect sizes (|log_2_(aFC)|) of eQTL and gene network connectivity (|kME|) of target genes.

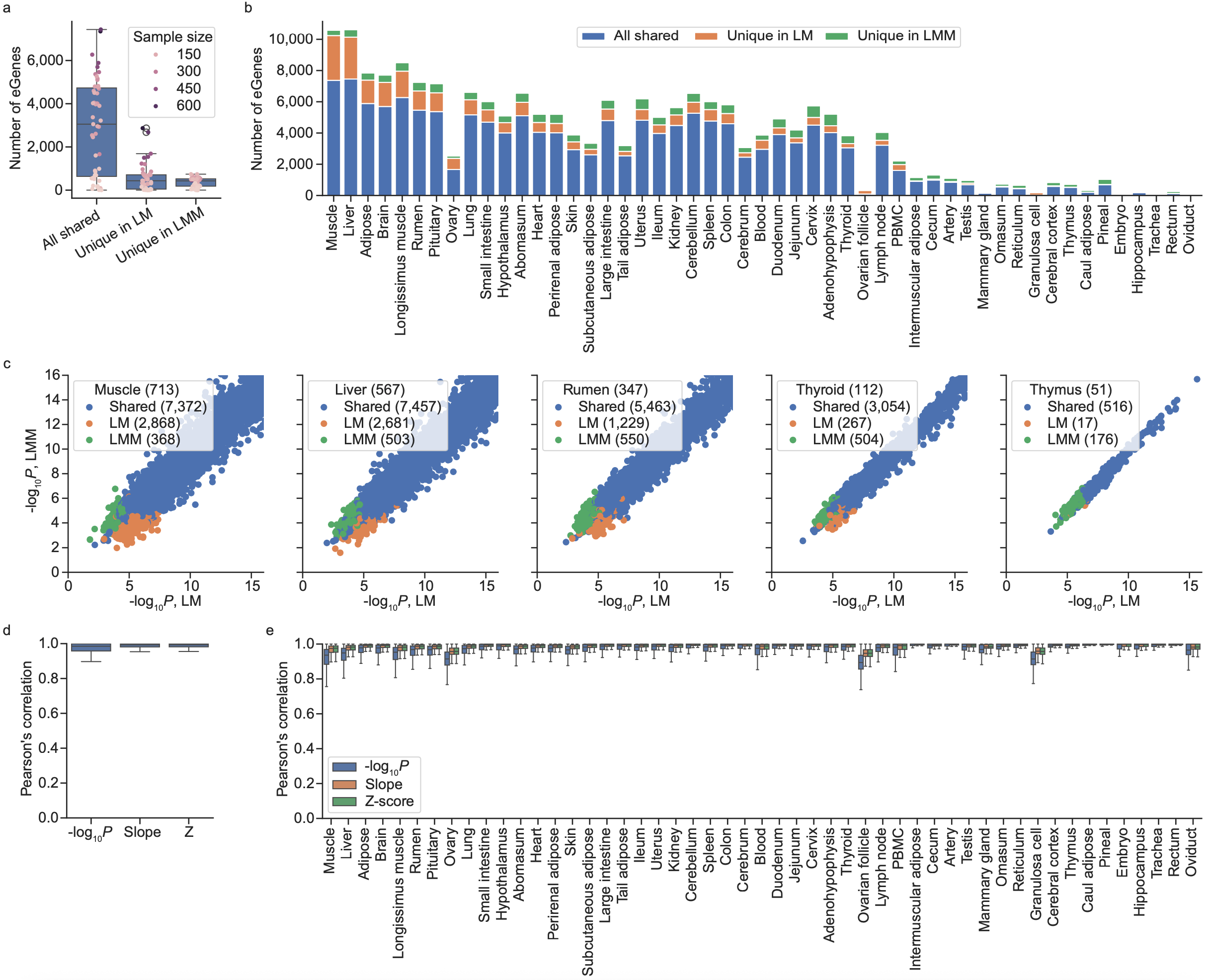

**Supplementary Fig. 17 | Comparison of *cis*-eQTL detection using** **linear regression model (LM) and linear mixed model (LMM). a–b**, Number of unique and shared eGenes identified by LM and LMM across 51 tissues. **c**, Comparison of nominal *P*-values (−log_10_ scale) of lead eQTL for eGenes categorized as shared between models, unique to LM, or unique to LMM, across five representative tissues with distinct sample size. **d–e**, Pearson’s correlation of nominal *P*-value (−log_10_ scale), effect size (Slope), and Z-score between LM-based and LMM-based *cis*-eQTL mapping.

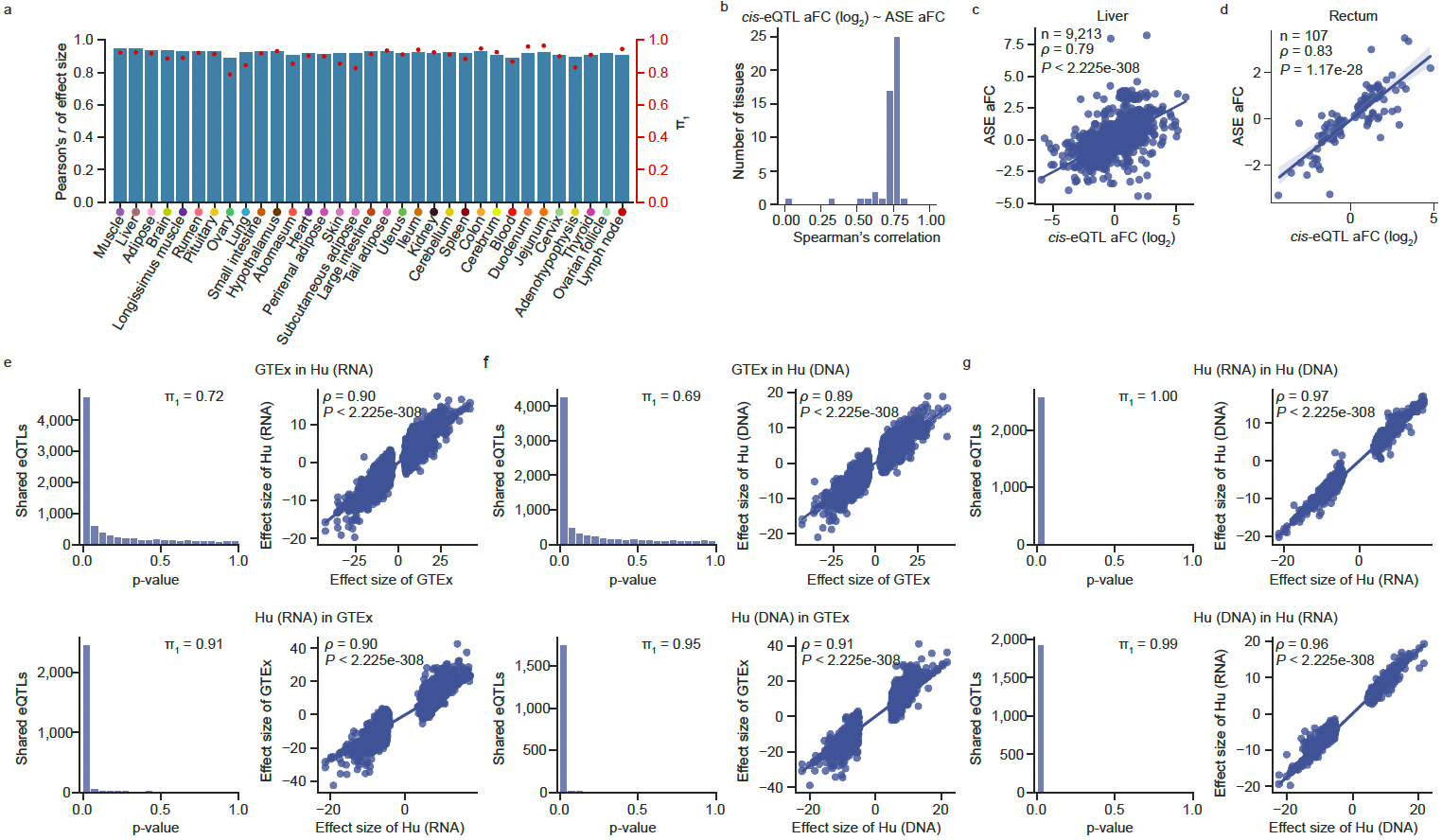

**Supplementary Fig. 18 | Validation of** ***cis*-eQTL. a**, Internal validation of *cis*-eQTL. We selected 33 tissues with sample sizes > 100 and randomly split samples into equally sized discovery and validation cohorts. eQTL mapping was performed independently in each cohort. Bars show Pearson’s correlation of normalized effect sizes between the two groups, and red dots represent Storey’s π_1_ statistics, indicating the replication rate of *cis*-eQTL. **b**–**d**, Spearman’s correlation between *cis*-eQTL effect sizes (allelic fold change, log₂ scale) and corresponding effect sizes derived from allele-specific expression (ASE) analysis. **e–g**, External validation of *cis*-eQTL using an independent Hu sheep population (*n* = 112). eQTL mapping was conducted using both imputed RNA-seq genotypes [Hu (RNA)] and WGS-based genotypes [Hu (DNA)]. Storey’s π_1_ statistics and Spearman’s correlation of normalized effect sizes were used to assess replication between GTEx discovery data and Hu (RNA) (**e**), GTEx and Hu (DNA) (**f**), and Hu (RNA) and Hu (DNA) (**g**).

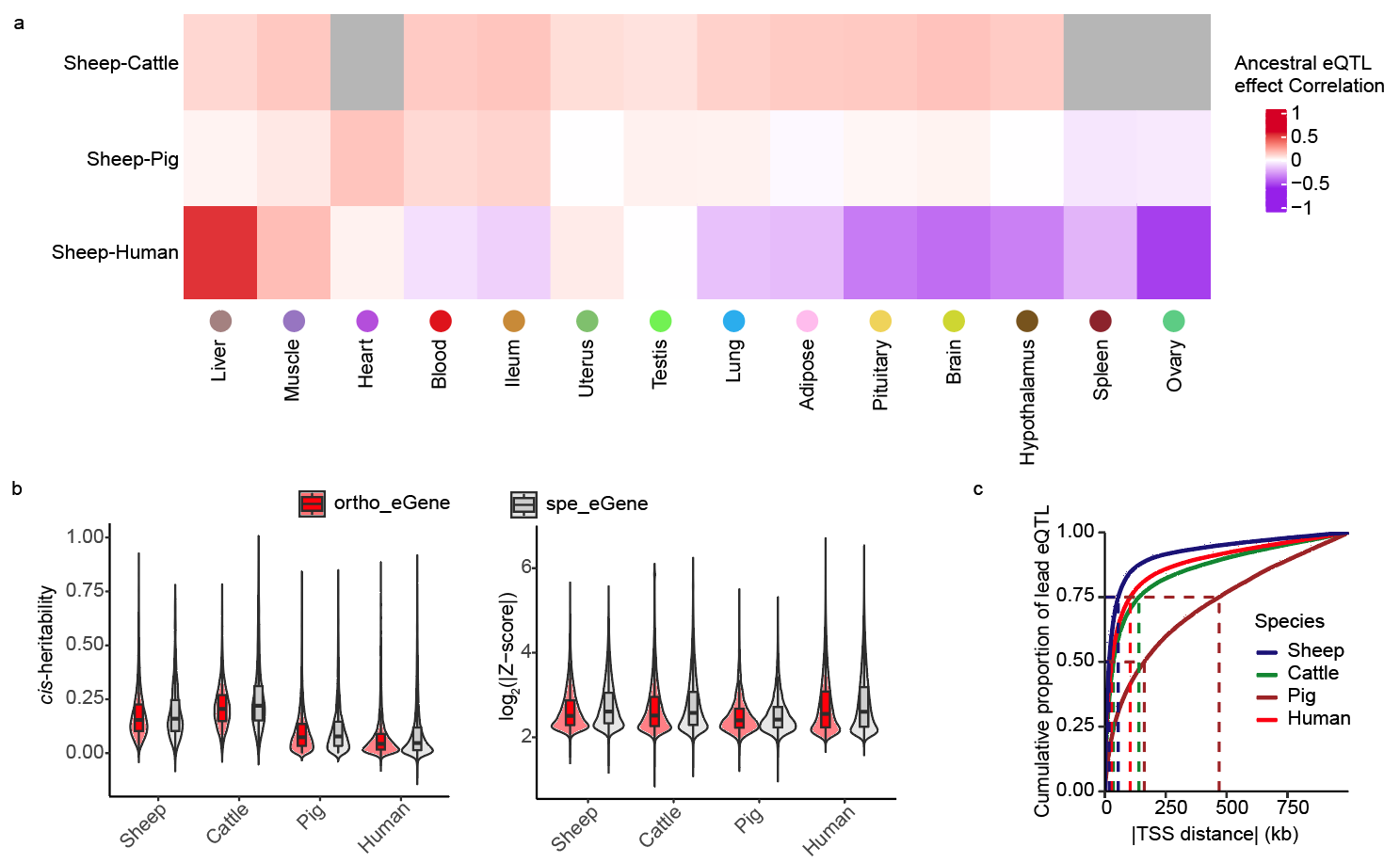

**Supplementary Fig. 19 | Cross-species comparison of regulatory effects based on ancestral alleles. a**, Pearson’s correlation of ancestral allele pairs between sheep and other species across various tissues. Grey cells indicate that no effect size (Z-score) of the ancestral allele pair was available for that tissue. **b**, comparison of *cis*-*h*^2^ and log_2_(|Z-score|) between orthologous and species-specific eGenes in 4 species. **c**, Cumulative distribution of distances between lead eQTL and transcriptional start site (TSS) of their target genes in sheep, cattle, pig, and human.

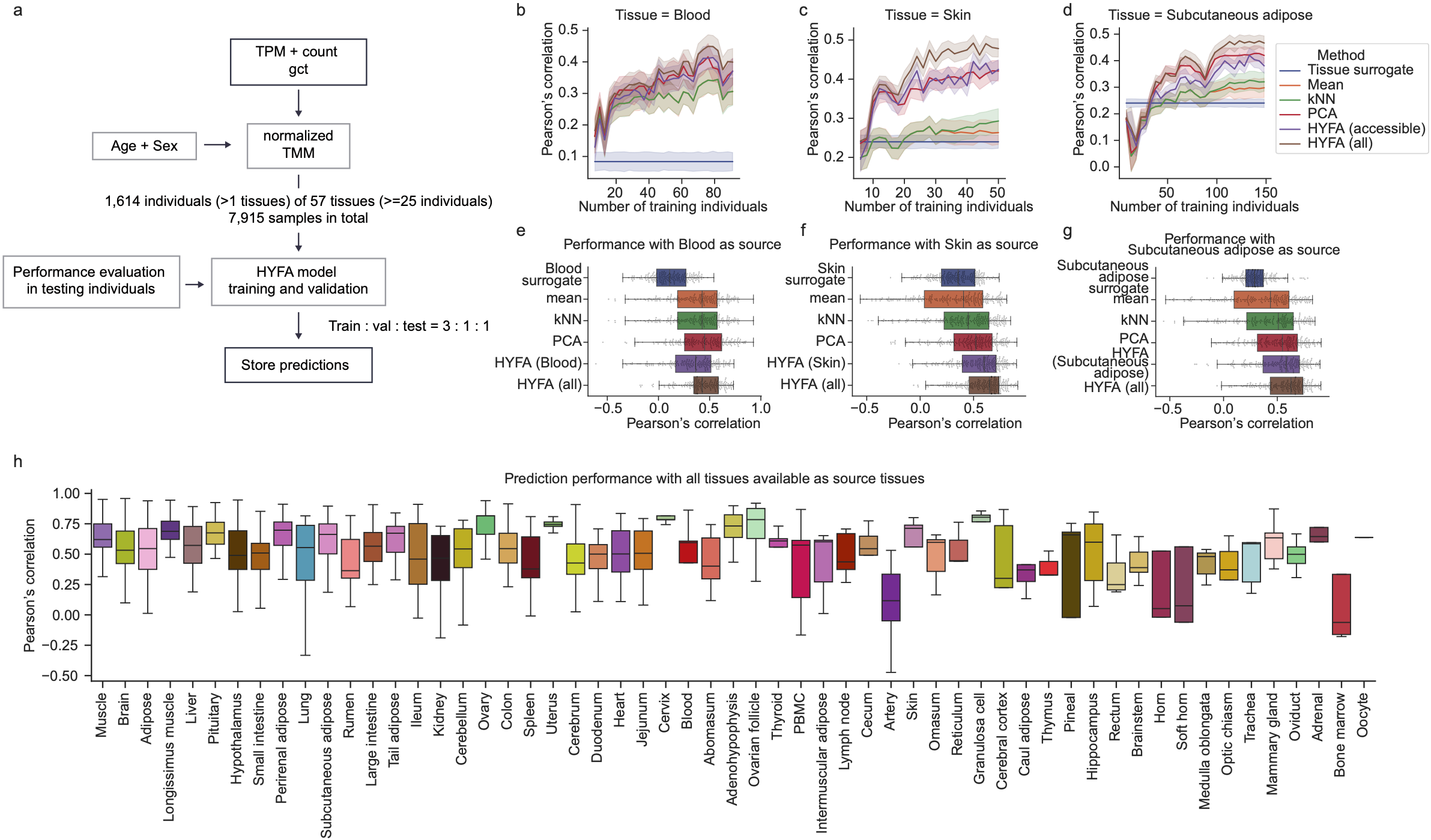

**Supplementary Fig. 20 | Gene expression imputation across 57 sheep tissues. a**, Overview of the gene expression imputation workflow using the HYFA (hypergraph factorization) approach. **b–d**, Imputation performance (measured as Pearson’s correlation between imputed and observed expression across genes) as a function of training sample size for different imputation methods, using blood (**b**), skin (**c**), and subcutaneous adipose tissue (**d**) as source tissues. “Tissue surrogate” uses gene expression information from a surrogate tissue as a proxy for the target tissue. “Mean” imputes missing values with the average expression of each gene. “kNN” uses k-nearest neighbors to estimate missing values. “PCA” projects reference gene expression into a lower-dimensional space using principal component analysis (30 components), followed by a linear regression model to predict target values. “HYFA (accessible)” uses gene expression data from accessible tissues (blood, skin, and subcutaneous adipose) from the same individual. “HYFA (all)” incorporates data from all available tissues of the individual. **e–g**, Performance comparison across imputation methods using blood (**e**), skin (**f**), and subcutaneous adipose (**g**) as source tissues. **h**, Prediction performance of HYFA with all tissues available as source tissues.

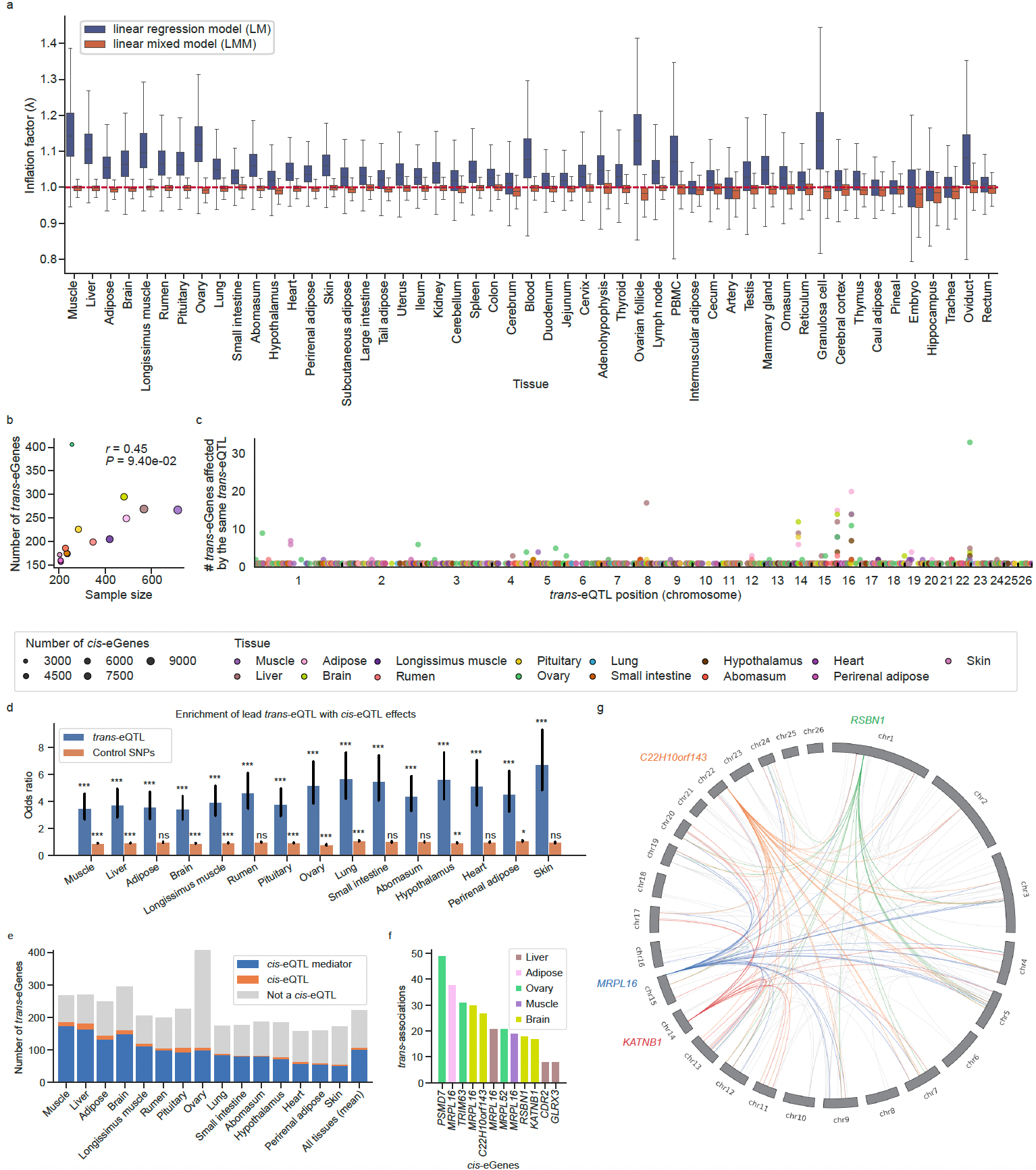

**Supplementary Fig. 21 | *trans*-eQTL mapping. a**, Comparison of genomic control inflation factor (λ) between linear regression models (LM) and linear mixed models (LMM) across tissues. **b**, Number of *trans*-eGenes identified as a function of tissue sample size. **c**, Genome-wide distribution of trans-eQTL hotspots, defined as loci associated with multiple *trans*-eGenes. **d**, Enrichment of lead *trans*-eQTL among *cis*-eQTL effects. Control SNPs were matched to lead *trans*-eQTL by minor allele frequency (MAF) and linkage disequilibrium (LD). ns: non-significant; *: *P* < 0.05; **: *P* < 0.01; ***: *P* < 0.001. **e**, Proportion of lead variants for *trans*-eGenes that are either significant *cis*-eQTL or mediated by *cis*-eQTL. **f**, *cis*-mediating eGenes with more than five colocalizing *trans*-eQTL. *trans*-associations were identified via colocalization (PP_H4_ > 0.8). **g**, Representative examples of *trans*-associations for four *cis*-eGenes with more than five colocalizing *trans*-eQTL in brain.

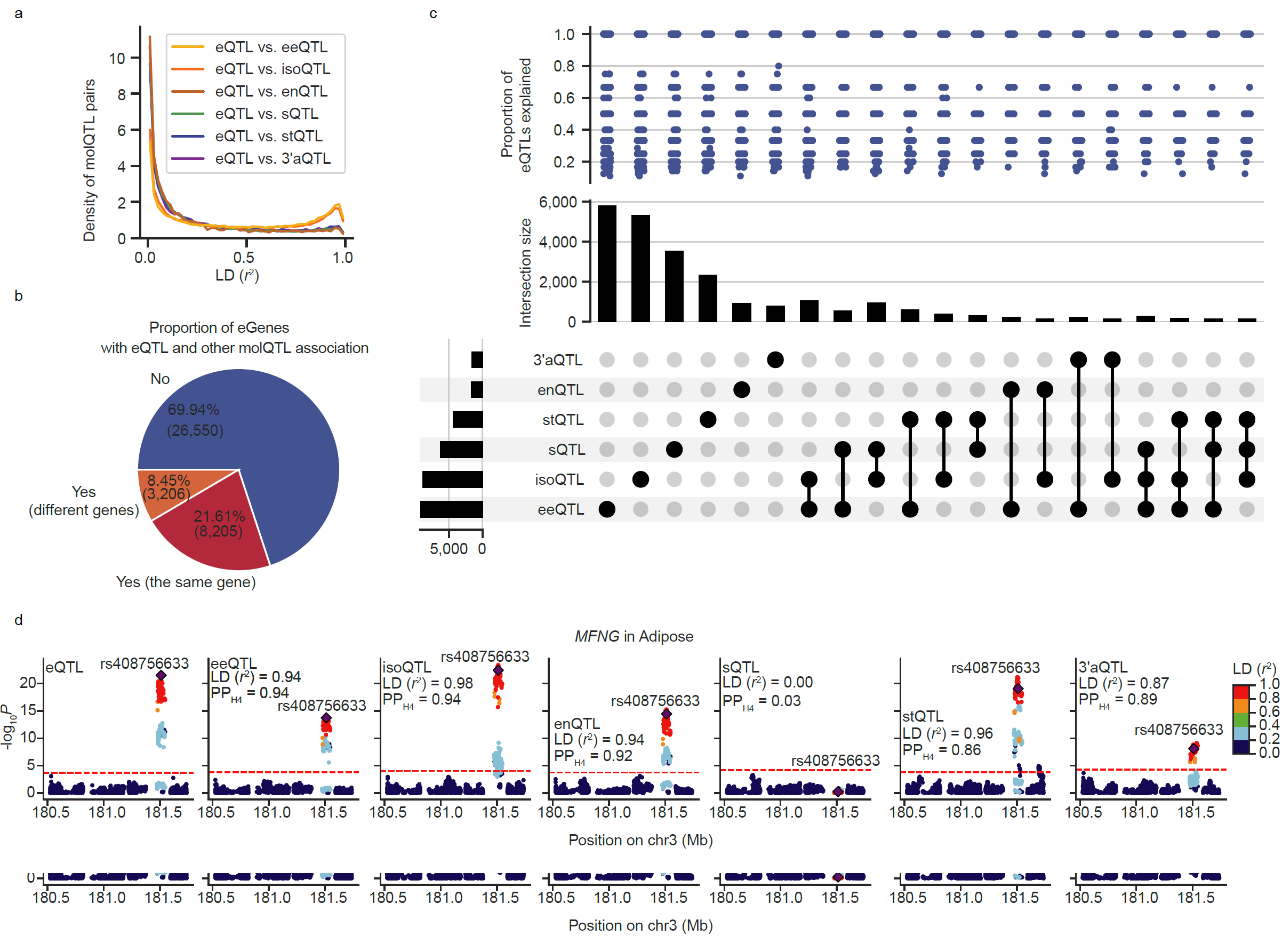

**Supplementary Fig. 22 | Shared and unique regulatory mechanisms across seven molQTL types. a**, Distribution of linkage disequilibrium (LD, measured as *r*^2^) between lead eQTL and lead variants of other molQTL types. **b**, Pie chart showing the proportion of eGenes with pleiotropic associations between eQTL and additional molQTL within the *cis*-window, as estimated using OPERA (v1.18)^66^, where pleiotropy was defined as shared genetic signals (PPA ≥ 0.9) across molQTL types. Associations are further stratified by whether the implicated molQTL affect the same gene or distinct genes. **c**, Upset plot illustrating the proportion of eQTL explained by each combination of pleiotropic associations across multiple molecular phenotypes. **d**, Example of a pleiotropic eQTL with regulatory effects on five additional molecular phenotypes within the same gene.

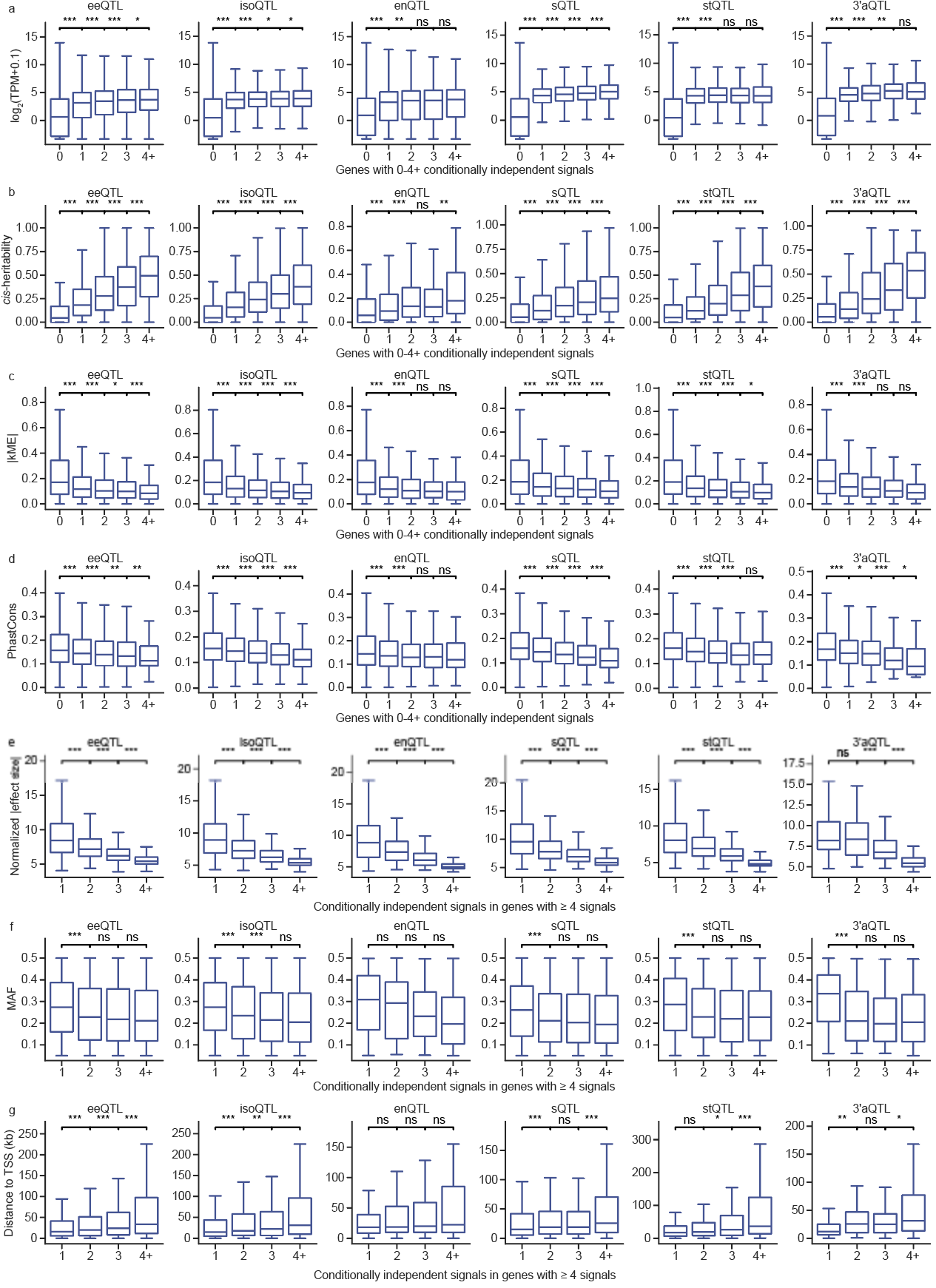

**Supplementary Fig. 23 | Characteristics of primary and non-primary molQTL. a–d**, Boxplots comparing gene expression levels (**a**), *cis*-heritability (**b**), gene co-expression network connectivity (|kME|) (**c**), and sequence conservation (PhastCons score) (**d**) for genes grouped by the number of independent *cis*-molQTL signals detected. **e-g**, Boxplots showing normalized |effect sizes| (**e**), minor allele frequency (MAF) (**f**), and distances to the transcriptional start site (TSS) (**g**) for molQTL signals ranked by order of discovery per molGene. *P* values were calculated using the two-sided Mann-Whitney U test. ns: non-significant; *: *P* < 0.05; **: *P* < 0.01; ***: *P* < 0.001.

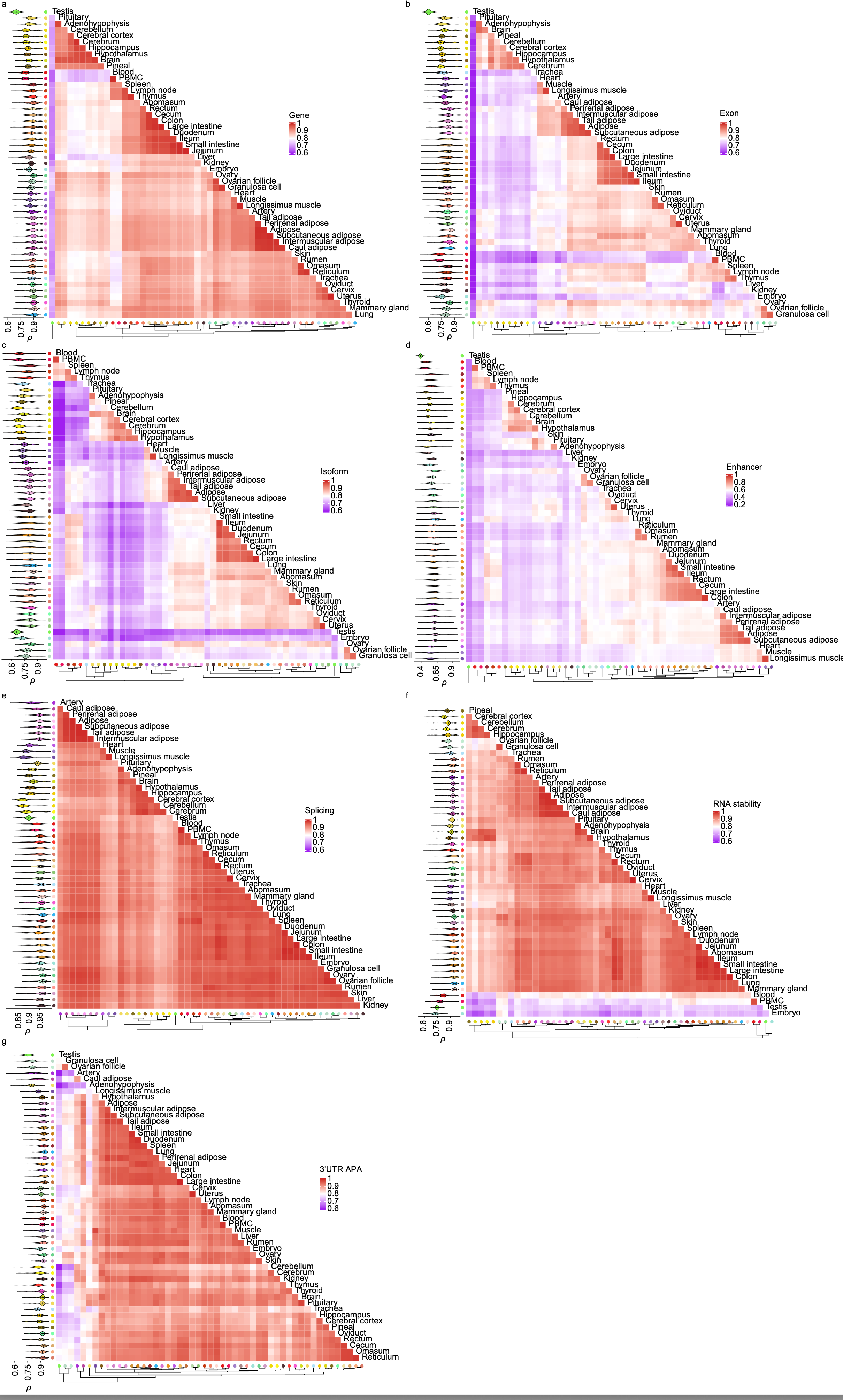

**Supplementary Fig. 24 | Tissue-sharing and specificity of molecular phenotypes. a–g**, Clustered heatmap showing pairwise Spearman’s correlation (*ρ*) of seven molecular phenotypes across tissues. Tissues were hierarchically clustered using the complete linkage method based on the maximum distance. Violin plots (left) display the distribution of Spearman’s *ρ* between each target tissue and all other tissues.

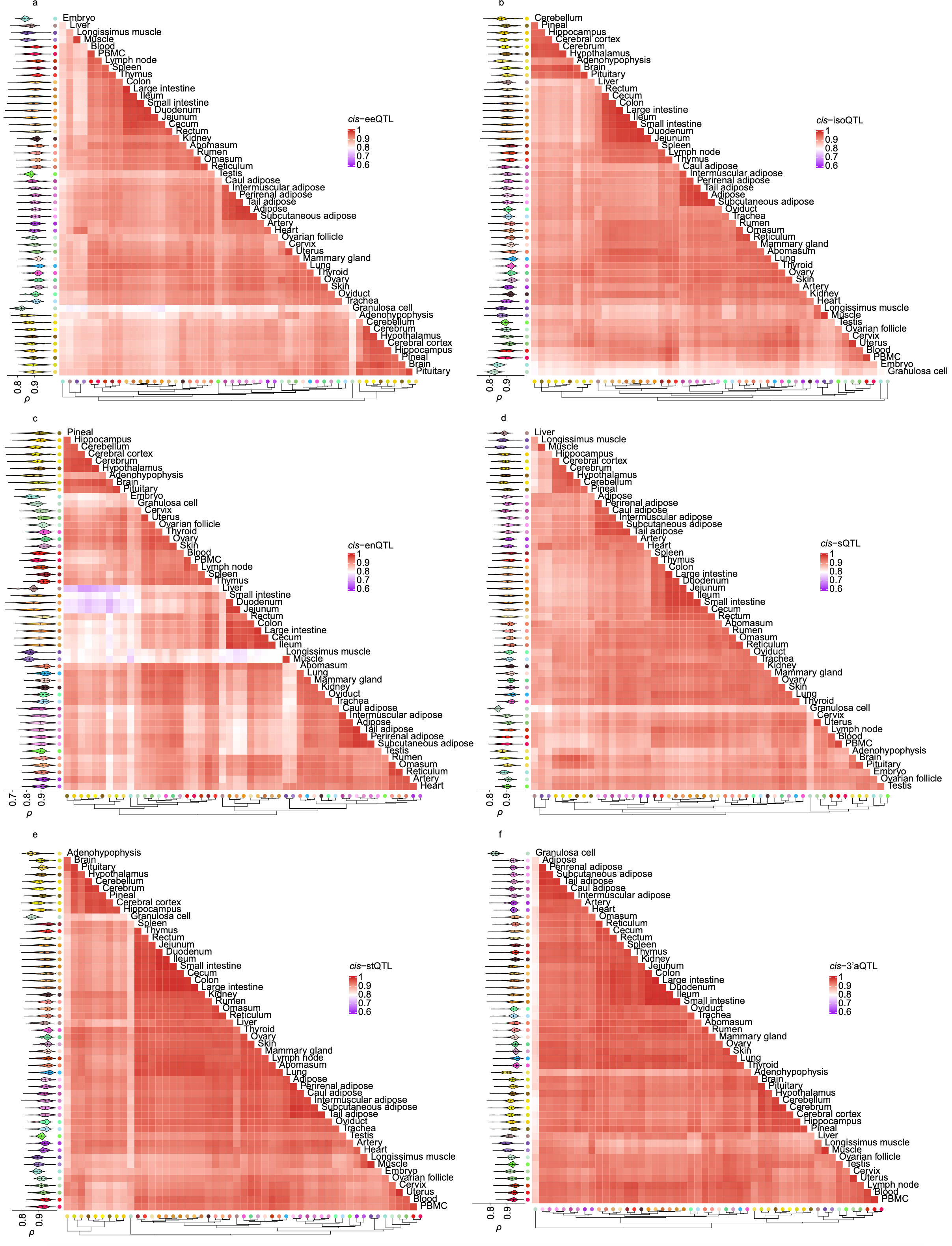
**Supplementary Fig. 25 | Tissue-sharing and specificity of molQTL. a–f**, Clustered heatmaps showing pairwise Spearman’s correlation (*ρ*) of *cis*-molQTL effect sizes across tissues for six molecular phenotypes other than *cis*-eQTL. Tissues were hierarchically clustered using the complete linkage method based on the maximum distance. Violin plots (left) display the distribution of Spearman’s *ρ* between each target tissue and all other tissues.

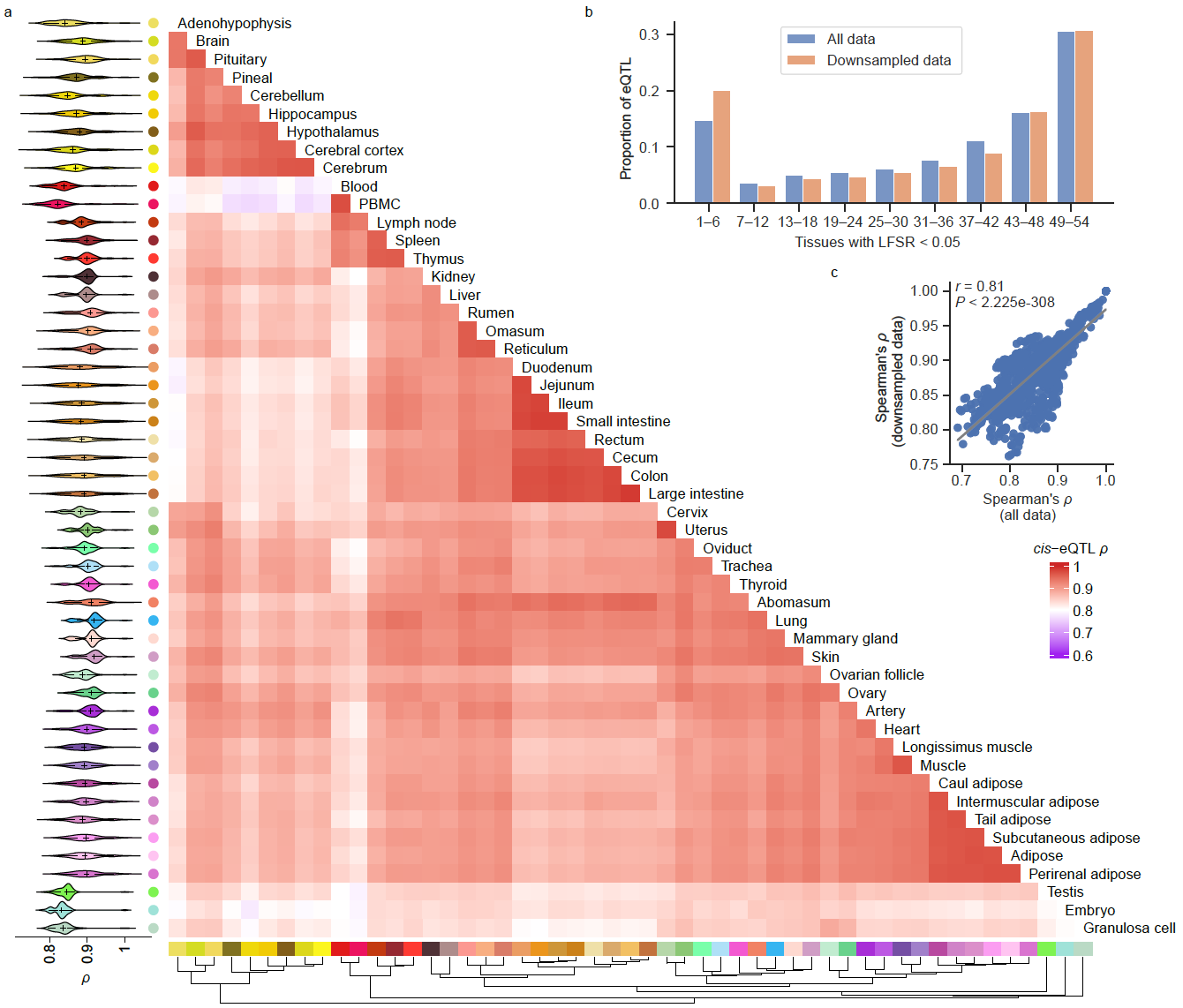

**Supplementary Fig. 26 | Tissue-sharing pattern of molQTL based on the down-sampled dataset. a**, Heatmap showing pairwise Spearman’s correlation (*ρ*) of *cis*-eQTL effect sizes across 51 tissues in the down-sampled dataset. Tissues were hierarchically clustered using the complete linkage method based on the maximum distance of *ρ*. Violin plots (left) display the distribution of Spearman’s *ρ* between each target tissue and all other tissues. **b**, Comparison of the proportion of eQTL active across tissues between the full and the down-sampled dataset, measured as the number of tissues with a *mashr* local false sign rate (LFSR; equivalent to FDR) < 0.05. **c**, Pearson’s correlation between Spearman’s *ρ* values of *cis*-eQTL effect sizes for the same pairwise tissue comparisons derived from the full and the down-sampled dataset.

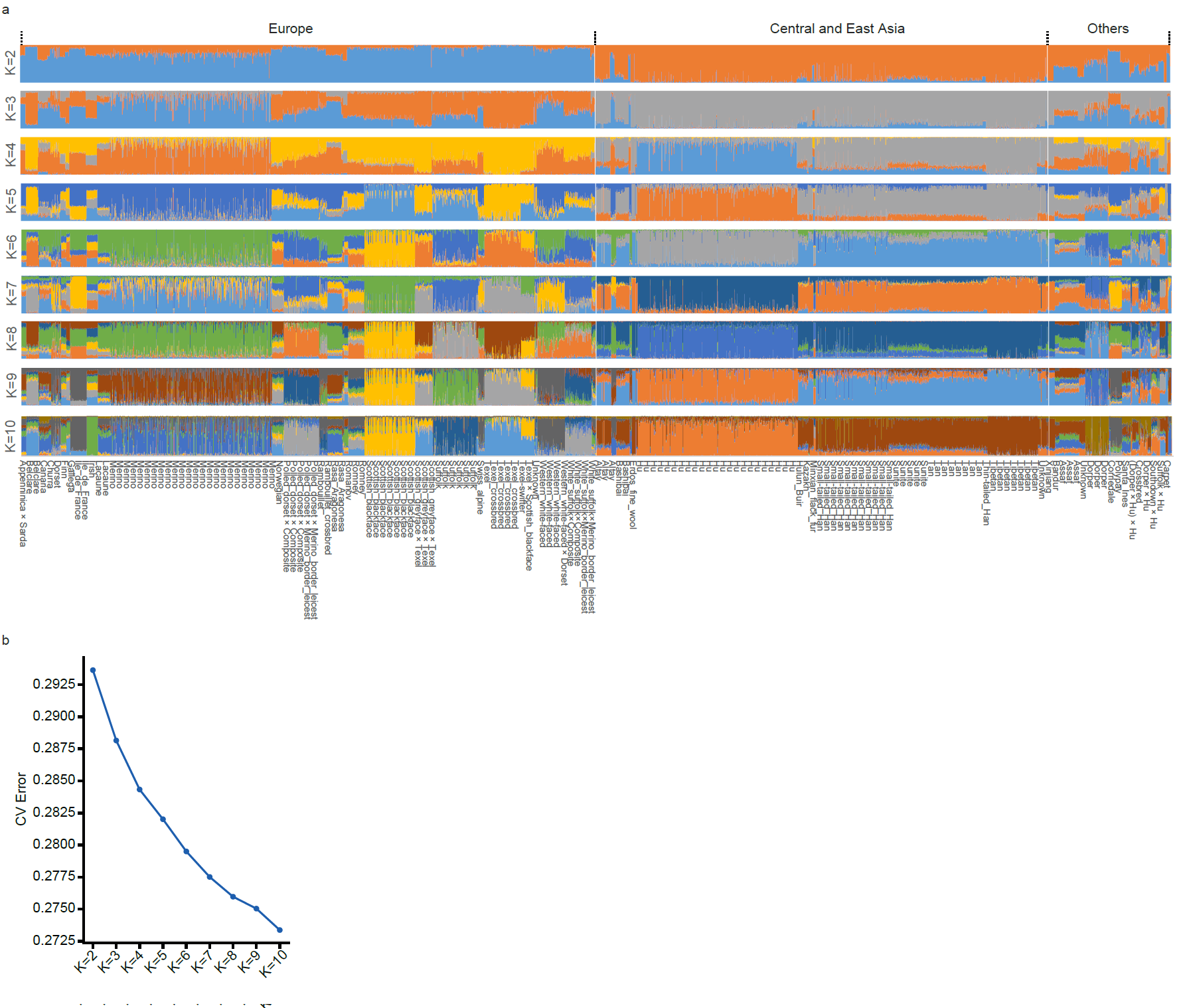

**Supplementary Fig. 27 | Global ancestry of 2,816 RNA-seq individuals. a**, Global ancestry profiles estimated using ADMIXTURE (v1.3.0)^67^, considering ancestry components from K = 2 to K = 10. **b**, Cross-validation (CV) error across different K values in the ADMIXTURE analysis.

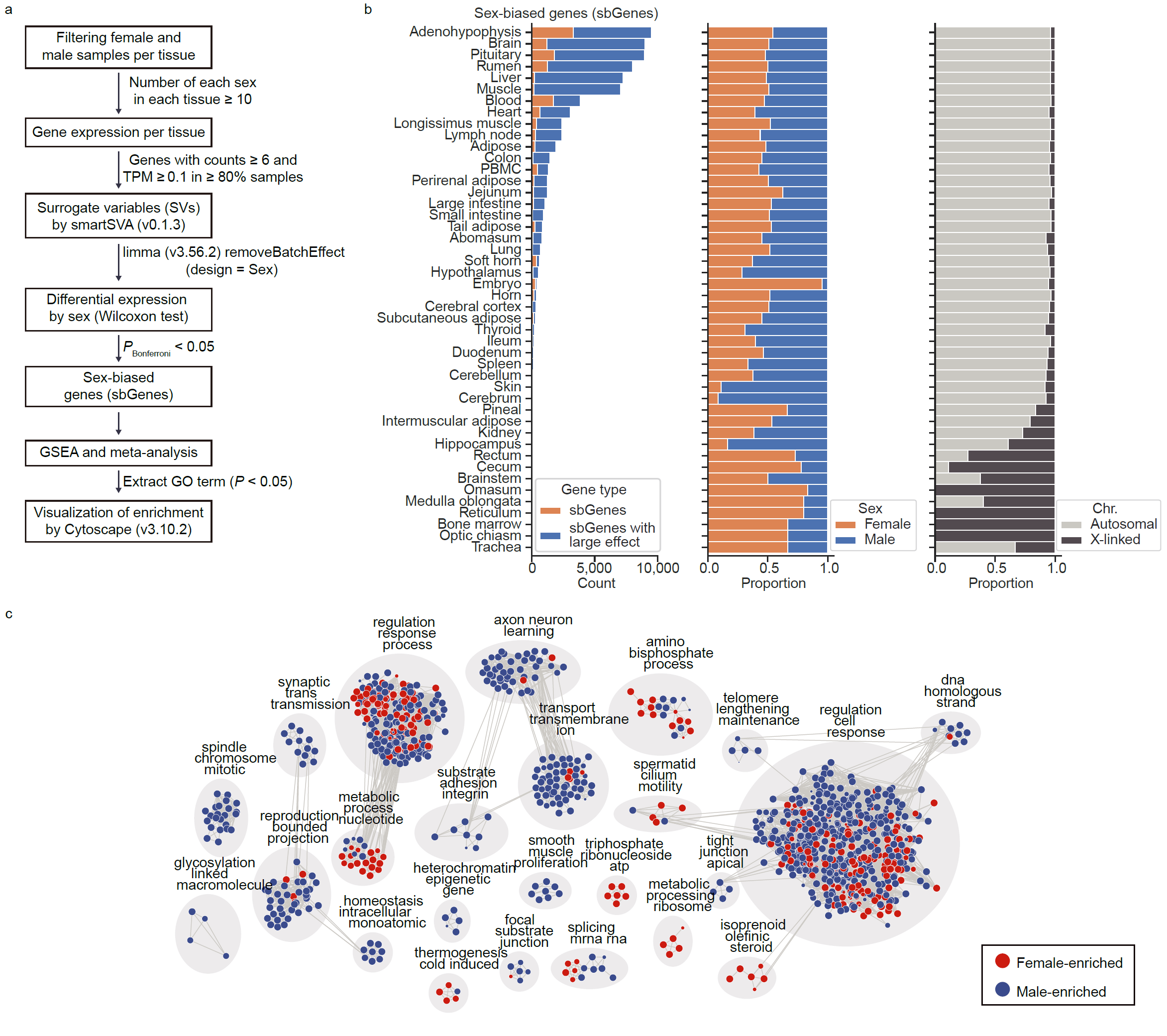

**Supplementary Fig. 28 | Sex-biased genes. a**, Workflow for the identification of sex-biased gene (sbGene) expression across 46 tissues. **b**, Discovery of sbGenes across 46 tissues (histogram) and their characteristics (stacked bar plots). Stacked bars indicate the proportions of female- and male-biased genes (Sex), as well as the distribution of sbGenes across autosomes and the X chromosome. **c**, Gene set enrichment clustering for sbGenes across tissues. Each grey circle represents a gene set enriched for either female- or male-biased expression, with circle size reflecting the *P*-value from a meta-analysis of GSEA results. Faint connecting lines indicate the degree of overlap (shared leading-edge genes) between gene sets.

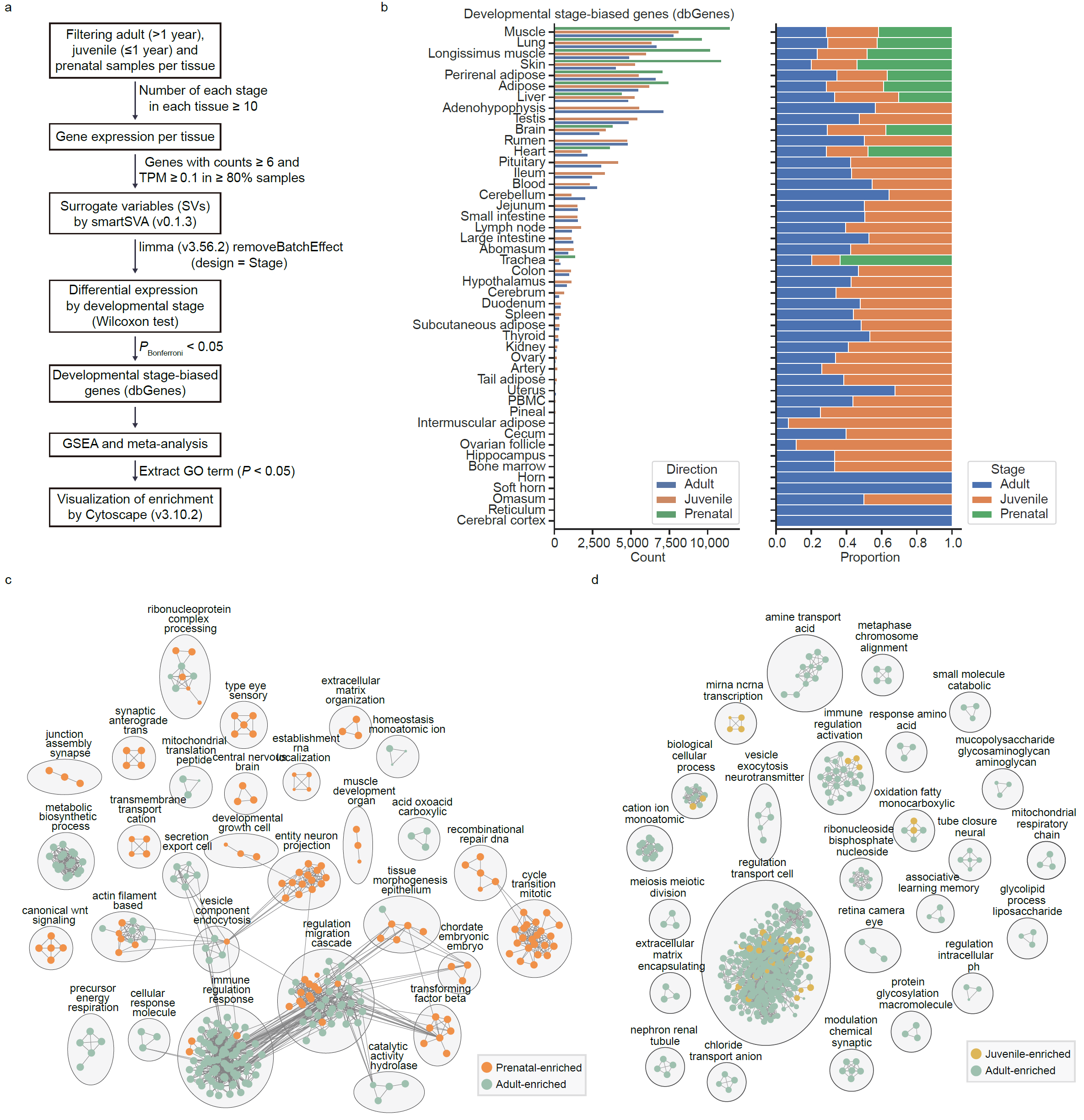

**Supplementary Fig. 29 | Development stage-biased genes. a**, Workflow for identifying developmental stage-biased gene (dbGene) expression across 51 tissues. **b**, Numbers (histogram) and proportions (stacked bar plots) of dbGenes across tissues. **c–d**, Gene set enrichment clustering of dbGenes across tissues. Each circle represents a gene set enriched for genes preferentially expressed at one of the three developmental stages, with circle size reflecting the *P*-value from a meta-analysis of GSEA results. Faint connecting lines indicate shared leading-edge genes between gene sets.

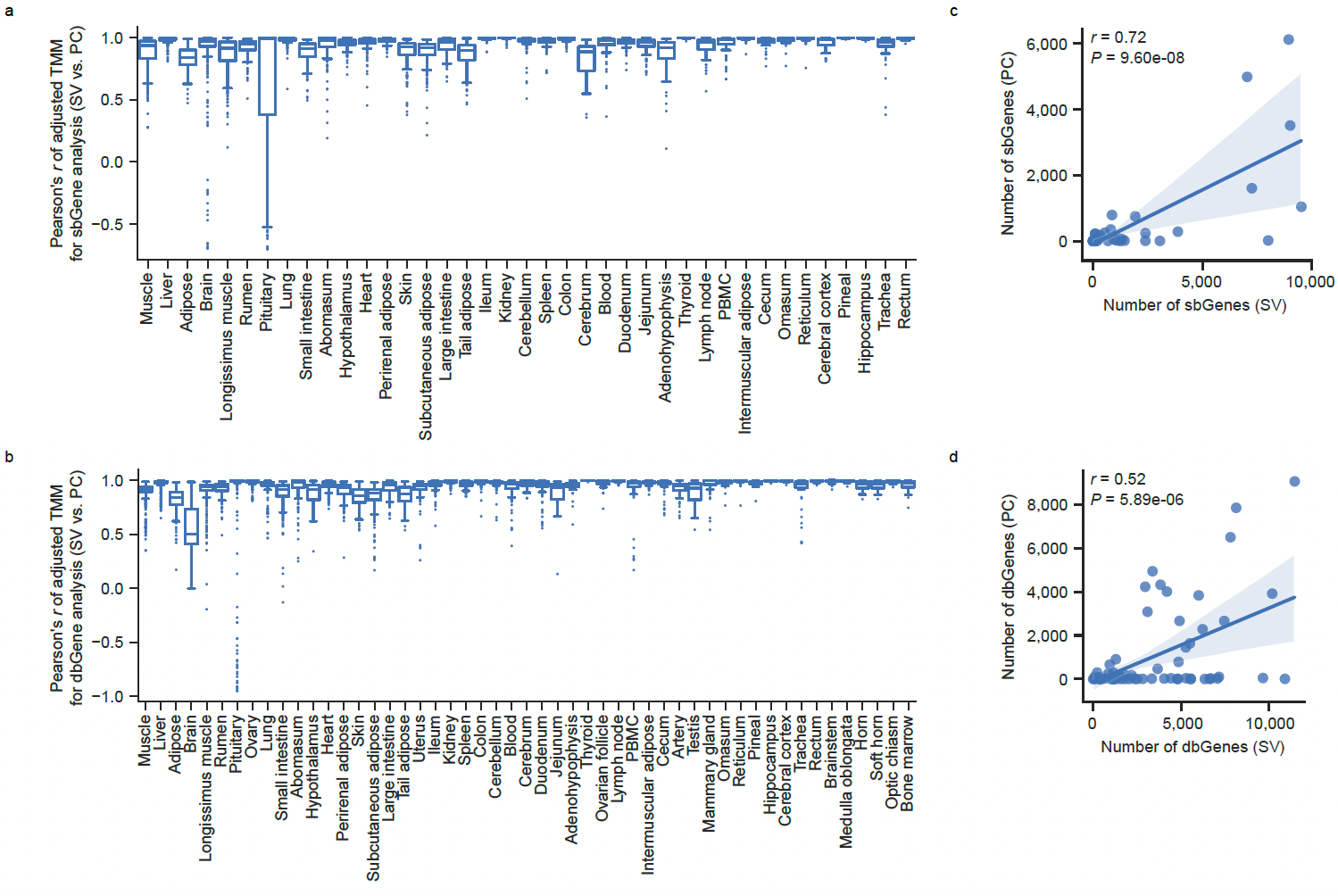

**Supplementary Fig. 30 | Impact of surrogate variable- and principal component-based covariate adjustment on biased gene expression analyses. a**, Pearson’s correlations between TMM-normalized expression matrices adjusted using surrogate variables (SVs) versus expression principal components (PCs) for sex-biased gene (sbGene) analyses across 46 tissues. **b**, Pearson’s correlations between TMM-normalized expression matrices adjusted using SVs versus PCs for developmental stage-biased gene (dbGene) analyses across 51 tissues. **c**, Comparison of the numbers of sbGenes identified using SV-adjusted and PC-adjusted strategies. **d**, Comparison of the numbers of dbGenes identified using SV-adjusted and PC-adjusted strategies.

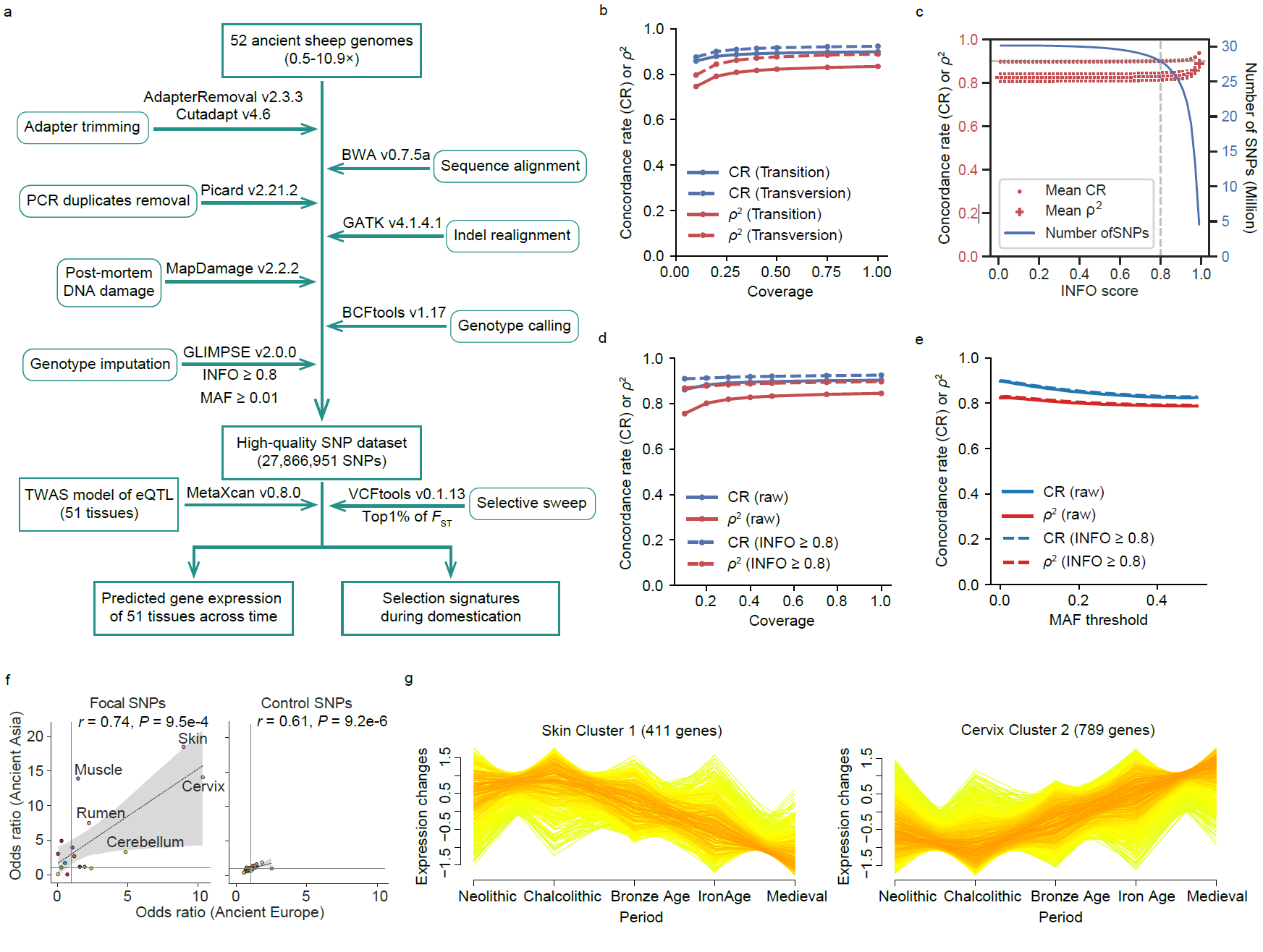

**Supplementary Fig. 31 | Ancient DNA analyses. a**, Workflow of ancient genome analysis. **b**, Comparison of imputation accuracy between transitions and transversions. **c**, Effect of INFO score thresholds on imputation accuracy (left) and retained SNP numbers on autosomes (right). **d**, Imputation accuracy across sequencing depths before and after INFO filtering. **e**, Imputation accuracy across minor allele frequency (MAF) bins before and after INFO filtering. **f**, Comparison of enrichment (odds ratio) of tissue-specific eQTL in genomic regions under strong selection in ancient European or Asian sheep populations. Ordinary least squares (OLS) regression indicates significant Pearson’s correlations (two-sided Student’s *t*-test). Focal *cis*-eQTL SNPs are shown on the left, with control SNPs matched for minor allele frequency (MAF) and linkage disequilibrium (LD) score on the right. **g**, Temporal trajectories of genetically predicted gene expression in ancient European sheep, showing decreased expression in skin (left) and increased expression in cervix (right).

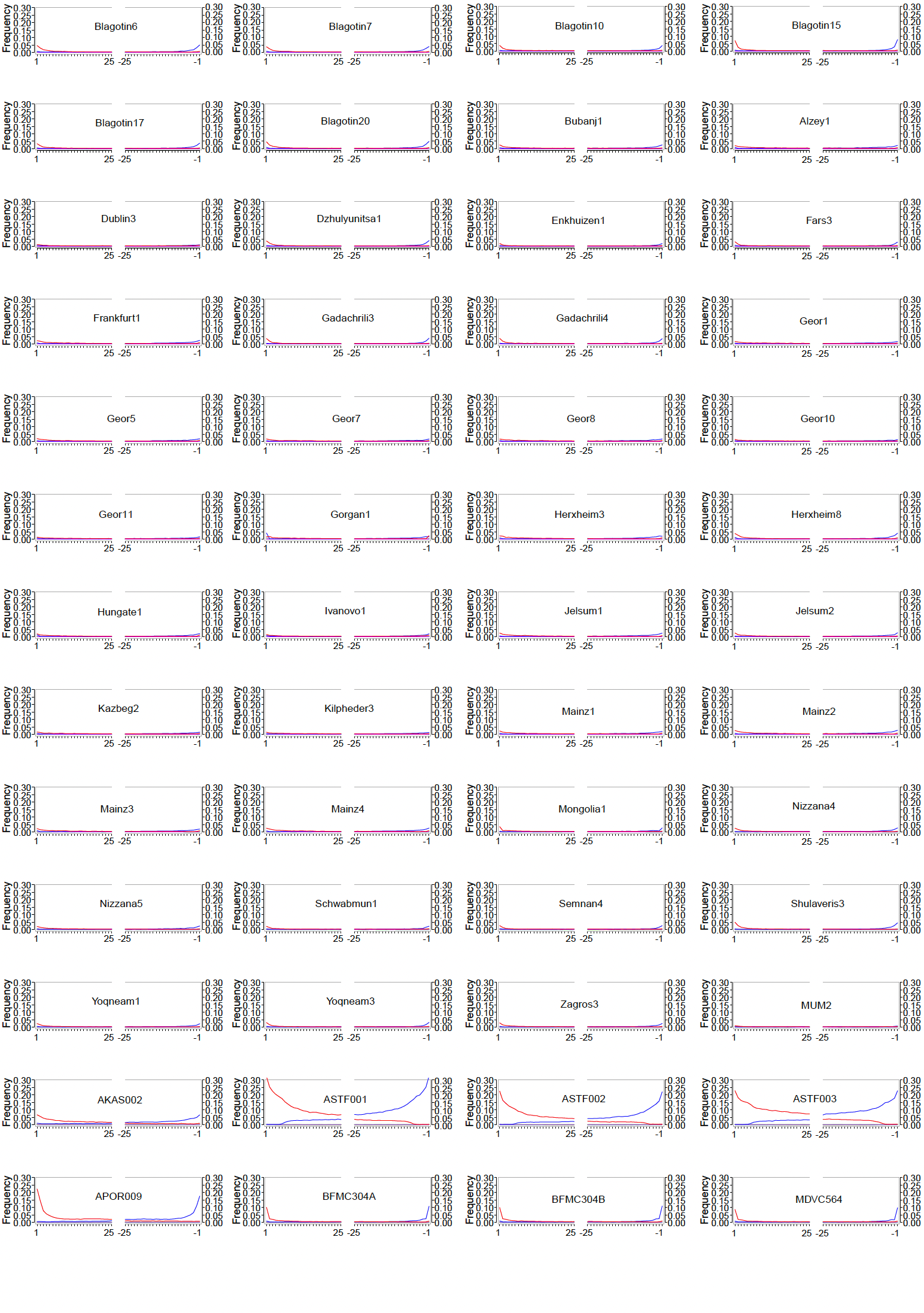

**Supplementary Fig. 32 | Nucleotide misincorporation profiles at the 5’ and 3’ termini of sequencing reads from 52 publicly available ancient genomes.** Misincorporation frequencies are shown across the first 25 bases from each read end. Red lines denote C→T substitutions and blue lines denote G→A substitutions.

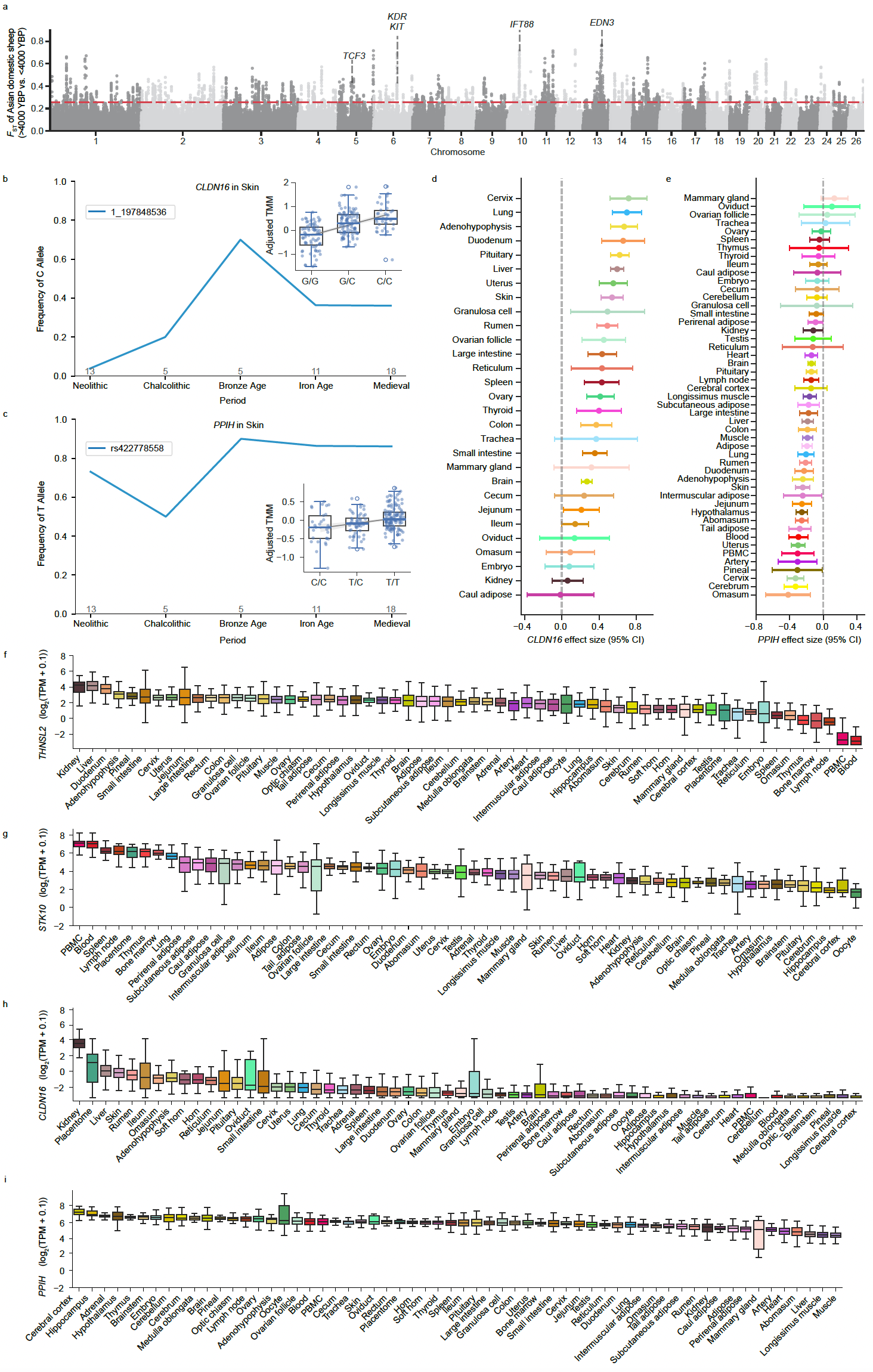

**Supplementary Fig. 33 | Spatiotemporal genomic differentiation and tissue-specific regulatory evolution in ancient domestic sheep. a**, Manhattan plot showing *F*_ST_ values between Asian domestic sheep dated to > 4,000 and < 4,000 years before present (yBP). The dashed red line indicates the top 1% threshold. **b–c**, Temporal allele frequency trajectories of SNPs 1_197848536 (C allele in *CLDN16*) and rs422778558 (T allele in *PPIH*) in ancient sheep. Line plots depict frequency changes over time; accompanying boxplots show genotype-dependent expression of each gene in modern skin samples. **d–e**, eQTL effect sizes (slope estimates) of 1_197848536 and rs422778558 on *CLDN16* and *PPIH* expression, respectively, across tissues. Error bars indicate 95% confidence intervals. **f-i**, Expression profiles of *THNSL2*, *STK10*, *CLDN16*, and *PPIH* across modern tissues, shown as log_2_(TPM + 0.1).

**Supplementary References**

1.	Hoffman, G.E. & Schadt, E.E. variancePartition: interpreting drivers of variation in complex gene expression studies. *BMC Bioinformatics* **17**, 483 (2016).

2.	Yu, G. Thirteen years of clusterProfiler. *Innovation (Camb)* **5**, 100722 (2024).

3.	Zhou, H.J., Li, L., Li, Y., Li, W. & Li, J.J. PCA outperforms popular hidden variable inference methods for molecular QTL mapping. *Genome Biology* **23**, 210 (2022).

4.	Ritchie, M.E. *et al.* limma powers differential expression analyses for RNA-sequencing and microarray studies. *Nucleic Acids Research* **43**, e47-e47 (2015).

5.	Antonio P. Camargo, Adrielle A. Vasconcelos, Mateus B. Fiamenghi, Gonçalo A. G. Pereira & Marcelo F. Carazzolle. tspex: a tissue-specificity calculator for gene expression data. *Research Square* (2020).

6.	Li, Y., Ge, X., Peng, F., Li, W. & Li, J.J. Exaggerated false positives by popular differential expression methods when analyzing human population samples. *Genome Biology* **23**, 79 (2022).

7.	Lenffer, J. *et al.* OMIA (Online Mendelian Inheritance in Animals): an enhanced platform and integration into the Entrez search interface at NCBI. *Nucleic Acids Res* **34**, D599-601 (2006).

8.	Langfelder, P. & Horvath, S. WGCNA: an R package for weighted correlation network analysis. *BMC Bioinformatics* **9**, 559 (2008).

9.	Durinck, S., Spellman, P.T., Birney, E. & Huber, W. Mapping identifiers for the integration of genomic datasets with the R/Bioconductor package biomaRt. *Nat Protoc* **4**, 1184-91 (2009).

10.	Bastian, M., Heymann, S. & Jacomy, M. Gephi: An Open Source Software for Exploring and Manipulating Networks. *Proceedings of the International AAAI Conference on Web and Social Media* **3**, 361-362 (2009).

11.	Zhang, Q., Liu, H. & Bu, F. High performance of a GPU-accelerated variant calling tool in genome data analysis. *bioRxiv*, 2021.12.12.472266 (2021).

12.	Chen, S., Zhou, Y., Chen, Y. & Gu, J. fastp: an ultra-fast all-in-one FASTQ preprocessor. *Bioinformatics* **34**, i884-i890 (2018).

13.	Li, H. Aligning sequence reads, clone sequences and assembly contigs with BWA-MEM. *arXiv* (2013).

14.	Danecek, P. *et al.* Twelve years of SAMtools and BCFtools. *GigaScience* **10**(2021).

15.	McKenna, A. *et al.* The Genome Analysis Toolkit: a MapReduce framework for analyzing next-generation DNA sequencing data. *Genome Research* **20**, 1297-303 (2010).

16.	Rubinacci, S., Hofmeister, R.J., Sousa da Mota, B. & Delaneau, O. Imputation of low-coverage sequencing data from 150,119 UK Biobank genomes. *Nature Genetics* **55**, 1088-1090 (2023).

17.	Purcell, S. *et al.* PLINK: a tool set for whole-genome association and population-based linkage analyses. *American Journal of Human Genetics* **81**, 559-75 (2007).

18.	Teng, J. *et al.* A compendium of genetic regulatory effects across pig tissues. *Nature Genetics* (2024).

19.	Guan, D. *et al.* Genetic regulation of gene expression across multiple tissues in chickens. *Nature Genetics* (2025).

20.	Liu, S. *et al.* A multi-tissue atlas of regulatory variants in cattle. *Nature Genetics* (2022).

21.	Browning, B.L., Tian, X., Zhou, Y. & Browning, S.R. Fast two-stage phasing of large-scale sequence data. *Am J Hum Genet* **108**, 1880-1890 (2021).

22.	Browning, B.L., Zhou, Y. & Browning, S.R. A One-Penny Imputed Genome from Next-Generation Reference Panels. *The American Journal of Human Genetics* **103**, 338-348 (2018).

23.	Wang, Q. *et al.* Weighted gene co-expression network analysis reveals genes related to growth performance in Hu sheep. *Scientific Reports* **14**, 13043 (2024).

24.	Cai, Y. *et al.* Ancient genomes reveal the evolutionary history and origin of cashmere producing goats in China. *Molecular Biology and Evolution* **37**, 2099-2109 (2020).

25.	Li, R. *et al.* A Hu sheep genome with the first ovine Y chromosome reveal introgression history after sheep domestication. *SCIENCE CHINA Life Sciences* (2020).

26.	Hadish, J.A., Honaas, L.A. & Ficklin, S.P. Predicting Phenotypic Traits Using a Massive RNA-seq Dataset. 2023.12.05.570195 (2023).

27.	Zhao, C. *et al.* Breed identification using breed-informative SNPs and machine learning based on whole genome sequence data and SNP chip data. *J Anim Sci Biotechnol* **14**, 85 (2023).

28.	Zhi, Y. *et al.* Advanced molecular system for accurate identification of chicken genetic resources. *Computers and Electronics in Agriculture* **231**, 109989 (2025).

29.	Yang, J., Lee, S.H., Goddard, M.E. & Visscher, P.M. GCTA: a tool for genome-wide complex trait analysis. *Am J Hum Genet* **88**, 76-82 (2011).

30.	The GTEx Consortium atlas of genetic regulatory effects across human tissues. *Science* **369**, 1318-1330 (2020).

31.	Stegle, O., Parts, L., Piipari, M., Winn, J. & Durbin, R. Using probabilistic estimation of expression residuals (PEER) to obtain increased power and interpretability of gene expression analyses. *Nat Protoc* **7**, 500-7 (2012).

32.	Stegle, O., Parts, L., Durbin, R. & Winn, J. A Bayesian framework to account for complex non-genetic factors in gene expression levels greatly increases power in eQTL studies. *PLoS Comput Biol* **6**, e1000770 (2010).

33.	Teng, J. *et al.* OmiGA for ultra-efficient molecular quantitative trait loci mapping. *Nature Communications* (2026).

34.	Gilmour, A.R., Thompson, R. & Cullis, B.R. Average information REML: an efficient algorithm for variance parameter estimation in linear mixed models. *Biometrics*, 1440-1450 (1995).

35.	Taylor-Weiner, A. *et al.* Scaling computational genomics to millions of individuals with GPUs. *Genome Biology* **20**, 228 (2019).

36.	Storey, J.D. & Tibshirani, R. Statistical significance for genomewide studies. *Proceedings of the National Academy of Sciences* **100**, 9440-9445 (2003).

37.	The Genotype-Tissue Expression (GTEx) pilot analysis: multitissue gene regulation in humans. *Science* **348**, 648-60 (2015).

38.	Mohammadi, P., Castel, S.E., Brown, A.A. & Lappalainen, T. Quantifying the regulatory effect size of cis-acting genetic variation using allelic fold change. *Genome Res* **27**, 1872-1884 (2017).

39.	Castel, S.E., Mohammadi, P., Chung, W.K., Shen, Y. & Lappalainen, T. Rare variant phasing and haplotypic expression from RNA sequencing with phASER. *Nature Communications* **7**, 12817 (2016).

40.	Pockrandt, C., Alzamel, M., Iliopoulos, C.S. & Reinert, K. GenMap: ultra-fast computation of genome mappability. *Bioinformatics* **36**, 3687-3692 (2020).

41.	Neph, S. *et al.* BEDOPS: high-performance genomic feature operations. *Bioinformatics* **28**, 1919-1920 (2012).

42.	Navarro Gonzalez, J. *et al.* The UCSC Genome Browser database: 2021 update. *Nucleic Acids Research* **49**, D1046-d1057 (2021).

43.	Cingolani, P. *et al.* A program for annotating and predicting the effects of single nucleotide polymorphisms, SnpEff: SNPs in the genome of Drosophila melanogaster strain w1118; iso-2; iso-3. *Fly (Austin)* **6**, 80-92 (2012).

44.	Mostafavi, H., Spence, J.P., Naqvi, S. & Pritchard, J.K. Systematic differences in discovery of genetic effects on gene expression and complex traits. *Nature Genetics* **55**, 1866-1875 (2023).

45.	Viñas, R. *et al.* Hypergraph factorization for multi-tissue gene expression imputation. *Nature Machine Intelligence* **5**, 739-753 (2023).

46.	Saha, A. & Battle, A. False positives in trans-eQTL and co-expression analyses arising from RNA-sequencing alignment errors. *F1000Res* **7**, 1860 (2018).

47.	Giambartolomei, C. *et al.* Bayesian test for colocalisation between pairs of genetic association studies using summary statistics. *PLoS Genet* **10**, e1004383 (2014).

48.	Urbut, S.M., Wang, G., Carbonetto, P. & Stephens, M. Flexible statistical methods for estimating and testing effects in genomic studies with multiple conditions. *Nature Genetics* **51**, 187-195 (2019).

49.	Zou, Y., Carbonetto, P., Wang, G. & Stephens, M. Fine-mapping from summary data with the “Sum of Single Effects” model. *PLOS Genetics* **18**, e1010299 (2022).

50.	Wang, G., Sarkar, A., Carbonetto, P. & Stephens, M. A simple new approach to variable selection in regression, with application to genetic fine mapping. *J R Stat Soc Series B Stat Methodol* **82**, 1273-1300 (2020).

51.	Oliva, M. *et al.* The impact of sex on gene expression across human tissues. *Science* **369**(2020).

52.	Chen, J. *et al.* Fast and robust adjustment of cell mixtures in epigenome-wide association studies with SmartSVA. *BMC Genomics* **18**, 413 (2017).

53.	Shannon, P. *et al.* Cytoscape: a software environment for integrated models of biomolecular interaction networks. *Genome Res* **13**, 2498-504 (2003).

54.	Daly, K.G. *et al.* Ancient genomics and the origin, dispersal, and development of domestic sheep. *Science* **387**, 492-497 (2025).

55.	Davenport, K.M. *et al.* An improved ovine reference genome assembly to facilitate in-depth functional annotation of the sheep genome. *Gigascience* **11**(2022).

56.	Erven, J.A.M. *et al.* A High-Coverage Mesolithic Aurochs Genome and Effective Leveraging of Ancient Cattle Genomes Using Whole Genome Imputation. *Molecular Biology and Evolution* **41**(2024).

57.	Erven, J.A.M. *et al.* Inferring Domestic Goat Demographic History Through Ancient Genome Imputation. *Genome Biology and Evolution* **17**(2025).

58.	Kielbasa, S.M., Wan, R., Sato, K., Horton, P. & Frith, M.C. Adaptive seeds tame genomic sequence comparison. *Genome Research* **21**, 487-93 (2011).

59.	Fu, W. *et al.* RGD v2.0: a major update of the ruminant functional and evolutionary genomics database. *Nucleic Acids Research* (2021).

60.	Pedersen, B.S. & Quinlan, A.R. cyvcf2: fast, flexible variant analysis with Python. *Bioinformatics* **33**, 1867-1869 (2017).

61.	Harris, C.R. *et al.* Array programming with NumPy. *Nature* **585**, 357-362 (2020).

62.	Keightley, P.D. & Jackson, B.C. Inferring the Probability of the Derived vs. the Ancestral Allelic State at a Polymorphic Site. *Genetics* **209**, 897-906 (2018).

63.	Dobin, A. *et al.* STAR: ultrafast universal RNA-seq aligner. *Bioinformatics* **29**, 15-21 (2013).

64.	Pertea, M., Kim, D., Pertea, G.M., Leek, J.T. & Salzberg, S.L. Transcript-level expression analysis of RNA-seq experiments with HISAT, StringTie and Ballgown. *Nat Protoc* **11**, 1650-67 (2016).

65.	Liao, Y., Smyth, G.K. & Shi, W. featureCounts: an efficient general purpose program for assigning sequence reads to genomic features. *Bioinformatics* **30**, 923-30 (2014).

66.	Wu, Y. *et al.* Joint analysis of GWAS and multi-omics QTL summary statistics reveals a large fraction of GWAS signals shared with molecular phenotypes. *Cell Genomics* **3**, 100344 (2023).

67.	Alexander, D.H., Novembre, J. & Lange, K. Fast model-based estimation of ancestry in unrelated individuals. *Genome Res* **19**, 1655-64 (2009).
